# Dissecting fluctuating selection: A unified population and quantitative genetics framework

**DOI:** 10.1101/2025.05.19.654983

**Authors:** Esdras Tuyishimire, Molly K. Burke, Elizabeth G. King

## Abstract

One of the longstanding debates in evolutionary biology is the effect of fluctuating selection on genetic changes in populations. However, the extent to which these periodic forces influence organisms at both genomic and phenotypic levels remains unclear. Despite the compelling evidence of fluctuating selection from recent studies, there is a disconnect between empirical and theoretical findings concerning the underlying mechanisms due to the limited evidence regarding the scale and processes that generate genome-wide oscillations. This study aims to elucidate how both genetic factors (e.g. heritability, number of causative loci) and ecological factors (e.g. season length, the difference in the phenotypic optima between seasons, population size dynamics) drive fluctuating selection and to identify the parameters that produce consistent oscillatory patterns. We developed a modeling framework integrating quantitative and population genetics to simulate a population under various selection regimes. We applied spectral analysis to detect periodicity, indicating cyclical selective environments. Our simulations highlight the conditions sustaining oscillations in allele frequencies over time. Spectral analysis successfully identifies the periodic patterns from allele frequency trajectories, even under highly complex selection regimes. Not only does our study clarify the conditions that yield oscillatory behaviors, but these parameters can also potentially be estimated in natural populations, providing a possibility of empirically testing these models.

**Significance statement:** As genomic data is becoming increasingly available for different species across time, one observation are patterns where alleles oscillate in a seasonal pattern, which has been interpreted as a signature of fluctuating selection. However, the field lacks theoretical models that predict these persistent oscillations in allele frequencies caused by fluctuating selection. We develop such a theoretical model, defining the conditions under which recurrent, strong oscillations in allele frequencies are predicted to occur because of fluctuating selection. In addition, we develop a novel method to detect patterns of fluctuating selection from genomic data using spectral analysis. Our model considers key parameters that are measurable in real populations, which is an added advantage to test them empirically. Our research paves the way for field biologists to test our predictions and brings us closer to reliably forecasting how populations will evolve in environments that are frequently changing.

## Introduction

Understanding the dynamic nature of evolution in populations is at the core of evolutionary biology. The direction and intensity of selection can vary significantly through time (Grant and Grant 2002; Bell 2010; Campbell-Staton et al. 2017; Endler 2020). Various factors can lead to these fluctuations, in some cases in a regular pattern, such as in the case of seasonal variability. In addition, as climate change imposes unprecedented alterations in conditions across the globe, fluctuations in selection pressure are predicted to intensify (Gienapp et al. 2014). Thus, the ability to predict how a population will evolve in response to dynamic environmental conditions has implications for the future of biodiversity. However, while there is widespread appreciation for the relevance of fluctuating selection for populations (Thomas E Reed et al. 2011; Bergland et al. 2014; Van Buskirk and Smith 2021; Pfenninger and Foucault 2022; Yair and Coop 2022; Bitter et al. 2024; Nunez et al. 2024), we still lack a clear understanding of the potential impact of fluctuating selection on genome-wide allele frequency dynamics, and what the genomic signature of fluctuating selection will be with different patterns of temporal variation in selection.

Many empirical studies have shown that strong selective pressures can produce rapid phenotypic and genomic changes across diverse species, a phenomenon that has been vigorously documented not only in domesticated plants and animals (Doebley et al. 2006; Trut et al. 2009) but also in wild populations exposed to well-defined selective pressures (Grant and Grant 2002; Campbell-Staton et al. 2017; Waldvogel et al. 2018) and through laboratory experiments (Burke et al. 2010; Tenaillon et al. 2016). Furthermore, the direction and intensity of selection have been shown to fluctuate over time (Grant and Grant 2002; Bell 2010; Gossmann et al. 2014; Campbell-Staton et al. 2017; Abdul-Rahman et al. 2021; Roberts Kingman et al. 2021; Kelly 2022) due to various factors, including seasonal temperature variability, boom and bust cycles of predator populations, and other biotic and abiotic agents (Louthan and Kay 2011; Waldvogel et al. 2018; McAdam et al. 2019; Van Buskirk and Smith 2021; Rusuwa et al. 2022). In natural populations, a recent observation is very rapid changes in allele frequencies, with some oscillating back and forth corresponding with the seasonal cycle, (e.g., (Behrman et al. 2018; Machado et al. 2021; Pfenninger and Foucault 2022; Rudman et al. 2022)), with the explanation for the pattern being fluctuating seasonal selection. For example, Behrman et al. 2018 suggested that seasonal changes in bacterial populations infecting fruit flies, *Drosophila melanogaster (D. melanogaster)* cause seasonal cycling of allele frequencies in immunity-related genes. In addition, Pfenninger & Foucault (2022), studied fluctuating environments using *Chironomus riparius* and found similar results with those reported from the previous studies of *D. melanogaster* (Behrman et al. 2018; Machado et al. 2021; Rudman et al. 2022; Bitter et al. 2024) about the genomic and phenotypic changes brought by the seasonal environment on natural populations.

Despite the many empirical examples identifying recurrently oscillating allele frequencies and attributing the cause to fluctuating selection, population genetic models that include fluctuating selection coefficients generally find only a narrow range of parameter space leading to stable oscillations in allele frequencies (Hoekstra et al. 1985; Wittmann et al. 2017; Bertram and Masel 2019). For instance, in their theoretical model, Wittmann et al. showed that seasonally fluctuating selection can maintain polymorphism at many loci when dominance reverses between seasons, particularly when the seasonally maladapted allele is partially or completely recessive (Wittmann et al. 2017). Consistently, another theoretical work showed that different mechanisms may maintain polymorphisms under weak versus strong fluctuating selection, including protection from selection, density-dependent effects coming from demographic change and genetic interactions (Bertram and Masel 2019) or mechanisms like dormancy (Glaser-Schmitt et al. 2021). Recently, Brud extended this framework by analyzing a broad range of dominance schemes and showed that the maintenance of polymorphism does not necessarily require reversal of dominance (Brud 2025). Furthermore, the experimental and theoretical works show that fluctuations in population size can generate large changes in allele frequency exceeding those expected from genetic drift; therefore, some allele frequency changes may also come from demographic noise instead of selection alone (Ascensao et al. 2024; Nunez et al. 2024). This disconnect between theory and empirical results has led to disagreement and uncertainty in whether the observed empirical patterns of oscillating allele frequencies are a genomic signature of fluctuating selection or an experimental artifact, or some other mechanism, such as short-term changes from recent selection and demographic alterations (Bertram and Masel 2019; Glaser-Schmitt et al. 2021; Lynch et al. 2024). Previous theoretical models of fluctuating selection have typically fallen broadly into two classes: 1) population genetic models that focus on predicting allele frequency trajectories, often focusing on a single or very few loci, and modeling fluctuating selection coefficients directly on loci; and 2) quantitative genetic models that focus on predicting phenotypic trajectories and typically assume a polygenic (often infinitesimal) genetic architecture defined by emergent genetic properties (i.e. heritabilities and genetic correlations) and often ignored the dynamics of individual loci. Past computational limitations of tracking loci genome-wide motivated the contrast in these two approaches, but while these limits have been diminishing rapidly over time, most models still fall largely into one of these two frameworks without fully bridging these approaches, a gap in the field that has been noted by others (e.g., Barghi *et al*. 2020). Many previous quantitative genetic models of fluctuating selection have produced robust predictions for resulting phenotypic evolution (Gillespie 1973; Gillespie 1977), the evolution of phenotypic plasticity and bet-hedging (Roff 1997), and the maintenance of genetic variation (Ellner and Hairston Jnr 1994; Bürger and Ghnelfarb 1999). While most previous population genetic approaches to fluctuating selection focused on allele frequency dynamics have found only very limited conditions that will lead to stable oscillations in allele frequencies, recently, some models have considered more complex scenarios to define the parameters under which allele frequency oscillations are possible (Wittmann et al. 2017; Bertram and Masel 2019; Glaser-Schmitt et al. 2021; Kim 2023). An important detail of most population genetic approaches is that they assign a specific selection coefficient to each locus under consideration, with the selection coefficient remaining constant within each seasonal period. However, in nature, selection acts on the phenotype and genome dynamics can be quite different when selection is modeled at the phenotypic level with fluctuating optima, rather than with a fluctuating selection coefficient acting directly at specific loci. In particular, under a fluctuating optima model, the selection pressure acting at loci will decrease as the population approaches the new optimum. Thus, we might expect very different patterns of genome dynamics in a model integrating quantitative genetic and population genetic approaches in exploring the genomic signature of fluctuating selection in different conditions.

Here, we develop a general modeling framework integrating population genetic and quantitative genetic approaches, implemented in the powerful and flexible simulation software SLiM (Haller and Messer 2023). As in other quantitative genetic models, we parameterize the model by assigning a trait heritability and defining specific trait optima and fitness functions based on phenotypic values. Like population genetic models, we specifically consider individual loci genome-wide and track frequency changes over time due to the forces of genetic drift and selection. We sought to identify:

1. What is the parameter space in which fluctuating phenotypic optima lead to a clear genomic signature of persistent oscillating allele frequencies?
2. Under what circumstances can the genomic signature of fluctuating selection be inferred from allele frequency and phenotypic data?

## Results

To answer our main questions, we used SLiM V4.2.2 (Haller and Messer 2023) to compare the genome dynamics in different models of the temporal pattern of the optima across seasons. We compared a model with no relationship between the phenotype and fitness as a null model, a model with a single shift to a new constant optimum, and four models of fluctuating optima across time: (1) instantaneous optima change with two equal seasons, (2) gradual optima change with two equal seasons (simple gradual), (3) gradual optima change with four uneven seasons and even optima (complex gradual I), and (4) gradual optima change with four uneven seasons and uneven optima(complex gradual II). In all models we varied the parameters for heritability (h^2^) and the number of causative loci (L). In three of the four fluctuating optima models, we also examined the effect of the parameter for the distance to the optimum (Δ), which determines the strength of the selective event of a changed optimum. Finally, in the two models with two equal seasons, we varied the season length (Gen) as a parameter. After running simulations accounting for ecological and genetic parameters, we used ggplot2 (Wickham 2016) to visualize the dynamics in allele frequencies and phenotypes over time across the parameter combinations. Furthermore, we used spectral analysis, a mathematical tool that decomposes a time-domain signal into its frequency-domain representation, to detect seasonality in allele frequencies, assuming cyclic behavior of selection forces. An overall schematic of our modeling approach, as well as the different models considered, is in Figure 1.

**Figure 1.**
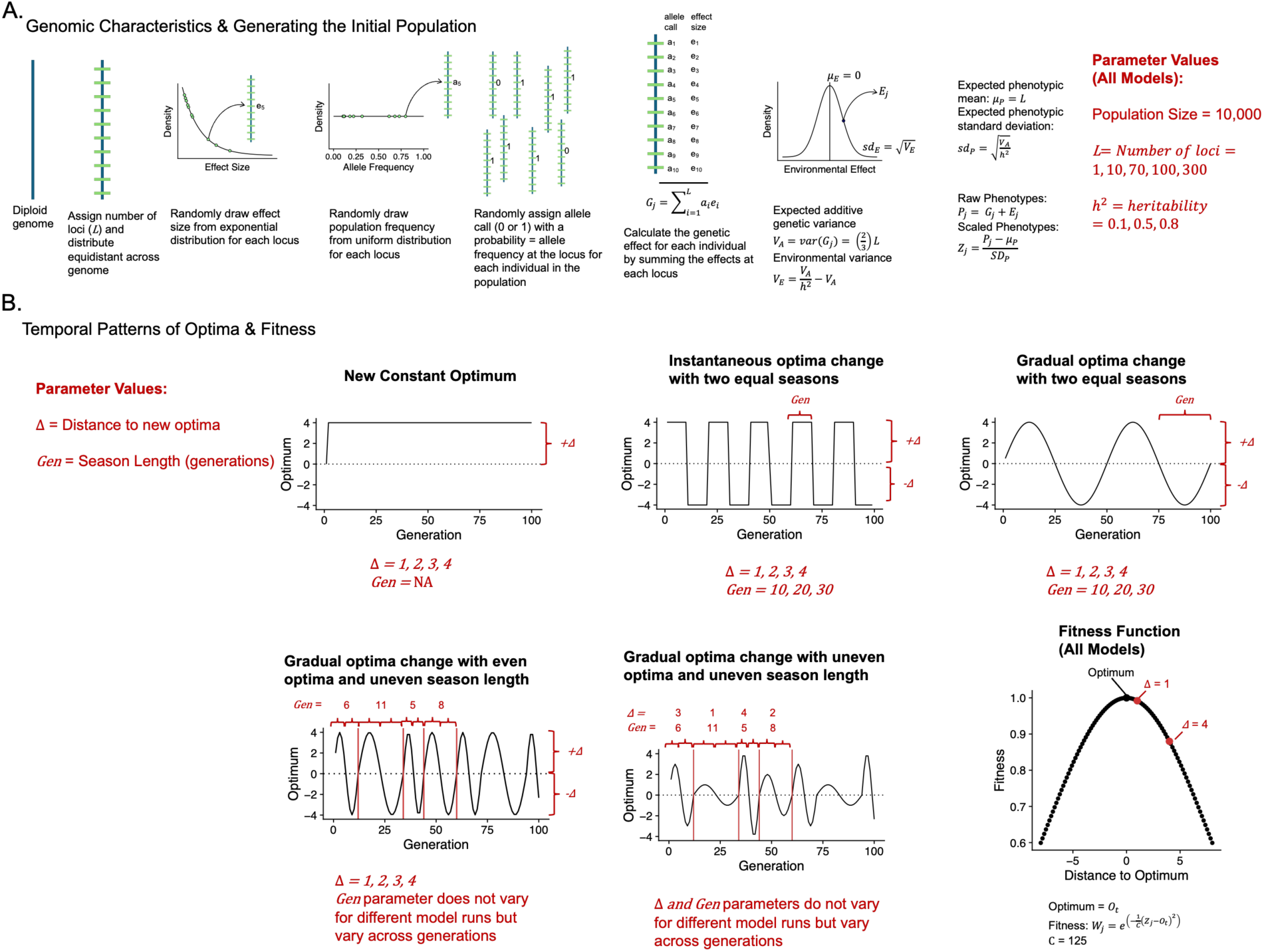
Schematic of the model simulations in SLiM with the model parameters defined. A. The genomic characteristics and process of generating the initial population allele frequencies and phenotypes. B. Overview of the models we considered excluding the null model showing the different patterns of optima over time and the fitness function used in all models.

Below, we summarize the major results, emphasizing the effects of different parameters and differences between the models. A full description of all results for all models is given in the supplementary information.

### Genome dynamics in null and constant selection models as a baseline comparison to fluctuating selection

Comparing fluctuating optima models with null and new constant optimum models is indispensable to fully understanding how fluctuating selection affects allele frequencies and phenotypes; thus, we present these results first to provide a baseline for comparison. Under both null and constant scenarios, neither allele frequencies (Figure 2) nor phenotype trajectories display clear seasonal oscillations, as confirmed by spectral density analyses (see Supplemental Information for more details: Sections 1 and 2 for null and constant selections, respectively).

**Figure 2.**
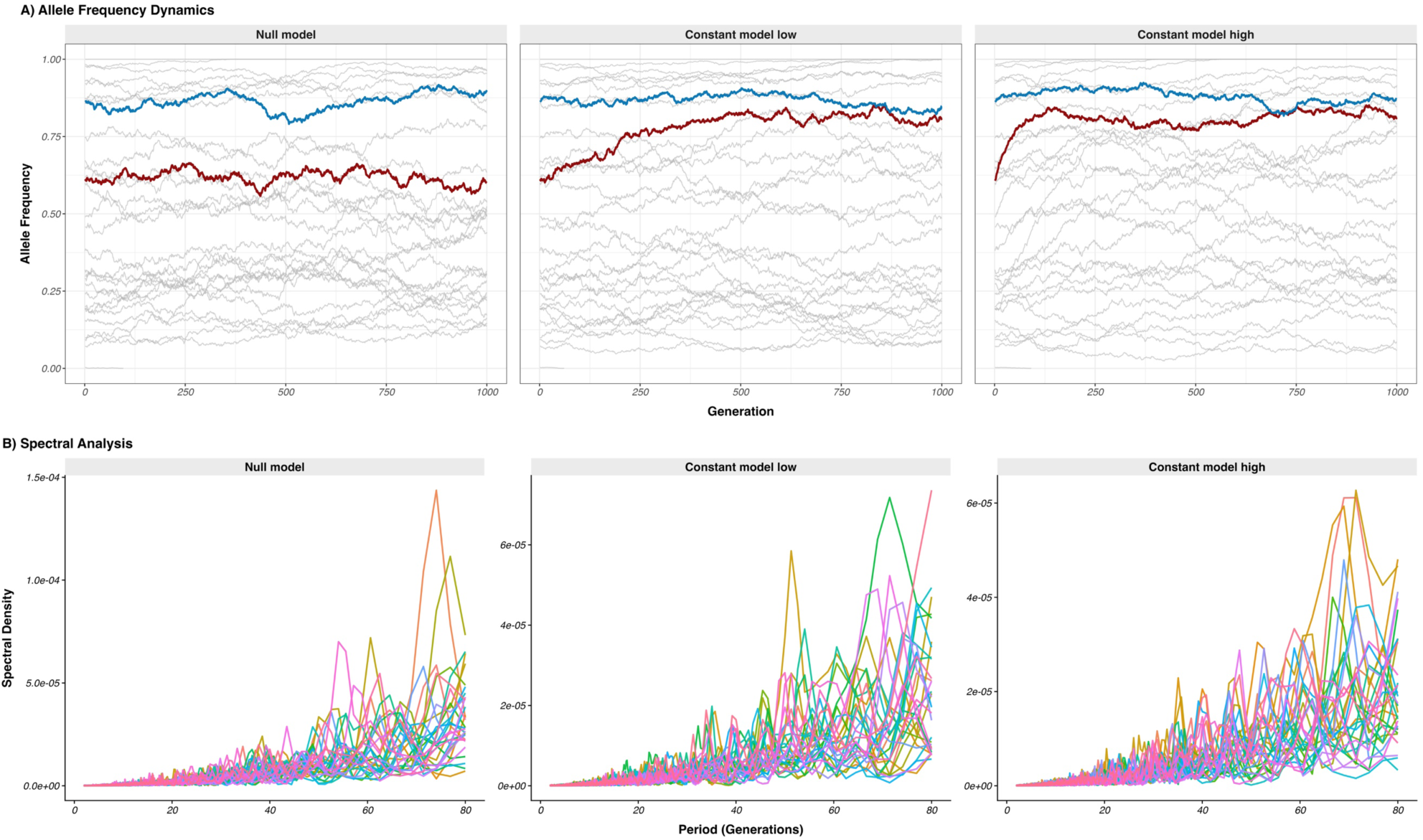
Comparison of polygenic allele frequency trajectories (L = 100 loci) in panel A and corresponding spectral densities (each line representing a single replicate) in panel B for the null model two constant optima models with low (Δ = 1 & h^2^ = 0.1) and high (Δ = 4 & h^2^ = 0.8) parameter combinations. For the allele frequency panel (A), only 30 loci of 100 are shown; the highlighted lines indicate loci with different initial frequency and effect sizes (dark-red position having highest effect with blue having the smallest effect). There is no clear peak in the spectral analysis that would be characteristic of fluctuating selection.

In null models, there are random fluctuations with no discernible pattern mainly because allele frequencies change solely via genetic drift. That is, all the genetic architectures show similar allele frequency dynamics, and loci with terminal initial frequency fix (Supplemental Information, figure 1.1). By contrast, constant models exert steady selective pressures, leading to predictable shifts in allele frequencies and phenotypes until they plateau, either when certain alleles fix, are lost, or when the population reaches its optimum. The plateauing behavior is particularly pronounced in polygenic models (Supplemental Information, sections 2.1 and 2.2 for allele frequencies and phenotypes, respectively, under constant selection). Monogenic traits under strong selection often go to fixation more quickly, while polygenic and oligogenic traits tend to remain within specific frequency ranges over many generations. Such behavior does not appear under the null model, where frequencies drift randomly. Despite these distinctions, spectral analyses of null and constant selection models reveal no evident periodicity in either regime (See Supplemental Information, section 1.3 and 2.3 for null and constant selections, respectively). Consequently, it is difficult to pinpoint out the effects of individual parameters via spectral analysis in these two models.

### Higher heritability, stronger selection, and longer season length produce greater evolutionary changes across all fluctuating selection models

In our models of fluctuating selection, as has been observed often in evolutionary models previously (Walsh and Lynch 2018), we observed larger changes in allele frequencies when the heritability was higher, the shift in the optimum was larger, and/or the season length and thus selection period was longer. This pattern is clear in the polygenic (100 causative loci) model of instantaneous optima change with two seasons, where one can see that seasonal oscillations in both the phenotype and the allele frequencies across the genome showed greater change when comparing low heritability to high heritability, a small shift in optimum to a larger one, and a short season length to a longer one (Figure 3A & 3B). These overall effects were consistent across all models that included selection (Supplementary Information, sections 4 and 7), showing that the population tracked the change in optima more closely in parameter combinations with high heritability, stronger selection, and longer selection periods, as expected.

**Figure 3.**
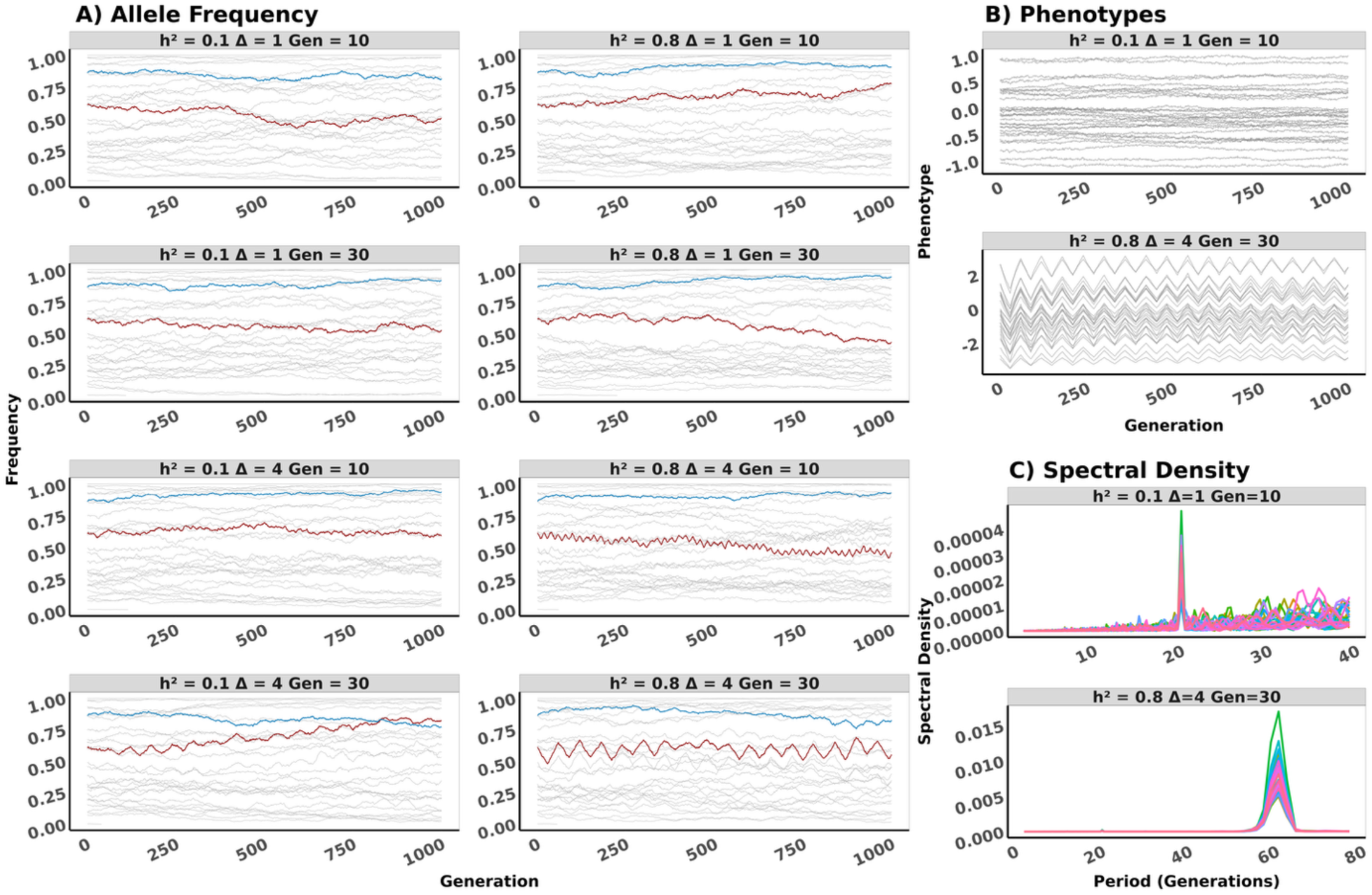
Evolutionary dynamics from the instantaneous optima change with two equal seasons model for a population with polygenic trait (loci = 100). **A**. represents allele frequency dynamics and shows parameter combinations from a single replicate for 1000 generations. Each line in the plot represents the frequency trajectory of a locus among 30 loci randomly sampled from 100 loci, and the highlighted lines indicate loci with different initial frequencies and effect sizs (dark-red position having highest effect with blue having the smallest effect). **B.** Distribution of population mean phenotype dynamics representing 30 replicates over 1000 generations. **C.** The spectral density from allele frequency for all 2000 generations; each colored line represents a single replicate. The peak in the spectral plots marks a complete cycle of season length. The variables are heritability (h^2^), selection strength (Δ: magnitude of the shift in the optimum), and the season length in terms of number of generations (Gen).

We performed spectral analysis to identify when the genome-wide allele frequency patterns in the population showed periodicity (also termed recurrent seasonal shifts). In our fluctuating selection models, spectral analysis confirmed the effects of heritability, selection strength, and season length by showing more pronounced peaks under these same parameter values (Figure 3C; Supplementary Information 4.2), indicating a stronger signal of periodicity for these parameter combinations.

### Clear fluctuations in allele frequencies persist for thousands of generations in fluctuating selection models in most parameter combinations

Many previous population genetic models have modeled fluctuating selection by altering the selection coefficient at specific loci between seasons while keeping it constant within each seasonal cycle (Huerta-Sanchez et al. 2008; Miura et al. 2013; Cvijović et al. 2015; Wittmann et al. 2017; Wittmann et al. 2023; Johnson et al. 2026). Some of these models show that fluctuating selection can change the site-frequency spectrum, while others show that it increases the likelihood of fixation compared to a neutral expectation. We find that our shifting optima models of fluctuating selection typically lead to persistent oscillations in allele frequencies and phenotypes over at least 2,000 generations. We see no evidence for an overall diminishment of these oscillations in either the allele frequency patterns or the phenotypic pattern across this timescale (Figure 3; Supplementary Information, section 4). This result is consistent for most parameter combinations in all of our fluctuating selection models (Supplementary Information). The two exceptions to this overall pattern are 1) when the parameter combination does not lead to discernable seasonal oscillations across the entire time period, or 2) when the phenotype is determined by few loci and these go to fixation, thus preventing future oscillation. We detail the parameter space in which we observe these two cases in the sections below.

To assess whether fluctuating selection maintains polymorphism beyond our 2,000 generation simulations to equilibrium, we tracked the fate of the initial positions (QTLs) over 10Ne generations. We compared null and fluctuating selection models with (mutation rate, µ = 1e-07) or without mutations. For each generation and parameter combination, considering high (*h^2^* = 0.8, *Gen =* 30, *Δ* = 4) and low (*h^2^* = 0.1, *Gen =* 10, *Δ* = 1) values, we calculated the initial fixation as the fraction of fixed or lost positions in the first generation. Our results show that across all parameter combinations, initial QTL fixation and loss increased through time and approached one at a similar rate to the null model, with most alleles fixing by ∼50,000 generations (Supplemental Information SI 3.1 and SI 3.2). This indicates that the initial QTL variation was eventually depleted across all models, regardless of whether selection was null or fluctuating, or whether the model allowed new mutations.

While in many parameter combinations the seasonal oscillation of allele frequencies is obvious visually (e.g. Figure 3A, bottom right plot), even in parameter combinations where these oscillations are more subtle, spectral analysis is often able to detect the recurrent periodicity. This ability is clear in the example of the polygenic model of instantaneous optima change with two seasons in the combination of low heritability (h^2^ = 0.1), weaker selection strength (Δ = 1), and shorter selection length (Gen = 10) where the oscillations in allele frequencies are not visually obvious (Figure 3A) but spectral analysis detects a clear peak at the correct periodicity of 20 generations for a full seasonal cycle of 10 generations in each of the two seasons (Figure 3C). This same ability of spectral analysis to correctly identify periodicity even for parameter combinations when individual allele frequency oscillations are subtle occurs for most of our fluctuating selection models (Supplemental Information, section 4.2). Although spectral analysis does successfully detect periodicity in parameter combinations with more subtle allele frequency shifts, we note there is more variability in spectral density in these parameter combinations (see Figure 3.C; Supplemental Information, section 4.2.4).

We also examined how the number of loci contributing to the trait in instantaneous selection model affects the observed evolutionary dynamics. Generally, the patterns described above for a polygenic model extend to models with fewer causative loci, although fewer positions affecting the trait for a given heritability tend to produce more pronounced oscillations in allele frequencies and phenotypes (Figure 4A & 4B).

**Figure 4.**
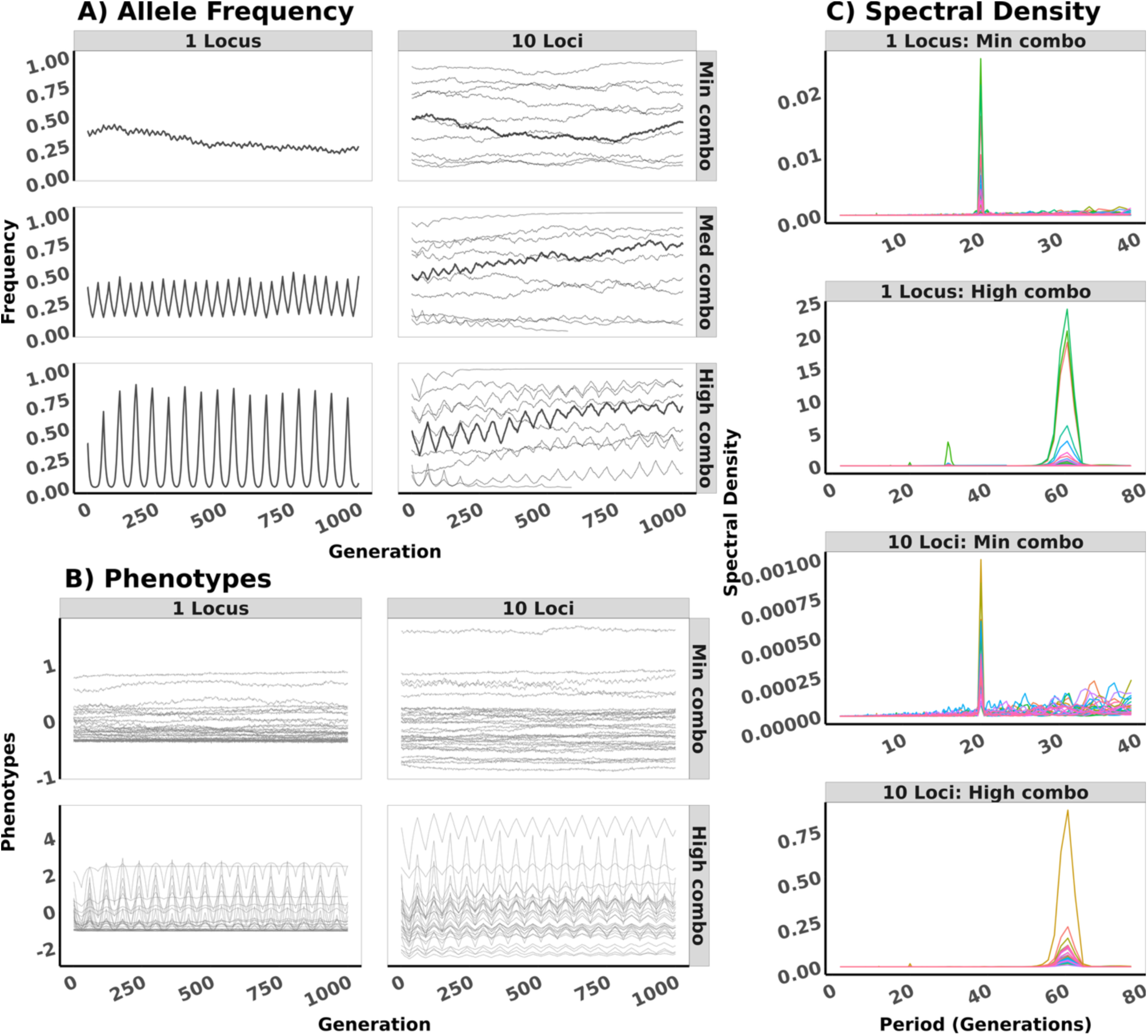
Comparison of genetic architectures (1 locus and 10 loci traits) under instantaneous fluctuating selection. **A.** represents allele frequency dynamics for a single replicate for small (Min combo: h^2^ = 0.1, Δ = 1, Gen = 10), medium (Med combo: h^2^ = 0.5, Δ = 2, Gen = 20), and large (High combo: h^2^ = 0.8, Δ = 4, Gen = 30) parameter combinations over 1000 generations. Each line represents the frequency trajectory of a locus, with a highlight of the dynamics in a single locus. **B.** Shows distribution of population means phenotype dynamics for 30 replicates over 1000 generations for both small and high parameter combinations in each architecture. **C.** Displays spectral density of allele frequency over 2000 generations, where each colored line represents a single replicate for both small and large parameter combinations for each architecture. The large peak in the spectral plot marks a complete cycle of the season length.

The differences among the genetic architectures shown in Figure 4 are driven by the fact that the heritability of the trait is held constant in each case. Thus, for a given heritability, having fewer causative loci means that each of those loci will contribute a larger share to the percentage of genetic variation for the trait and thus have a larger genetic effect. Therefore, the observation that individual loci show greater fluctuations when there are fewer loci is driven by this effect rather than by the genetic architecture itself. We observe similar oscillations in the phenotype for different numbers of causative loci, though in the case of a single locus, we do see cases where the phenotype becomes constant when one allele becomes fixed, which does not occur when more than one locus affects the trait (Figure 4B).

Spectral analysis successfully identifies periodicity regardless of the number of causative loci. For a given heritability, models with fewer causative loci often show distinct peaks with high spectral densities, while polygenic traits typically have lower spectral densities (Figure 4C & Supplemental information 4.2). Nevertheless, the greater variation in spectral peaks across replicates in single-locus traits may demonstrate the susceptibility to genetic drift and high fixation rate. As a result, even though the shifts are more subtle when there are more loci sharing the effect, the analysis across all of them gives a well-defined signal. Ultimately, despite the differences, spectral analysis reliably detects seasonality across all genetic architectures, particularly when considering aggregate behavior across loci.

While the changes in amplitude of our allele frequency oscillations in different parameter combinations are visually obvious, we can quantify the average amplitude across time for different parameter values. We calculated the absolute value of the difference in allele frequency at the end of each season and averaged these values across 2000 generations. The effects of each of our main parameters and the number of causative loci on the amplitude of the allele frequency oscillations is shown in Figure 5. Here, once again, the larger amplitudes when fewer loci contribute to the heritability of the trait are evident, as are the individual impacts of each parameter on changing the amplitude of allele frequency oscillations.

**Figure 5.**
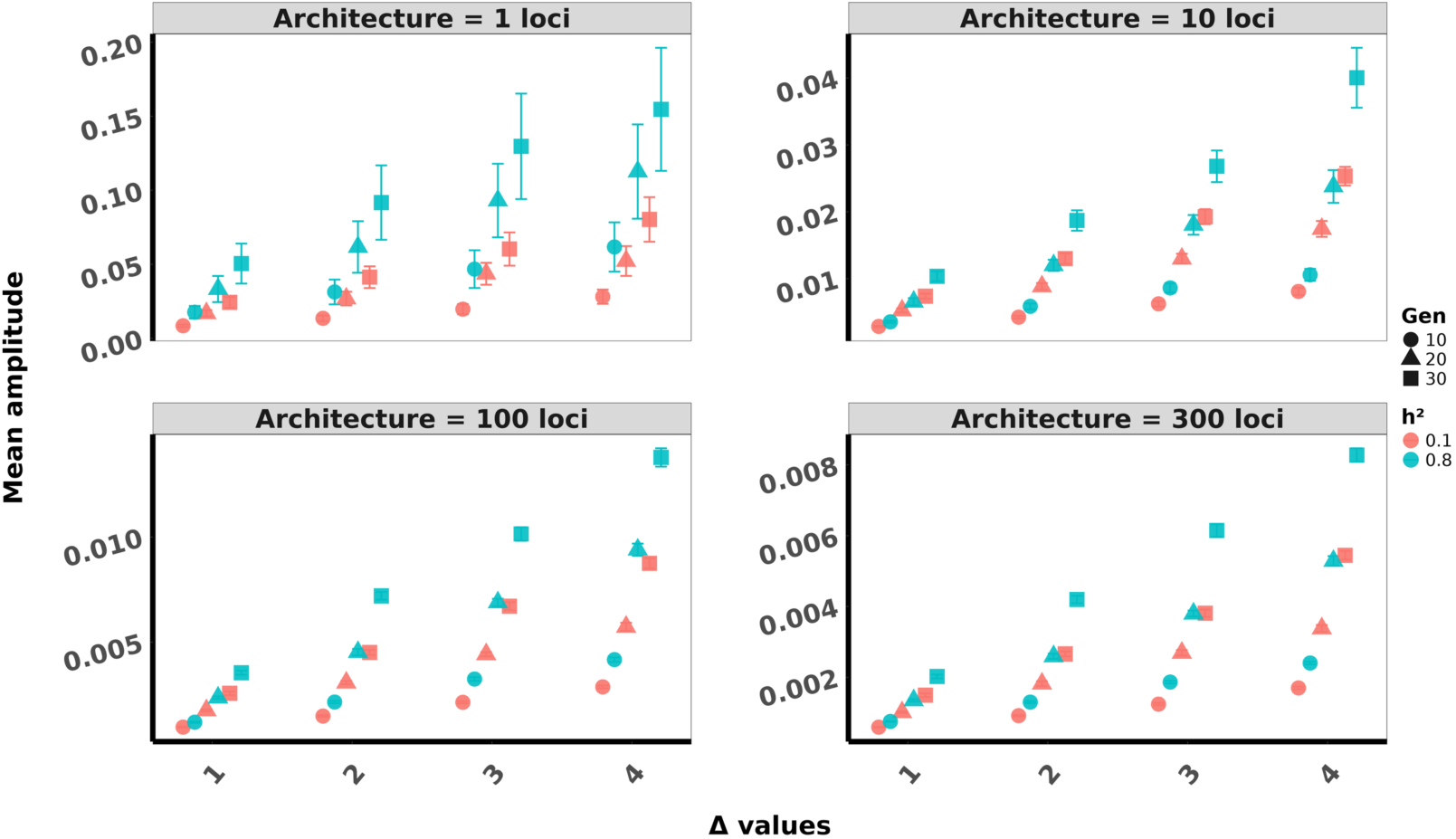
Mean amplitude of allele frequency oscillations averaged over 2000 generations versus the distance to the new optima for the instantaneous two season model. Note the y-axis has a different range for each plot. The amplitude is defined as the absolute value of the difference between the allele frequency at the end of each season. These values are calculated for each locus and then averaged across loci and then across 2000 generations. Different shapes denote different season lengths and different colors denote different heritabilities. Each panel represents a different number of causative loci. Error bars are +/- 1 standard error, calculated across replicate simulation runs.

### Genetic architecture influences the rate of fixation of alleles

In addition to the described means of differentiating behaviors per genomic architecture, we calculated the percentage of fixed loci in our simulations. The fixation rates are percentages that were calculated as the ratio of the total number of fixed loci to the total number of loci from all replicates, multiplied by 100. The results show that when relatively few loci affect the trait, the proportion of fixed alleles is substantially higher than in polygenic models (Supplemental Information, section 5). In polygenic traits, increasing the number of causative loci does not notably alter fixation rates, and other parameters (e.g., heritability or selection duration) have little impact on this outcome (Supplemental Information, section 5). When there are few causative loci and they become fixed, obvious continued oscillations in the phenotype and in allele frequencies cease. Thus, this increased rate of fixation when few loci influence the trait highlight a key parameter showing when we fail to see persistent seasonal oscillations in allele frequencies.

### Spectral analysis succeeds in detecting fluctuating selection with fewer, more realistic sampling timepoints when parameter combinations are favorable

The spectral analysis we have employed thus far uses allele frequency data from every generation of the simulations. Obviously, this sampling density and timescale is much greater than any dataset that would be collected on a real population. We down-sampled our simulated datasets from the instantaneous model to approximate a more realistic dataset to determine if spectral analysis is likely to detect fluctuating selection in empirical time-series allele frequency datasets. In some experiments, such as those studying seasonality, investigators attempt to time sampling for the ends of each season, or for the midpoints and ends of each season (Bergland et al. 2014; Machado et al. 2021). Thus, we used these two strategies to “sample” our simulated datasets either at the end of each seasonal cycle or at the endpoints and midpoints of each seasonal cycle, starting from after the first generation interval. We chose 10 sampling points as a reasonable number in terms of cost and effort, while still encompassing several seasonal cycles. For favorable parameter combinations of high heritability, strong selection, and long season length, spectral analysis succeeds in detecting fluctuating selection at the correct periodicity (Figure 6), demonstrating that the method can work in a much smaller sampling dataset. Not surprisingly, in parameter combinations that produce more subtle allele frequency oscillations, spectral analysis with 10 sampling points does not produce a clear signal (Supplemental Information, section 6), requiring more sampling points to identify fluctuating selection.

**Figure 6.**
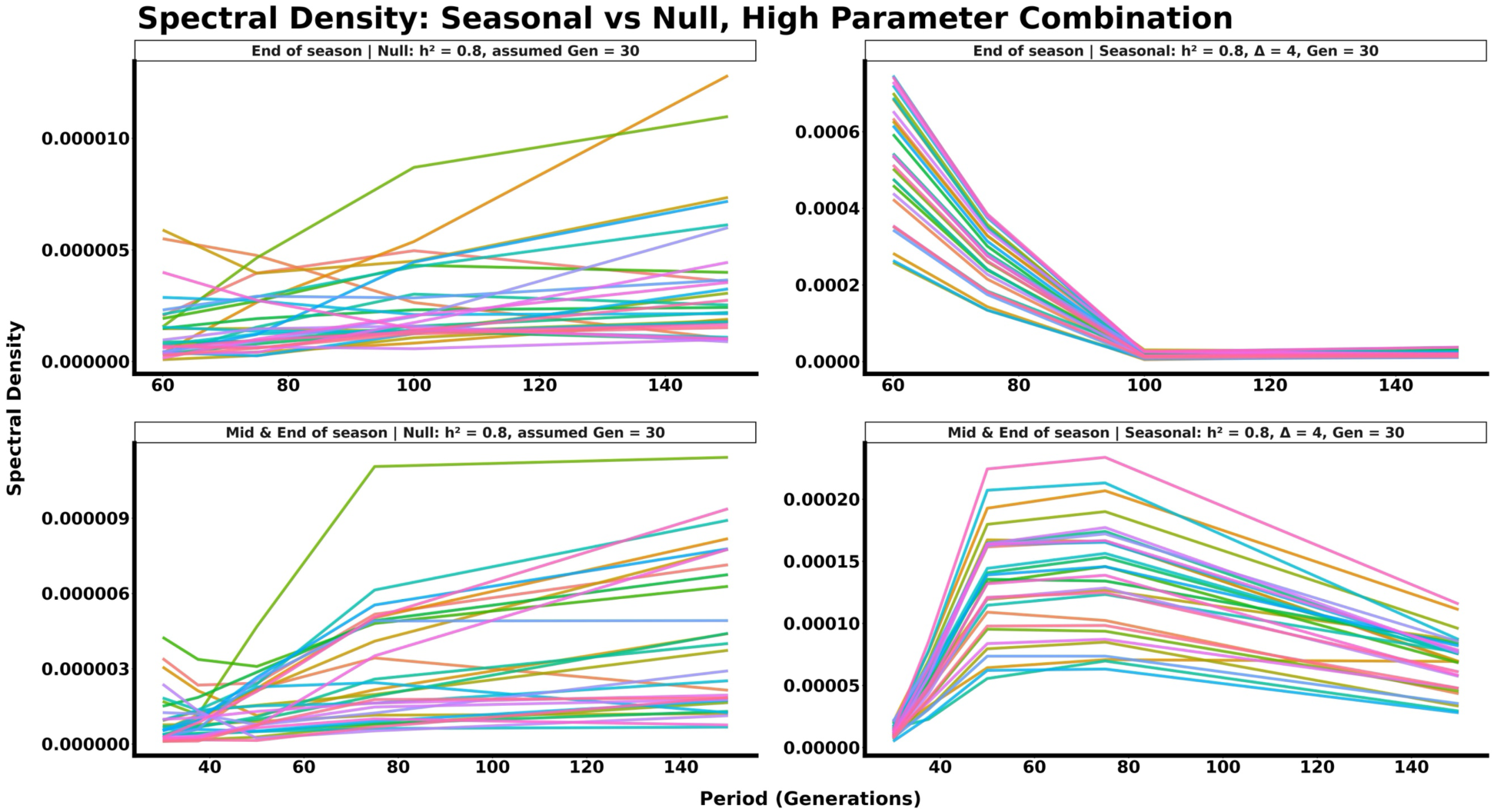
Spectral analysis for 10 sampling points. Panels on the left show the spectral density versus the period in generations for a null model. The panel to the right shows the spectral density versus the period in generations for the instantaneous fluctuating selection model with sampling occurring at the ends of each season (top) or the ends and midpoints of each season (bottom). In both cases, the spectral density peaks at the correct interval of 60 (one full seasonal cycle).

**Figure 7:**
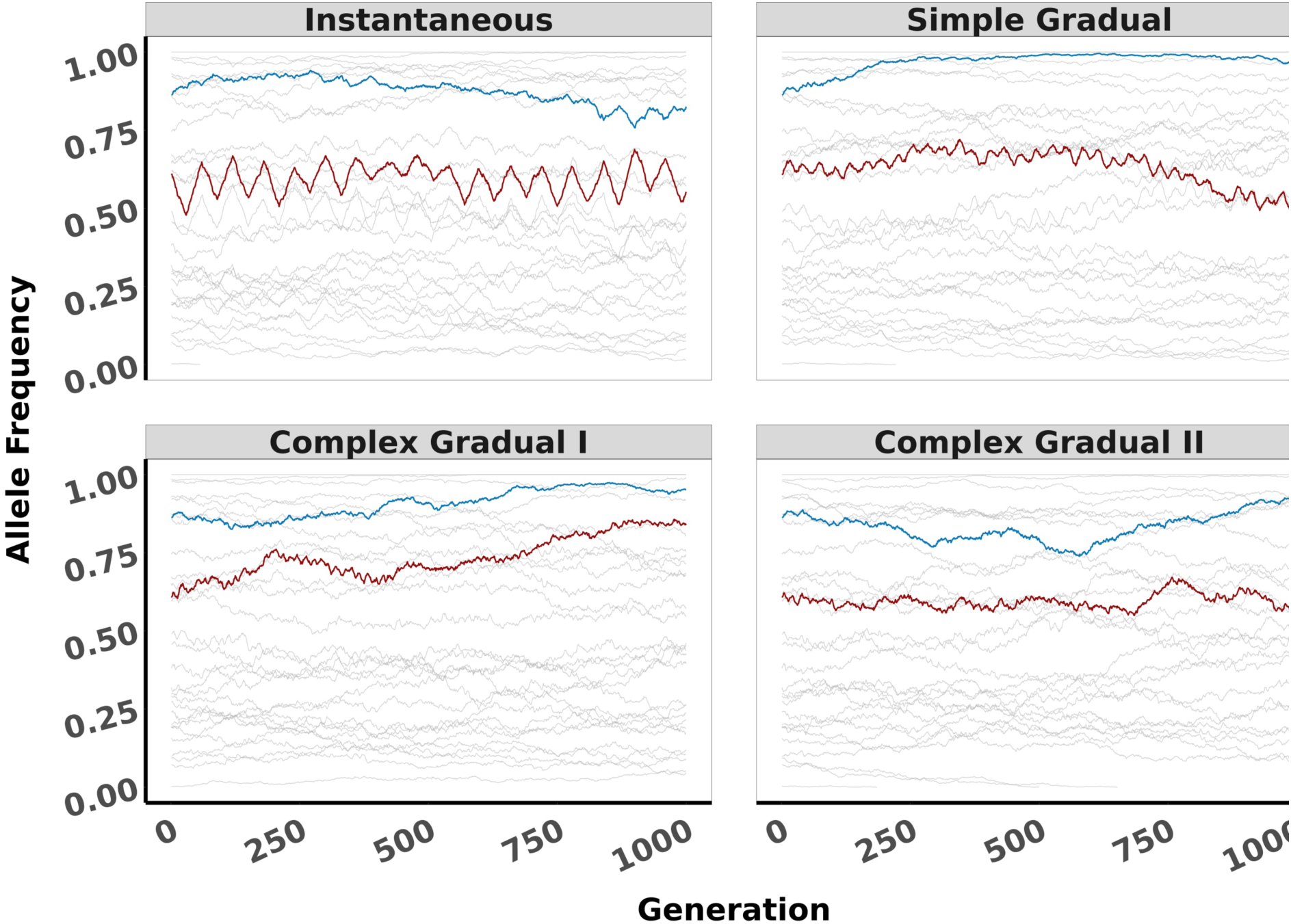
Allele frequency dynamics at 30 of 100 simulated loci comparing all fluctuating selection models. The instantaneous model shows clearer seasonal oscillations, followed by the simple and, finally, complex gradual models. For two equal season models(instantaneous and simple gradual), the heritability is h^2^ = 0.8, with Δ=4, and Gen = 30. For the four uneven season length models with different distances to optima (Complex gradual II), Δ values were set to 3, 1, 4, and 2 for generations 12, 22, 10, and 16, respectively. In the version of even optima (Complex gradual I), the value is Δ=4 for all seasons. The highlighted lines indicate loci with different initial frequencies and effect sizes, where the dark red line shows positions with the highest effect, while the blue shows positions with the smallest effect.

### More complex seasonal patterns are more difficult to detect

Above, we emphasized results from our most straightforward model, the instantaneous optima change with two seasons model. However, we also explored more complex and closer to realistic scenarios including a model of gradual optima change with two equal seasons and models of gradual optima change with uneven season lengths where each season consisted of two equal sub-seasons with even or uneven optima. These gradual-change models introduce selection more slowly, with smooth transitions between seasonal optima; hence, they may reflect natural environments where conditions do not shift abruptly. As the temporal pattern of change in the optima becomes more complex, it becomes more difficult to visually observe seasonal oscillations in allele frequences for a given parameter combination (Figure 8). Our most complex model is the gradual optima change with uneven season length models because it involves different season length (12, 22, 10, and 16 generations) in a single cycle, and it was further subdivided into two variants 1) One with a Δ value that varies across models but not generation and 2) another that Δ remains the same across models, but changes depending on the season length (see Figure 1). This setup reflects scenarios with an asymmetric season length throughout the year or across life stages, complicating evolutionary dynamics and blurring the distinction between allele frequency changes driven by selection and by drift. In these more complex models, we see similar effects of heritability, selection strength, and the number of causative loci as we described above (Supplemental information, sections 7, 8 and 9).

**Figure 8.**
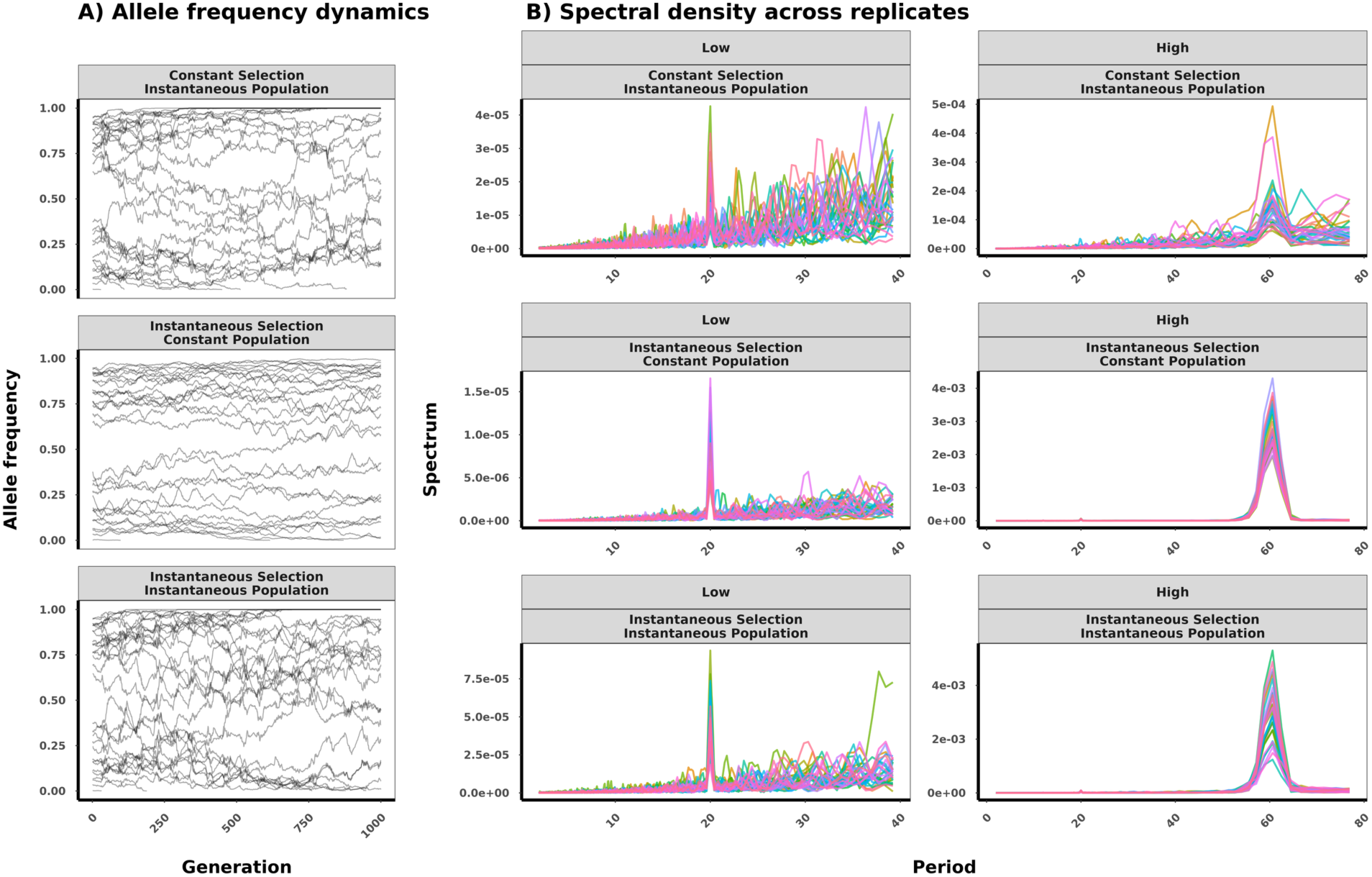
Comparison of allele frequency dynamics A), and Spectral analysis B for models under constant selection with fluctuating population size, fluctuating selection with constant population size, and fluctuating selection with fluctuating population size, given parameter combinations under the polygenic trait model (number of loci = 300). Where low is the combination low parameters (h^2^ = 0.1, Gen = 10, D = 1) and high is large parameter combination (h^2^ = 0.8, Gen = 30, D = 4). For the allele frequency dynamics, only large parameter combinations and 30 positions are shown for 1000 generations while the spectral analysis takes in all 2000 generations.

Despite more subtle seasonal allele frequency shifts, spectral analysis can identify periodicity effectively for most parameter combinations (Supplemental information 8.2 & 9.2), though we expect detection to become more challenging when fewer generations are sampled. The limits to detecting periodicity via spectral analysis even with perfect sampling become clear in the highly complex gradual optima change with uneven season length model in parameter combinations with low heritability and/or weak selection (Supplemental information, figures 8.2.1 & 8.2.2 upper left corner). These parameter combinations are some of the few in which clear spectral peaks are difficult to detect for any of our fluctuating selection models and where the patterns look similar to those we see in null or constant models. While the complex model with varying *Δ* values and season lengths may produce a peak under certain conditions, for instance low heritability, this peak reflects mainly the selection strength during that period (Supplemental information section 9.2). Consequently, it emphasizes how selection strength influences the ability to detect such signal from genomic data. Once again, we observe that for a given heritability, while models where fewer loci contribute to the heritability provide more readily observable patterns in allele frequencies, spectral analysis across many loci produces more consistent results (Supplemental Information, section 10). Thus, in polygenic models, one is more likely to reliably detect fluctuating selection genome-wide, though it may be less discernable when examining individual loci in isolation. The results emphasize that the complexity of the fluctuating selection pattern significantly influence how populations respond to fluctuating environments.

### The effect of fluctuating population size in addition to fluctuating selection

In seasonal environments, population sizes can also fluctuate with the seasons. We incorporated the effect of a fluctuating population size into our instantaneous two season model to explore the effect of both the optima and the population size fluctuating concurrently, where in one season the population size was 10,000 and in the other 1,000, and compared this model to a model with constant selection and the model with fluctuating selection with a constant population size. The introduction of fluctuating population size into the selection models visibly altered the allele frequency dynamics. Figure 8A compares the dynamics under constant selection with fluctuating population size, fluctuating selection with constant population size, and fluctuating selection with fluctuating population size, for two parameter combinations under the polygenic trait model (number of loci = 300), a high combination (*h^2^* = 0.8, *Gen =* 30, Δ = 4) and a low combination (*h^2^* = 0.1, *Gen =* 10, Δ = 1). Adding fluctuating population size introduces stochasticity into the allele frequency dynamics, with increased variation in allele frequency dynamics occurring during the intervals of small population size, and some positions reach fixation more readily than under fluctuating selection alone (Supplemental Information, section 11).

In the models including fluctuating selection, spectral analysis produces a spectral peak recovering the seasonal signal, though the signal is weak at lower parameter combinations. In the model with the addition of fluctuating population size to fluctuating selection, there is greater spectral variation, even though the peaks are well defined in both cases. This suggests that the addition of fluctuating population size adds variability to the allele frequency oscillations, making them more difficult to recover clearly.

In the model with constant selection and fluctuating population size, spectral analysis does recover a peak corresponding to the periodicity of population size changes. This result is likely due to allele frequencies shifting predictably sometimes in response to selection at high population sizes and less predictably, and thus often in an opposite direction when population sizes are small, producing a signal that mimics what is seen under fluctuating selection, or sometimes due to the periodic occurrence of bottleneck events. Importantly, even under null model with fluctuating population, recurrent bottlenecks may generate a peak at the seasonal frequency in the spectrum, and whether that peak is detectable depends on the severity, duration, and regularity of the bottlenecks. Therefore, the current result highlights the challenges associated with reliably assigning causation based on allele frequency dynamics alone. However, the spectral analysis and allele frequency dynamics are much clearer in the case of fluctuating selection (Figure 8B), indicating it may be possible to distinguish between these two forces in some cases.

## Discussion

Our results show the expectations for allele frequency dynamics genome-wide under different parameter combinations within a modeling paradigm where the phenotypic optima fluctuate through time. Overall, we showed that fluctuating optima models result in recurrent oscillations in allele frequencies that can persist stably for well over 1000 generations. As would be expected from quantitative genetic theory, the amplitude of these oscillations was most dramatic when the shift in optima was large, season length was long, and heritability was high (Barton and Turelli 1989; Hill and Mackay 2004; Bijma 2011). In addition, we employed a new approach to detecting fluctuating selection from allele frequency dynamics, spectral analysis. We showed this approach can successfully recover the signal of fluctuating selection in all but the most complex simulations.

### A fluctuating optima modeling paradigm

Our modeling paradigm described above is a phenotype-based modeling approach (Lande 2007; Lande 2009) of fluctuating selection where recurrent changes in environmental conditions shift the optimal phenotype producing emergent shifts in selection on the causative loci influencing that phenotype, which is in contrast to the many commonly employed modeling approach to study fluctuating selection, where the selection strength is modeled directly on the focal loci with a constant value throughout each selection period (Ellner and Hairston Jnr 1994; Wittmann et al. 2017; Bertram and Masel 2019; Glaser-Schmitt et al. 2021; Johnson et al. 2026). We focus our results on describing the expectations for recurrent allele frequency dynamics from standing genetic variation across experimental timescales within this fluctuating optima paradigm, defining expectations for both simple and complex patterns of optima change across time.

Our modeling paradigm made several simplifying assumptions across most models, including non-overlapping generations, constant population sizes, no new mutations, and fixed recombination rates, all of which may limit the extent to which these results can generalize. In addition, we did not allow for any genotype-by-environment interaction at any loci. Previous quantitative genetic models of fluctuating selection have frequently identified the conditions under which fluctuating selection is expected to lead to the evolution of phenotypic plasticity driven by genotype-by-environment interactions (Via and Lande 1985; Lande 2014; Oostra et al. 2018). Our models did not allow for the evolution of plasticity in this way, removing a major potential way for populations to adapt to fluctuating selection pressures. Thus, our results shed light on whether it is ever possible for fluctuating selection to drive recurrent, stable, allele frequency oscillations in a population across a timescale of hundreds to thousands of generations, given these simplifying assumptions and provide a baseline for future modeling efforts relaxing these assumptions.

### Increasing model complexity highlights limits to detecting fluctuating selection

The ecological conditions experienced by natural populations will often be much more complex and dynamic than our simplest model where the optima changes on a consistent seasonal cycle and always to the same values each season. In nature, temporal patterns of selection will often vary in both timing and strength (Grant and Grant 2002; Bell 2010; Abdul-Rahman et al. 2021; Roberts Kingman et al. 2021), and allele frequency dynamics are known to be affected by other factors such as variation in population size (Bertram and Masel 2019; Kim 2023; Nunez et al. 2024). In our models, we explored both more complex temporal patterns of selection, and a case where population size varies concurrently with fluctuating optima. Although the simulated more complex fluctuating selection models do not correspond to a specific biological scenario, they rather serve as one example of more complex patterns over time, of which there are many varied examples in nature (Park 2019; Hernández-Carrasco et al. 2025). A good analogy is the effect of El Niño on the southern oscillation index where the oscillation in the index is associated with the variation in fish recruitment in the Pacific Ocean. However, these oscillations do not happen every year and occur in addition to the regular annual seasonal cycle (Shumway and Stoffer 2017), a scenario that can be experienced by a population under various evolutionary forces. When there are four uneven season lengths with/without same magnitude of the shift in the optimum, the population struggles to adapt; thus, the complex oscillatory dynamics. Nevertheless, gradual models better represent complex natural environments where multiple seasons may have different generation turnover. These more complex scenarios, especially those with four uneven season lengths, create subtle patterns that can resemble null or constant selection when parameters (e.g., selection strength, heritability) are low. This observation is consistent with prior studies showing that the outcomes of fluctuating selection depend on the selection strength, genetic variance, and sampling time (de Vladar and Barton 2014; Wittmann et al. 2017; Johnson et al. 2026). Indeed, a particularly long or short season under these gradual models can either strengthen or weaken selection signals, respectively, making detection more difficult (Cvijović et al. 2015; West and Mobilia 2020; Abdul-Rahman et al. 2021) for the latter. We observed a similar result when incorporating the added complexity of population size fluctuating along with selection, with increased variability in allele frequency trajectories, though spectral analysis still succeeded in detecting a clear signature of fluctuating selection (Figure 8).

### Spectral analysis as a novel approach to detecting signatures of fluctuating selection: potential, limitations, and future directions

We employed spectral analysis to tackle the challenge of identifying recurrent oscillatory signals from allele frequency trajectories. With complete data for every generation simulated, this approach successfully discerned cyclical patterns in allele frequency across most parameter combinations, particularly in simpler models (i.e., the instantaneous and two equal seasons models). In empirical datasets, ideally, sequencing multiple sequential generations for a long time would help identify seasonality from genomic data using spectral analysis. However, its feasibility is almost impossible due to the associated cost and labor. To explore whether spectral analysis can identify fluctuating selection with a more realistic dataset, we subsampled our simulated datasets to consist of only 10 sequencing timepoints. Spectral analysis is able to correctly identify the signal of fluctuating selection at the correct periodicity under favorable parameter combinations (high heritability, strong selection, and long season length; Figure 6), demonstrating that spectral methods provide a promising framework to distinguish signals of fluctuating selection, worthy of future study. While our simulations have demonstrated the potential of this method, our work also highlights important limits of the method and areas that require more extensive study to define the conditions under which spectral analysis is likely to succeed that we detail below.

We identified limits to using spectral analysis to identify fluctuating selection, even in cases with complete data across 2,000 generations. In more complex patterns of fluctuating selection (as in the Complex gradual I and II) along with weak selection and/or low heritability, spectral analysis results mirrored those from our null or constant selection only models, demonstrating that highly complex temporal patterns will be challenging to identify even with dense sampling. Future work could use the distribution of spectral analyses from null simulations to quantify the null distribution and then calculate the proportion of null analyses that produce a spectral signal as great or greater than an observed analysis as a potential measure of the significance of the fluctuating signal. Another limit to spectral analysis we identified was its inability to distinguish between fluctuating selection and fluctuating population size in the constant new optima model. While spectral analysis easily distinguished between the fluctuating optima model and either the constant optima only model or our null model, when fluctuating population size acts with the constant optima model, it produces periodic allele frequency changes identified by spectral analysis, mimicking a fluctuating optimum. This may also be true for null selection models with fluctuating population size, where the periodic recurrence of a strong bottleneck can yield identifiable spectral peaks, since these time-series methods are designed to detect periodic patterns (Parzen 1967; Platt and Denman 1975). In contrast, in the weak recurrent bottleneck, the signal would be distorted by subtle changes in allele frequency; consequently, one could not expect any signal. Regardless, all of that would also be determined by the interval between occurrences and the regularity of changes in population size. We considered only a single pattern of fluctuating population size out of a whole range of realistic patterns experienced by different natural populations. Further large-scale simulations of many possible seasonal changes in population size (Gloss and Whiteman 2016; Barghi et al. 2019; Agashe et al. 2023; Holstad et al. 2024) along with different temporal patterns of optima could define the parameter space where spectral analysis can and cannot distinguish between these conditions to test the robustness of spectral analysis in realistic scenarios.

A major challenge for empirical studies of the genome dynamics in fluctuating selection is the need for multiple sequencing timepoints, which can quickly become prohibitively expensive. Phillips et al. argued that taking more samples over time can dramatically improve our ability to understand the complexity of evolutionary dynamics, though they did not specifically examine fluctuating selection(Phillips et al. 2020). While we did not apply our approach directly to empirical data, our framework establishes a foundation for refining spectral methods to detect fluctuating selection, a field that is far less developed than approaches for positive/directional and other forms of selection (Hancock and Di Rienzo 2008; Haasl and Payseur 2016; Qin et al. 2022). While we showed spectral analysis can still succeed with favorable parameters in a realistic (though still larger than many empirical studies) empirical dataset consisting of 10 sampling points, this result did not hold for our unfavorable combination of low heritability, weak selection, and a short season length. Additional simulations of several different sampling schemes across many different models could provide guidelines for the sequencing effort needed to reliably detect fluctuating selection using spectral analysis or other methods and these simulations could then be compared to real empirical datasets. Evolve-and-resequence time-series datasets stemming from fluctuating selection regimes would be especially valuable in validating both the patterns of allele frequencies across time and the use of spectral analysis to detect fluctuating selection.

Finally, while spectral analysis provides a tractable first step, alternative methods such as wavelet transforms or the S-transform may offer greater robustness in detecting oscillations under nonstationary conditions or asymmetric selection regimes (Shumway and Stoffer 2017; Khan and Pierre 2019; Arnab et al. 2023). Furthermore, new computational methods, including machine learning (Greener et al. 2021), may be applied to images of allele frequency trajectories to detect cyclical selection signals which may better handle sparser datasets.

## Methods

We used SLiM V4.2.2 (Haller and Messer 2023) to simulate a set of Wright-Fisher models, including a null model, a new constant-optimum model, and models with fluctuating optima. Except where noted below, the population size was fixed at 10,000 individuals, and the simulation ran for 2000 generations, and we assumed sexual reproduction with an equal sex ratio, random mating, no new mutations, and a recombination rate of 10^-8, a commonly used value in other SLiM models (Haller and Messer 2023) and reflecting the average recombination rate for many mammals (Dumont and Payseur 2008). The computation for this work was performed on the high-performance computing infrastructure operated by Research Support Solutions in the Division of IT at the University of Missouri, Columbia, MO https://doi.org/10.32469/10355/97710 (University of Missouri Cluster (Hellbender)).

### Genomic Characteristics & Generating the Initial Population

We modeled a phenotype with a known set of causative quantitative trait loci (QTL). The number of QTL contributing to the phenotype was set as *L*, with parameter values of 1, 10, 70, 100, or 300. We set a genome size to be approximately 1.2MB for an autosomal chromosome, with QTL spaced equidistantly across the genome. For each QTL, the effect of the locus on the phenotype was drawn from exponential distribution with a rate parameter *λ* = 1, which assumes most loci have minor effects on the phenotype and a few have a large effect size (Barton et al. 2017). The initial allele frequency also varied among the causative loci. In the initial population, by drawing from a uniform distribution ranging from 0 to 1, we assigned a probability that an individual carries a given allele at the QTL. A draw from a binomial distribution determined whether a given individual harbored the allele at that locus. From this set of QTL, we assumed additivity and calculated a phenotype for each individual based on the initial heritability (ℎ^2^) of the phenotype. In all of our models, we considered initial heritability (ℎ^2^) parameter values of 0.1, 0.5, and 0.8. To generate the phenotype, we first calculated the genetic effect (*G_j_*) for the *j*th individual as 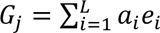, where *a_i_* is the allele call at the *i*th locus, *e_i_* is the effect size at the *i*th locus, and *L* is the number of QTL. To determine the environmental effect (*E_j_*) for each individual, we first calculated the expected environmental variance from the heritability parameter (ℎ^2^) by rearranging the classic equation, 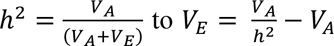, where *V_A_*is the additive genetic variance and *V_E_* is the environmental variance (Sella and Barton 2019; Hayward and Sella 2022). Based on the distributions we used to generate the genetic effect, the expected additive genetic variance is 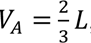, thus we used this value for all simulations to set the environmental variance. We then obtained an environmental effect (*E_j_*) for each individual by drawing random values from a normal distribution with a mean of zero and standard deviation of 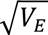. Thus, for the *j*th individual, the raw phenotype value was *P_j_*= *G_j_* + *E_j_*. We did not include a term for genotype by environment interaction in this model, meaning it is not possible for phenotypic plasticity to evolve. Our method of generating effect sizes and a model of additivity means that models with more causative loci will produce phenotypes with higher numerical values. To ease comparison across models and maintain values on the same scale, we scaled all phenotypes by using the expected mean and standard deviation for the population phenotype. For our model parameters above, the expected phenotypic mean is: *μ_p_* = *L*, and the expected phenotypic standard deviation is 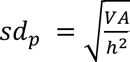. Thus, the scaled phenotype is to 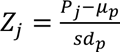, with an expected adjusted mean of 0 and standard deviation of 1 to keep the phenotypes on a similar scale regardless of the number of contributing loci. The structure described above is summarized in Figure 1A.

### Temporal Patterns of Optima and Fitness

We simulated six different types of models that differed in the temporal pattern of the optima over time including a null model, a new constant optimum model, and four different models in which the optima fluctuate over time. These models and the fitness function are summarized in Figure 1B. As a null model, we performed simulations with the same genomic characteristics described above but with individuals assigned a random fitness value drawn from a uniform distribution ranging from 0.1 to 1. For all other models, an individual’s fitness was dependent on the distance between an individual’s phenotype and the optimum at a given timepoint as follows: *W_j_* = 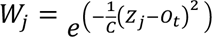, where *W_j_* is the *j*th individual’s fitness, *Z_j_* is the *j*th individual’s scaled phenotype value, *O_t_* is the optimum at time *t,* and *C* is a constant that influences the steepness of the fitness surface. When the optimum shifts, the strength of selection experienced in the population can both be altered by altering how far the optimum moves and/or by altering *C.* Because it is not necessary to change both these factors to change selection strength, and would be unnecessarily complicated to do so, we chose to keep *C* constant throughout all simulations at *C = 125* and only adjusted the distance the optimum shifts as a parameter. We used this value of *C* for convenience in that it allowed us to change the optimum in simple integer values (0, 1, 2, 3, 4) while keeping the selection strength realistic. For context, an individual with a phenotypic value of 0 (i.e. the starting mean population value) would have a fitness of 0.88 when the optimum shifts our highest amount to 4 or a fitness of 0.99 when the optimum shifts to our lowest value of 1 (Figure 1 shows this visually).

#### (i) Constant new optimum

In the model where there is a single new constant optimum, the optimum is *O_t_* = *μ_z_* + Δ*σ_z_*, where *μ_z_* is the mean scaled phenotype in the initial population, *σ*_<_is the standard deviation of the scaled phenotype in the initial population, and Δ is the distance to the new optima. We considered parameter values of 1, 2, 3, and 4 for the Δ parameter, which determined how far away the new optimum was from the initial population mean in units of standard deviations. In this constant model, the optimum shifted immediately at the start of the simulation and was constant for the entire simulation.

#### (ii) Two-season models with fluctuating optima

In our simplest fluctuating optima model, we simulated an optimum that shifted instantaneously between two values, each lasting a season length (*Gen)* in generations. The two optima were set in the same way as the constant model, based on the initial population mean as follows: *O_t_* = *μ_z_* ± Δ*σ_z_*. The optimum switches between the (+) and (-) values, each for a period of *Gen* generations. We considered the following *Gen* parameter values: 10, 20, and 30. In nature, shifts between seasons often occur gradually (Gregory 2009; Oz et al. 2014). Thus, we also considered a two-season model (referred to a simple gradual model) where the shift in the two optima occurred gradually with the optimum at time *t* defined as: 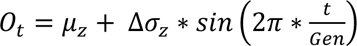.

### Models with complex patterns of optima and season lengths

The above two season models kept the distance to the optima (Δ) and the season length (*Gen*) consistent across the entire simulation to serve as simple, general models. Actual patterns of fluctuating selection in nature are expected to more complex and variable over time. Thus, we simulated two models with additional complexity. These models included a situation in which the season length differed across four time periods (gradual optima change with even optima and uneven season length - referred to as complex gradual I), and a situation in which both the distance to the optima and the season length differed across four time periods (gradual optima change with uneven optima and uneven season length - referred to as complex gradual II). For both models, the optimum at time *t* is defined as: 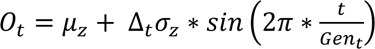. The difference here is that Δ*_t_* and *Gen_t_* vary across time. In the model of gradual optima change with even optima and uneven season length, Δ*_t_* remains constant throughout the simulation and we consider the same parameter values of either 1, 2, 3, or 4. The *Gen_t_* values follow a pattern of four time periods that then repeat of 12, 22, 10, 16. Within each of these four time periods, there is a single sinusoidal cycle, for example, the 12 generation period consists of 6 generations of a high optimum and six of a low optimum. In the model of gradual optima change with uneven optima and uneven season length, the *Gen_t_* parameter is set in the exact same way but the optimum also shifts in each of the four time periods in the following order: 3, 1, 4, 2. These models help understand complex evolutionary scenarios in which a population may experience a gradual change in optima with multiple sub-seasons, each resulting in a different number of generations and corresponding selection pressure, a typical situation for multivoltine organisms that exhibit different generation numbers over the seasons (Mendonça, Jr. and Romanowski 2012; Bjørnstad et al. 2016; Buckley et al. 2017; Harvey et al. 2023; Subedi et al. 2023).

### Modeling fluctuating population size

As natural populations have been shown to fluctuate periodically (Nunez et al. 2024; Nunez et al. 2025), we likewise included additional models to evaluate the effects of fluctuating population size alongside fluctuating selection. The fluctuating population size models followed an instantaneous change, with the population alternating between 1000 individuals for one season and 10000 individuals for the other. In a model involving both fluctuating selection and fluctuating population size, changes occurred in ways that population size and selection were synchronized. The models allowed us to test whether seasonal allele frequency oscillations driven by fluctuating selection remain identifiable when demographic fluctuations generate overlapping temporal changes in allele frequency.

### Exploration of equilibrium dynamics

Because short-term allele frequency oscillations do not necessarily imply the long-term maintenance of genetic variation, we also ran additional simulations designed to assess the long-term evolutionary consequences of fluctuating selection for the following models of instantaneous selection: We ran five replicate simulations for 10Ne generations corresponding to 100,000 generations, where Ne is the effective population size, allowing allele frequencies to approach the mutation-drift equilibrium expectation (Haller and Messer 2023). These models test both the presence and absence of de novo mutation. Models with mutation were run using a mutation rate of 10^-7^ and a recombination rate of 10^-6^ per site per generation, and the interpretation of these models is as a control exploratory simulation rather than a reproduction of a known empirical system.

### Identifying signals of fluctuating selection from allele frequencies and phenotypes

We used ggplot2 (Wickham 2016) and tidyverse packages(Wickham et al. 2019) to visualize changes in allele frequency and the population mean phenotype over time. Furthermore, spectral analysis as a mathematical tool that decomposes a time-domain signal into its frequency-domain representation, it reveals how variance in a time series is distributed across different frequencies (Jenkins 1965; Shumway and Stoffer 2017). Although it may not identify the underlying biological background, spectral analysis can detect periodic components and their strength in a well sampled timeseries. Consequently, we used spectral analysis to decompose the allele frequency signal and identify seasonality (selection length), which enabled us to detect fluctuating selection in our dataset assuming the cyclic selection. Specifically, we first averaged the allele frequencies across all loci to obtain a single mean frequency per generation. We then organized these mean values into time-series, with each generation corresponding to a time point, to illustrate changes in allele frequency over time. Finally, we estimated the spectral density of each time series using the spectrum function from base R (R Core Team 2022 Apr) and applied a smoothing span of 2 to reduce noise and reveal the dominant periodic components. This approach enabled us to detect and characterize cyclical patterns in allele frequencies across generations.

## Data availability

The code used to generate data during simulation and for the analysis is available on GitHub repository: https://github.com/EGKingLab/evogen-sims

## Acknowledgments

We thank the University of Missouri’s ITRSS group for providing the computing resources to perform the simulations in this paper. Dr. Kevin Middleton, Dr. David Kang, Associate Editor Dr. Kirk Lohmueller, two anonymous reviewers provided many helpful comments that improved this work, and King lab members for their helpful feedback.

## References

Abdul-Rahman F, Tranchina D, Gresham D. 2021. Fluctuating Environments Maintain Genetic Diversity through Neutral Fitness Effects and Balancing Selection Townsend J, editor. Mol Biol Evol. 38(10):4362–4375 https://academic.oup.com/mbe/article/38/10/4362/6300528. 10.1093/molbev/msab173

Agashe D, Sane M, Singhal S. 2023. Revisiting the Role of Genetic Variation in Adaptation. Am Nat. 202(4):486–502 [accessed 2023 Oct 25]. https://www.journals.uchicago.edu/doi/10.1086/726012. 10.1086/726012

Arnab SP, Amin MR, DeGiorgio M. 2023. Uncovering Footprints of Natural Selection Through Spectral Analysis of Genomic Summary Statistics Kim Y, editor. Mol Biol Evol. 40(7) [accessed 2025 Sep 1]. https://academic.oup.com/mbe/article/doi/10.1093/molbev/msad157/7222828. 10.1093/molbev/msad157

Ascensao JA, Lok K, Hallatschek O. 2024. Asynchronous abundance fluctuations can drive giant genotype frequency fluctuations. Nature Ecology & Evolution 2024 9:1. 9(*1*):166–179 [accessed 2026 May 28]. https://www.nature.com/articles/s41559-024-02578-3. 10.1038/s41559-024-02578-3

Barghi N et al. 2019. Genetic redundancy fuels polygenic adaptation in Drosophila. PLoS Biol. 17(2). 10.1371/journal.pbio.3000128

Barton NH, Etheridge AM, Véber A. 2017. The infinitesimal model: Definition, derivation, and implications. Theor Popul Biol. 118:50–73 [accessed 2024 Feb 21]. http://creativecommons.org/licenses/by/4.0/. 10.1016/j.tpb.2017.06.001

Barton NH, Turelli M. 1989. Evolutionary quantitative genetics: How little do we know? Annu Rev Genet. 23(Volume 23,):337–370 [accessed 2024 Jul 25]. 10.1146/annurev.ge.23.120189.002005

Behrman EL et al. 2018. Rapid seasonal evolution in innate immunity of wild Drosophila melanogaster. Proceedings of the Royal Society B: Biological Sciences. 285(1870) [accessed 2024 Jan 18]. https://royalsocietypublishing.org/doi/10.1098/rspb.2017.2599. 10.1098/rspb.2017.2599

Bell G. 2010. Fluctuating selection: the perpetual renewal of adaptation in variable environments. Philosophical Transactions of the Royal Society B: Biological Sciences. 365(1537):87–97 [accessed 2023 Oct 30]. https://royalsocietypublishing.org/doi/10.1098/rstb.2009.0150. 10.1098/rstb.2009.0150

Bergland AO et al. 2014. Genomic Evidence of Rapid and Stable Adaptive Oscillations over Seasonal Time Scales in Drosophila. PLoS Genet. 10(11):1004775 [accessed 2023 Jul 11]. www.plosgenetics.org. 10.1371/journal.pgen.1004775

Bertram J, Masel J. 2019. Different mechanisms drive the maintenance of polymorphism at loci subject to strong versus weak fluctuating selection. Evolution (N Y). 73(5):883–896. 10.1111/evo.13719

Bijma P. 2011. A general definition of the heritable variation that determines the potential of a population to respond to selection. Genetics. 189(4):1347–1359 [accessed 2024 Jun 29]. https://dx.doi.org/10.1534/genetics.111.130617. 10.1534/genetics.111.130617

Bitter MC et al. 2024. Continuously fluctuating selection reveals fine granularity of adaptation. Nature 2024 634:8033. 634(8033):389–396 [accessed 2025 Feb 22]. https://www.nature.com/articles/s41586-024-07834-x. 10.1038/s41586-024-07834-x

Bjørnstad ON, Nelson WA, Tobin PC. 2016. Developmental synchrony in multivoltine insects: generation separation versus smearing. Popul Ecol. 58(4):479–491. 10.1007/s10144-016-0564-z

Brud E. 2025. Season-specific dominance broadly stabilizes polymorphism under symmetric and asymmetric multivoltinism. Genetics. 229(4):28 [accessed 2026 Jun 24]. https://dx.doi.org/10.1093/genetics/iyaf028. 10.1093/GENETICS/IYAF028

Buckley LB et al. 2017. Insect development, thermal plasticity and fitness implications in changing, seasonal environments. Integr Comp Biol. 57(5):988–998. 10.1093/icb/icx032

Bürger R, Ghnelfarb A. 1999. Genetic Variation Maintained in Multilocus Models of Additive Quantitative Traits Under Stabilizing Selection. Genetics. 152(2):807–820 [accessed 2024 Sep 26]. https://dx.doi.org/10.1093/genetics/152.2.807. 10.1093/GENETICS/152.2.807

Burke MK et al. 2010. Genome-wide analysis of a long-term evolution experiment with Drosophila. Nature. 467(7315):587–590 [accessed 2024 Jan 14]. https://www.nature.com/articles/nature09352. 10.1038/nature09352

Van Buskirk J, Smith DC. 2021. Ecological causes of fluctuating natural selection on habitat choice in an amphibian. Evolution (N Y). 75(7):1862–1877 https://academic.oup.com/evolut/article/75/7/1862/6728832. 10.1111/evo.14282

Campbell-Staton SC et al. 2017. Winter storms drive rapid phenotypic, regulatory, and genomic shifts in the green anole lizard. Science (1979). 357(6350):495–498 [accessed 2024 Jan 14]. https://www.science.org/doi/10.1126/science.aam5512. 10.1126/science.aam5512

Cvijović I, Good BH, Jerison ER, Desai MM. 2015. Fate of a mutation in a fluctuating environment. Proceedings of the National Academy of Sciences. 112(36):E5021–E5028 [accessed 2024 Jul 26]. https://pnas.org/doi/full/10.1073/pnas.1505406112. 10.1073/pnas.1505406112

Doebley JF, Gaut BS, Smith BD. 2006. The Molecular Genetics of Crop Domestication. Cell. 127(7):1309–1321 [accessed 2024 Jan 14]. https://linkinghub.elsevier.com/retrieve/pii/S0092867406015923. 10.1016/j.cell.2006.12.006

Dumont BL, Payseur BA. 2008. Evolution of the gnomic rate of recombination in mammals. Evolution (N Y). 62(2):276–294 [accessed 2025 May 6]. https://dx.doi.org/10.1111/j.1558-5646.2007.00278.x. 10.1111/J.1558-5646.2007.00278.X

Ellner S, Hairston Jnr NG. 1994. Role of overlapping generations in maintaining genetic variation in a fluctuating environment. Am Nat. 143(3):403–417 [accessed 2024 Jul 31]. https://www.journals.uchicago.edu/doi/10.1086/285610. 10.1086/285610

Endler JA. 2020. Natural Selection in the Wild. (MPB-21), Volume 21. Princeton University Press. [accessed 2025 Feb 7]. http://www.jstor.org/stable/10.2307/j.ctvx5w9v9. 10.2307/j.ctvx5w9v9

Gienapp P, Reed TE, Visser ME. 2014. Why climate change will invariably alter selection pressures on phenology. Proceedings of the Royal Society B: Biological Sciences. 281(1793):20141611 [accessed 2025 Feb 7]. https://royalsocietypublishing.org/doi/10.1098/rspb.2014.1611. 10.1098/rspb.2014.1611

Gillespie J. 1973. Polymorphism in random environments. Theor Popul Biol. 4(2):193–195 [accessed 2024 Jun 29]. 10.1016/0040-5809(73)90028-2

Gillespie JH. 1977. Natural Selection for Variances in Offspring Numbers: A New Evolutionary Principle. Source: The American Naturalist. 111(981):1010–1014 [accessed 2024 Sep 25]

Glaser-Schmitt A, Wittmann MJ, Ramnarine TJS, Parsch J. 2021. Sexual Antagonism, Temporally Fluctuating Selection, and Variable Dominance Affect a Regulatory Polymorphism in *Drosophila melanogaster* Larracuente A, editor. Mol Biol Evol. 38(11):4891–4907 [accessed 2024 Jun 29]. https://academic.oup.com/mbe/article/38/11/4891/6325090. 10.1093/molbev/msab215

Gloss AD, Whiteman NK. 2016. Balancing Selection: Walking a Tightrope. Current Biology. 26(2):R73–R76. 10.1016/j.cub.2015.11.023

Gossmann TI, Waxman D, Eyre-Walker A. 2014. Fluctuating selection models and Mcdonald-Kreitman type analyses. PLoS One. 9(1). 10.1371/journal.pone.0084540

Grant PR, Grant BR. 2002. Unpredictable evolution in a 30-year study of Darwin’s finches. Science (1979). 296(5568):707–711 [accessed 2024 Jul 25]. https://www.science.org/doi/10.1126/science.1070315. 10.1126/science.1070315

Greener JG, Kandathil SM, Moffat L, Jones DT. 2021. A guide to machine learning for biologists. Nature Reviews Molecular Cell Biology 2021 23:1. 23(1):40–55 [accessed 2024 Dec 9]. https://www.nature.com/articles/s41580-021-00407-0. 10.1038/s41580-021-00407-0

Gregory TR. 2009. Understanding Natural Selection: Essential Concepts and Common Misconceptions. Evolution: Education and Outreach. 2(2):156–175 [accessed 2024 Feb 22]. https://evolution-outreach.biomedcentral.com/articles/10.1007/s12052-009-0128-1. 10.1007/s12052-009-0128-1

Haasl RJ, Payseur BA. 2016. Fifteen years of genomewide scans for selection: trends, lessons and unaddressed genetic sources of complication. Mol Ecol. 25(1):5–23 [accessed 2025 Sep 1]. /doi/pdf/10.1111/mec.13339. 10.1111/MEC.13339

Haller BC, Messer PW. 2023. SLiM 4: Multispecies Eco-Evolutionary Modeling. American Naturalist. 201(5). 10.1086/723601

Hancock AM, Di Rienzo A. 2008. Detecting the Genetic Signature of Natural Selection in Human Populations: Models, Methods, and Data. Annu Rev Anthropol. 37:197 [accessed 2023 Mar 12]. /pmc/articles/PMC2901121/. 10.1146/ANNUREV.ANTHRO.37.081407.085141

Harvey JA et al. 2023. Scientists’ warning on climate change and insects. Ecol Monogr. 93(1). 10.1002/ecm.1553

Hayward LK, Sella G. 2022. Polygenic adaptation after a sudden change in environment. Elife. 11 [accessed 2023 Oct 30]. https://elifesciences.org/articles/66697. 10.7554/eLife.66697

Hernández-Carrasco D, Tylianakis JM, Lytle DA, Tonkin JD. 2025. Ecological and evolutionary consequences of changing seasonality. Science (1979). 388(6750) [accessed 2025 Jul 14]. https://www.science.org/doi/10.1126/science.ads4880. 10.1126/science.ads4880

Hill WG, Mackay TFC. 2004. D. S. Falconer and Introduction to Quantitative Genetics. Genetics. 167(4):1529–1536 https://academic.oup.com/genetics/article/167/4/1529/6050406. 10.1093/genetics/167.4.1529

Hoekstra RF, Bijlsma R, Dolman AJ. 1985. Polymorphism from environmental heterogeneity: models are only robust if the heterozygote is close in fitness to the favoured homozygote in each environment. Genet Res (Camb). 45(3):299–314 [accessed 2025 Feb 25]. https://www.cambridge.org/core/journals/genetics-research/article/polymorphism-from-environmental-heterogeneity-models-are-only-robust-if-the-heterozygote-is-close-in-fitness-to-the-favoured-homozygote-in-each-environment/E533B7611E0C6100C367A8E84A2D3E37. 10.1017/S001667230002228X

Holstad A et al. 2024. Evolvability predicts macroevolution under fluctuating selection. Science (1979). 384(6696):688–693 [accessed 2024 Jun 29]. https://www.science.org/doi/10.1126/science.adi8722. 10.1126/science.adi8722

Huerta-Sanchez E, Durrett R, Bustamante CD. 2008. Population Genetics of Polymorphism and Divergence Under Fluctuating Selection. Genetics. 178(1):325–337. 10.1534/genetics.107.073361

Jenkins GM. 1965. A Survey of Spectral Analysis. Appl Stat. 14(1):2 [accessed 2024 Jul 31]. 10.2307/2985352

Johnson OL, Tobler R, Schmidt JM, Huber CD. 2026. Genetic Footprints of Seasonal Fluctuating Selection: A Comparison With Established Selection Forms. Genome Biol Evol. 18(4) [accessed 2026 Apr 24]. https://doi.org/10.1093/gbe/evag082. 10.1093/GBE/EVAG082

Kelly JK. 2022. The genomic scale of fluctuating selection in a natural plant population. Evol Lett. 6(6):506–521. 10.1002/evl3.308

Khan MA, Pierre JW. 2019. Detection of Periodic Forced Oscillations in Power Systems Using Multitaper Approach. IEEE Transactions on Power Systems. 34(2):1086–1094 [accessed 2025 Aug 25]. 10.1109/TPWRS.2018.2870838

Kim Y. 2023. Partial protection from fluctuating selection leads to evolution towards wider population size fluctuation and a novel mechanism of balancing selection. Proceedings of the Royal Society B: Biological Sciences. 290(2001). 10.1098/rspb.2023.0822

Lande R. 2007. Expected relative fitness and the adaptive topography of fluctuating selection. Evolution (N Y). 61(8):1835–1846 [accessed 2023 Jun 14]. 10.1111/J.1558-5646.2007.00170.X

Lande R. 2009. Adaptation to an extraordinary environment by evolution of phenotypic plasticity and genetic assimilation. J Evol Biol. 22(7):1435–1446 [accessed 2023 Oct 30]. 10.1111/J.1420-9101.2009.01754.X

Lande R. 2014. Evolution of phenotypic plasticity and environmental tolerance of a labile quantitative character in a fluctuating environment. J Evol Biol. 27(5):866–875 [accessed 2025 May 6]. https://doi.org/10.1111/jeb.12360. 10.1111/JEB.12360

Louthan AM, Kay KM. 2011. Comparing the adaptive landscape across trait types: larger QTL effect size in traits under biotic selection. BMC Evol Biol. 11(1):60 [accessed 2024 Jun 25]. http://bmcevolbiol.biomedcentral.com/articles/10.1186/1471-2148-11-60. 10.1186/1471-2148-11-60

Lynch M, Wei W, Ye Z, Pfrender M. 2024. The genome-wide signature of short-term temporal selection. Proceedings of the National Academy of Sciences. 121(28):e2307107121 [accessed 2025 Feb 5]. https://pnas.org/doi/10.1073/pnas.2307107121. 10.1073/pnas.2307107121

Machado HE et al. 2021. Broad geographic sampling reveals the shared basis and environmental correlates of seasonal adaptation in Drosophila. Elife. 10 [accessed 2023 Nov 5]. https://elifesciences.org/articles/67577. 10.7554/eLife.67577

McAdam AG, Boutin S, Dantzer B, Lane JE. 2019. Seed Masting Causes Fluctuations in Optimum Litter Size and Lag Load in a Seed Predator. Am Nat. 194(4):574–589 [accessed 2024 Jun 24]. https://www.journals.uchicago.edu/doi/10.1086/703743. 10.1086/703743

Mendonça, Jr. M de S, Romanowski HP. 2012. Population ecology of the multivoltine Neotropical gall midge Eugeniamyia dispar (Diptera, Cecidomyiidae). Iheringia Ser Zool. 102(2):170–176 [accessed 2024 Dec 15]. http://www.scielo.br/scielo.php?script=sci_arttext&pid=S0073-47212012000200009&lng=en&tlng=en. 10.1590/S0073-47212012000200009

Miura S, Zhang Z, Nei M. 2013. Random fluctuation of selection coefficients and the extent of nucleotide variation in human populations. Proc Natl Acad Sci U S A. 110(26):10676–10681. 10.1073/pnas.1308462110

Nunez JCB et al. 2024. A cosmopolitan inversion facilitates seasonal adaptation in overwintering Drosophila. Genetics. 226(2). 10.1093/genetics/iyad207

Nunez JCB et al. 2025. Footprints of Worldwide Adaptation in Structured Populations of Drosophila melanogaster Through the Expanded DEST 2.0 Genomic Resource. Mol Biol Evol. 42(8):1–28 [accessed 2025 Nov 12]. https://doi.org/10.1093/molbev/msaf132. 10.1093/MOLBEV/MSAF132

Oostra V, Saastamoinen M, Zwaan BJ, Wheat CW. 2018. Strong phenotypic plasticity limits potential for evolutionary responses to climate change. Nature Communications 2018 9:1. 9(1):1–11 [accessed 2025 May 6]. https://www.nature.com/articles/s41467-018-03384-9. 10.1038/s41467-018-03384-9

Oz T et al. 2014. Strength of Selection Pressure Is an Important Parameter Contributing to the Complexity of Antibiotic Resistance Evolution. Mol Biol Evol. 31(9):2387–2401 [accessed 2024 Feb 22]. https://academic.oup.com/mbe/article-lookup/doi/10.1093/molbev/msu191. 10.1093/molbev/msu191

Park JS. 2019. Cyclical environments drive variation in life-history strategies: a general theory of cyclical phenology. Proceedings of the Royal Society B: Biological Sciences. 286(1898):20190214 [accessed 2025 Aug 17]. https://royalsocietypublishing.org/doi/10.1098/rspb.2019.0214. 10.1098/rspb.2019.0214

Parzen E. 1967. The Role of Spectral Analysis in Time Series Analysis. Revue de l’Institut International de Statistique / Review of the International Statistical Institute. 35(2):125 [accessed 2024 Jul 31]. 10.2307/1401395

Pfenninger M, Foucault Q. 2022. Population Genomic Time Series Data of a Natural Population Suggests Adaptive Tracking of Fluctuating Environmental Changes. Integr Comp Biol. 62(6):1812–1826 https://academic.oup.com/icb/article/62/6/1812/6619072. 10.1093/icb/icac098

Phillips MA, Kutch IC, Long AD, Burke MK. 2020. Increased time sampling in an evolve-and-resequence experiment with outcrossing Saccharomyces cerevisiae reveals multiple paths of adaptive change. Mol Ecol. 29(24):4898–4912 [accessed 2023 Mar 12]. https://onlinelibrary.wiley.com/doi/full/10.1111/mec.15687. 10.1111/MEC.15687

Platt T, Denman KL. 1975. Spectral Analysis in Ecology. Source: Annual Review of Ecology and Systematics. 6:189–210 [accessed 2023 Oct 25]. https://www.jstor.org/stable/2096830

Qin X, Chiang CWK, Gaggiotti OE. 2022. Deciphering signatures of natural selection via deep learning. Brief Bioinform. 23(5):1–10 [accessed 2025 Sep 1]. https://doi.org/10.1093/bib/bbac354. 10.1093/BIB/BBAC354

R Core Team. 2022. [accessed 2025 Feb 14]. https://www.r-project.org/

Roberts Kingman GA et al. 2021. Predicting future from past: The genomic basis of recurrent and rapid stickleback evolution. Sci Adv. 7(25):5285–5303 [accessed 2023 Nov 1]. https://www.science.org/doi/10.1126/sciadv.abg5285. 10.1126/sciadv.abg5285

Roff DA. 1997. Phenotypic Plasticity and Reaction Norms. Evolutionary Quantitative Genetics. 196–240 [accessed 2025 May 5]. https://link.springer.com/chapter/10.1007/978-1-4615-4080-9_6. 10.1007/978-1-4615-4080-9_6

Rudman SM et al. 2022. Direct observation of adaptive tracking on ecological time scales in Drosophila. Science (1979). 375(6586). 10.1126/science.abj7484

Rusuwa BB et al. 2022. Natural variation at a single gene generates sexual antagonism across fitness components in Drosophila. Current Biology. 32(14):3161–3169.e7 [accessed 2024 Jun 24]. https://linkinghub.elsevier.com/retrieve/pii/S0960982222008351. 10.1016/j.cub.2022.05.038

Sella G, Barton NH. 2019. Thinking About the Evolution of Complex Traits in the Era of Genome-Wide Association Studies. Annu Rev Genomics Hum Genet. 20(1):461–493 [accessed 2023 Nov 1]. https://www.annualreviews.org/doi/10.1146/annurev-genom-083115-022316. 10.1146/annurev-genom-083115-022316

Shumway RH, Stoffer DS. 2017. Time Series Analysis and Its Applications. Springer International Publishing. (Springer Texts in Statistics). [accessed 2025 Aug 19]. https://link.springer.com/10.1007/978-3-319-52452-8. 10.1007/978-3-319-52452-8

Subedi B, Poudel A, Aryal S. 2023. The impact of climate change on insect pest biology and ecology: Implications for pest management strategies, crop production, and food security. J Agric Food Res. 14:100733 https://linkinghub.elsevier.com/retrieve/pii/S2666154323002405. 10.1016/j.jafr.2023.100733

Tenaillon O et al. 2016. Tempo and mode of genome evolution in a 50,000-generation experiment. Nature. 536(7615):165–170 [accessed 2024 Jan 14]. https://www.nature.com/articles/nature18959. 10.1038/nature18959

Thomas E Reed, Daniel E Schindler, Robin S Waples. 2011. Interacting Effects of Phenotypic Plasticity and Evolution on Population Persistence in a Changing Climate. Conservation Biology. 25(1):56–63 [accessed 2023 Oct 30]. https://onlinelibrary.wiley.com/doi/10.1111/j.1523-1739.2010.01552.x. 10.1111/j.1523-1739.2010.01552.x

Trut L, Oskina I, Kharlamova A. 2009. Animal evolution during domestication: the domesticated fox as a model. BioEssays. 31(3):349–360 [accessed 2024 Jan 14]. https://onlinelibrary.wiley.com/doi/10.1002/bies.200800070. 10.1002/bies.200800070

University of Missouri Cluster (Hellbender). [accessed 2026 Jun 29]. https://hdl.handle.net/10355/97710. 10.32469/10355/97710

Via S, Lande R. 1985. Genotype-Environment Interaction and the Evolution of Phenotypic Plasticity. Evolution (N Y). 39(3):505 [accessed 2025 May 6]. https://www.jstor.org/stable/2408649?origin=crossref. 10.2307/2408649

de Vladar HP, Barton N. 2014. Stability and Response of Polygenic Traits to Stabilizing Selection and Mutation. Genetics. 197(2):749–767 [accessed 2024 Jul 29]. https://academic.oup.com/genetics/article/197/2/749/6074097. 10.1534/genetics.113.159111

Waldvogel A et al. 2018. The genomic footprint of climate adaptation in *Chironomus riparius*. Mol Ecol. 27(6):1439–1456 [accessed 2024 Jun 24]. https://onlinelibrary.wiley.com/doi/10.1111/mec.14543. 10.1111/mec.14543

Walsh B, Lynch M. 2018. Evolution and Selection of Quantitative Traits. Vol 1 Oxford University Press. https://academic.oup.com/book/40062. 10.1093/oso/9780198830870.001.0001

West R, Mobilia M. 2020. Fixation properties of rock-paper-scissors games in fluctuating populations. J Theor Biol. 491:110135 [accessed 2024 Jul 26]. https://linkinghub.elsevier.com/retrieve/pii/S0022519319305041. 10.1016/j.jtbi.2019.110135

Wickham H. 2016. ggplot2. (Use R!). [accessed 2024 Feb 25]. http://link.springer.com/10.1007/978-3-319-24277-4. 10.1007/978-3-319-24277-4

Wickham H et al. 2019. Welcome to the Tidyverse. J Open Source Softw. 4(43):1686 [accessed 2026 Jun 29]. https://joss.theoj.org/papers/10.21105/joss.01686. 10.21105/JOSS.01686

Wittmann MJ et al. 2017. Seasonally fluctuating selection can maintain polymorphism at many loci via segregation lift. Proceedings of the National Academy of Sciences. 114(46):E9932–E9941 https://pnas.org/doi/full/10.1073/pnas.1702994114. 10.1073/pnas.1702994114

Wittmann MJ, Mousset S, Hermisson J. 2023. Modeling the genetic footprint of fluctuating balancing selection: from the local to the genomic scale. Genetics. 223(4) [accessed 2024 Jan 10]. https://doi.org/10.1093/genetics/iyad022. 10.1093/GENETICS/IYAD022

Yair S, Coop G. 2022. Population differentiation of polygenic score predictions under stabilizing selection. Philosophical Transactions of the Royal Society B: Biological Sciences. 377(1852) https://royalsocietypublishing.org/doi/10.1098/rstb.2020.0416. 10.1098/rstb.2020.0416

