## Supplemental figures for "Dissecting fluctuating selection: A unified population and quantitative genetics framework"

E.T

### 1. Null model

#### 1.1. Null selection Allele frequency

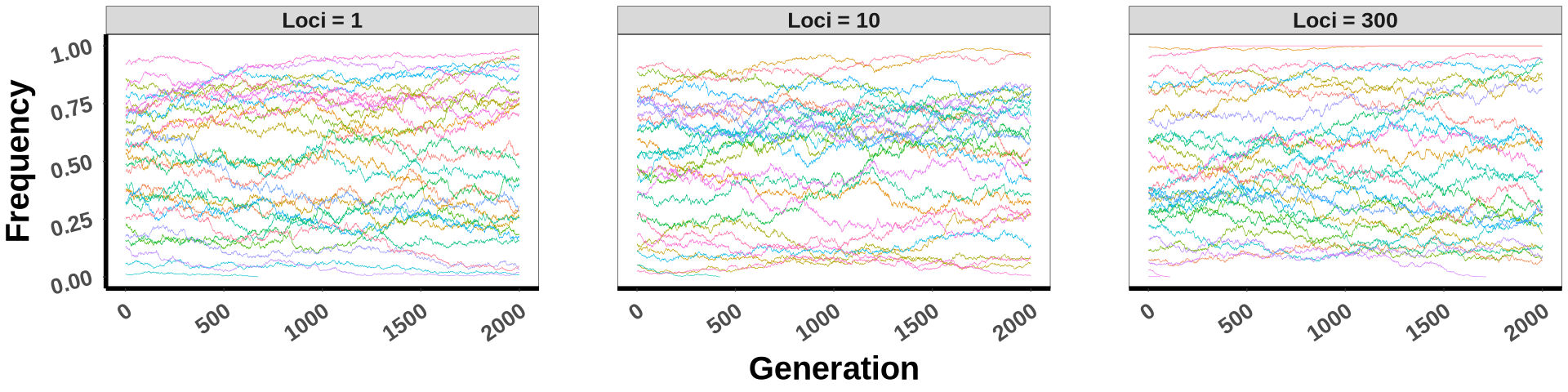

SI 1.1. Allele frequency dynamics for monogenic, oligogec (10 loci), and polygenic (300 loci) traits models. Each line represent a single locus sampled from all loci within a single replicate of each model.

#### 1.2. Null selection phenotypes

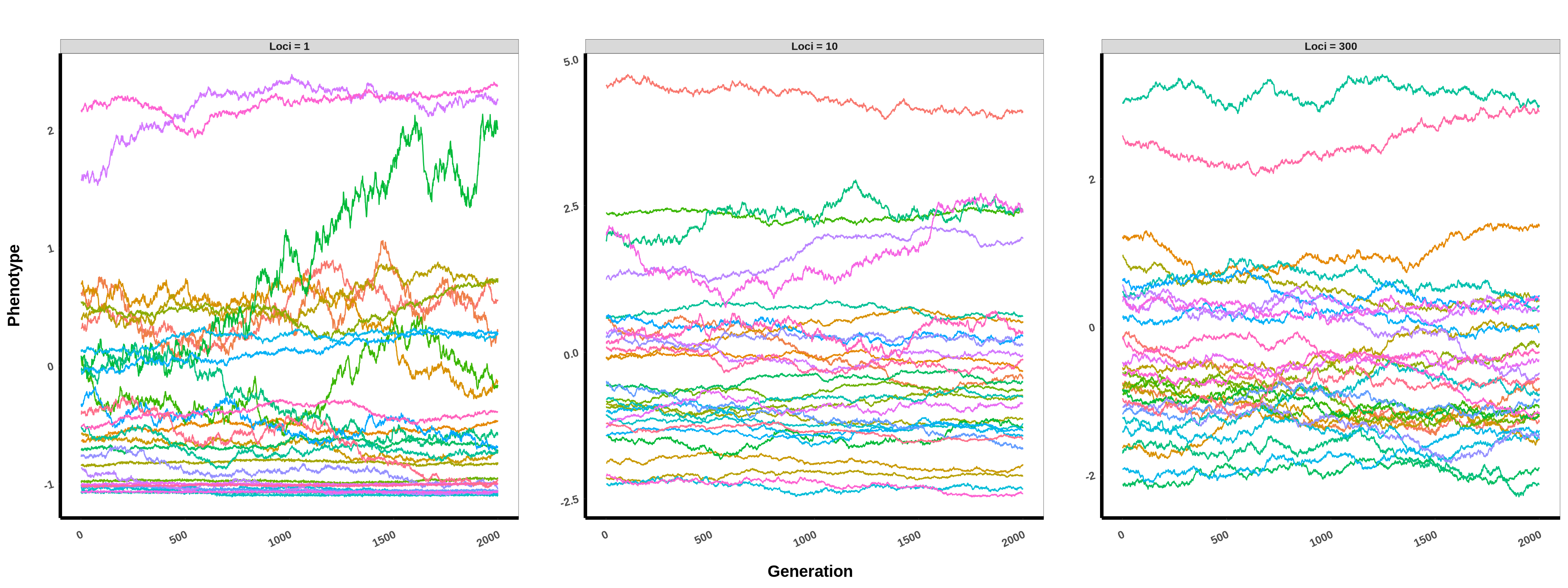

SI 1.2. Mean population phenotypic plot across 30 replicates (each line represents a single replicate) for Null selection, showing parameter interactions.

#### 1.3. Null selection spectral analysis

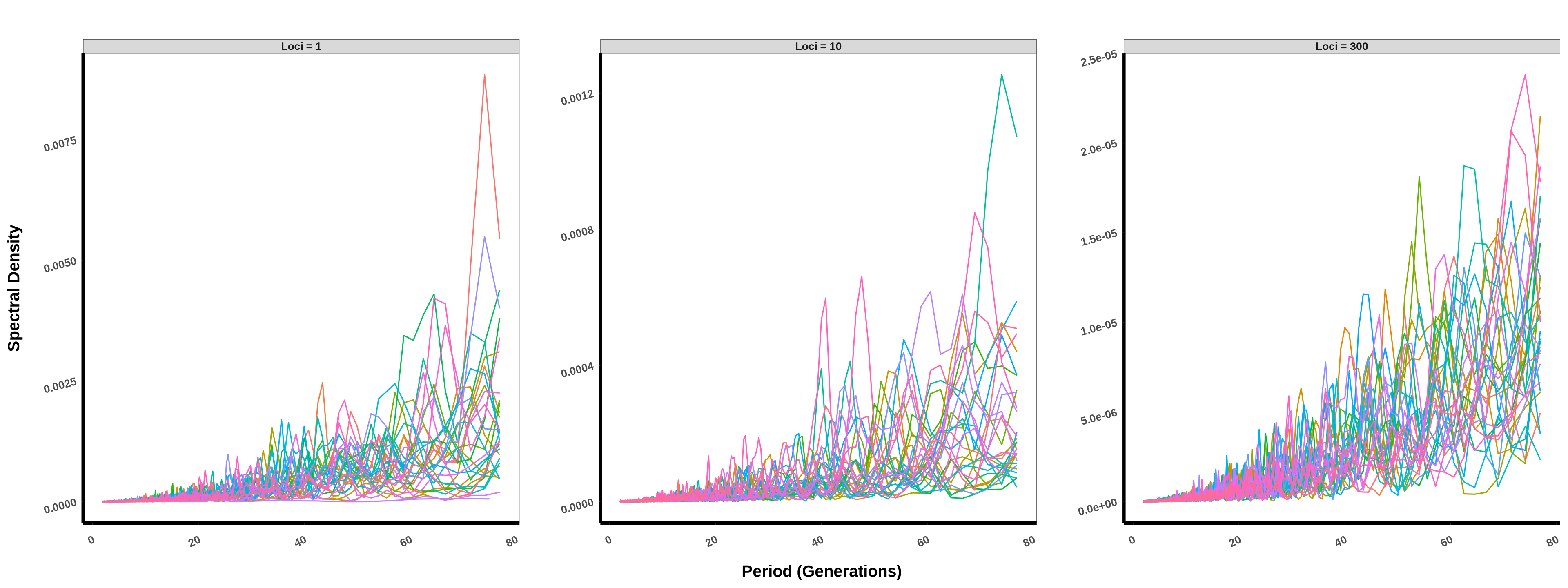

SI 1.3. Spectral density across 30 replicates for all genetic architecture under Null selection model. The density plots do not show any peak as selection is purely driven by random drift.

### 2. Constant model

#### 2.1. Allele frequency all parameters for a single replicate

##### 2.1.1. Monogenic

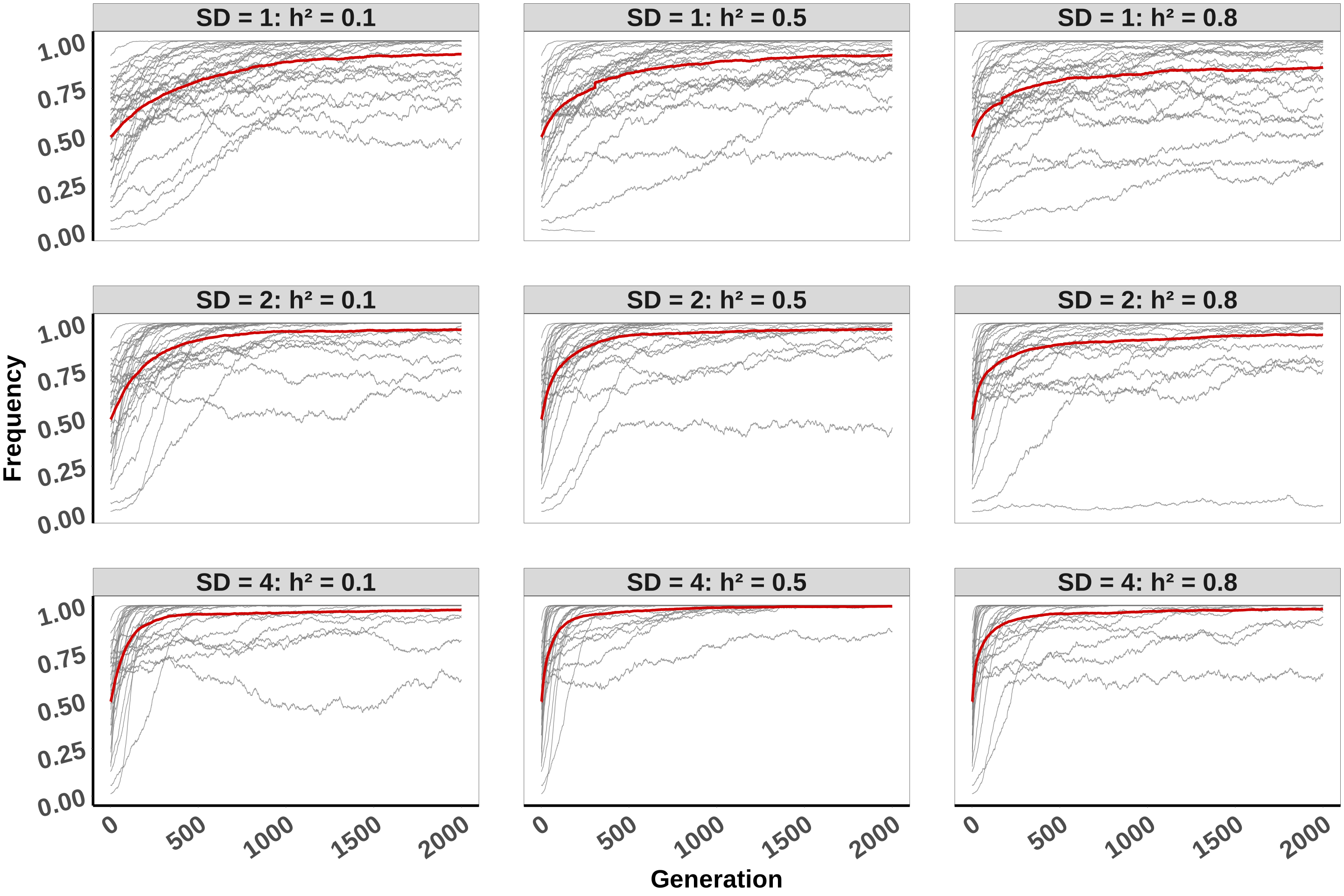

SI 2.1.1. Single locus 30 replicates allele frequency dynamics for constant selection model, showing all parameter interactions. The highlighted line indicates the mean allele frequency for each generation.

##### 2.1.2. Oligogenic

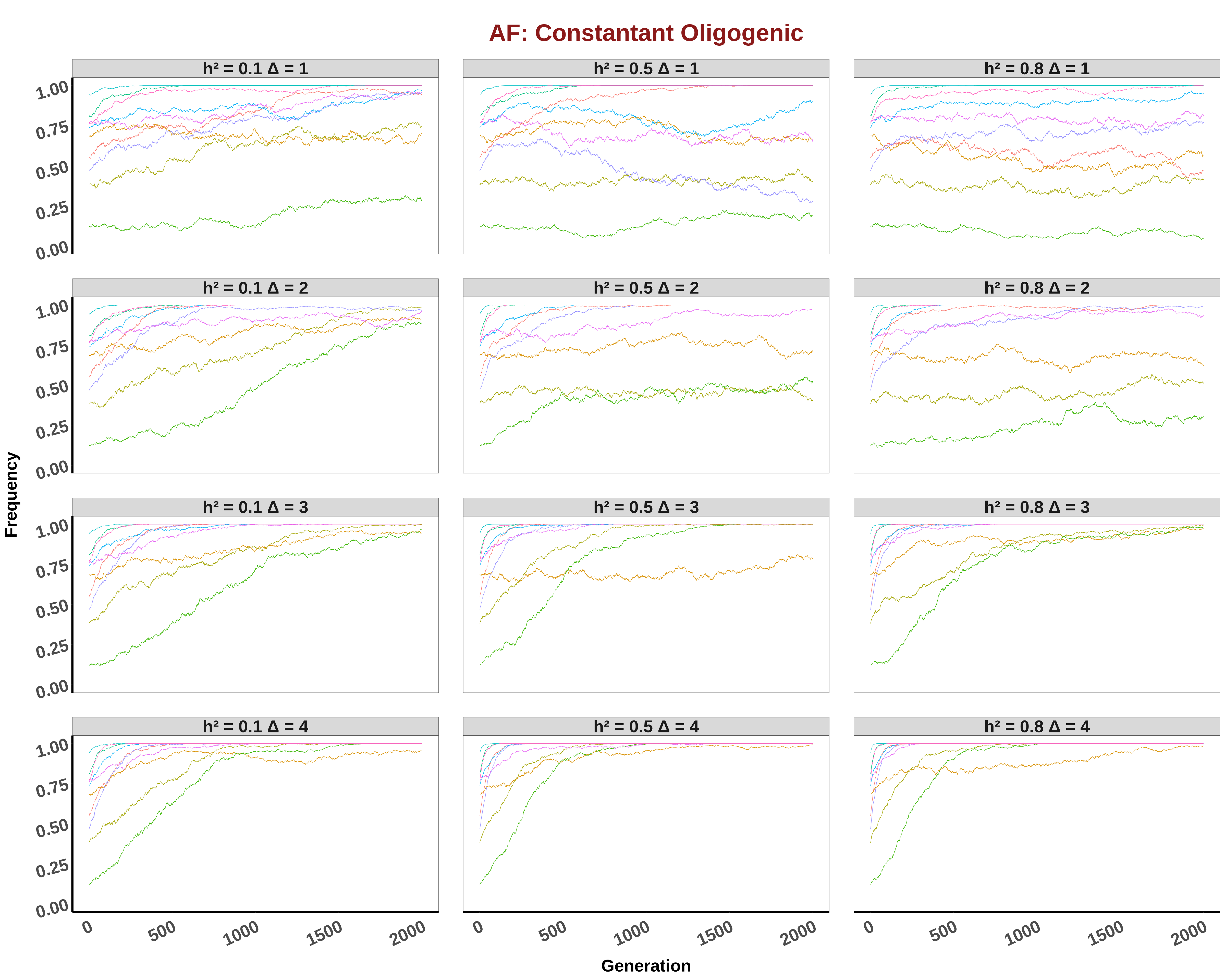

SI 2.1.2. Allele frequency dynamics for all parameter combinations from a single replicate, constant selection model. Each line represents the frequency trajectory for a locus from oligogenic trait (10 loci).

##### 2.1.3. Polygenic 70 loci

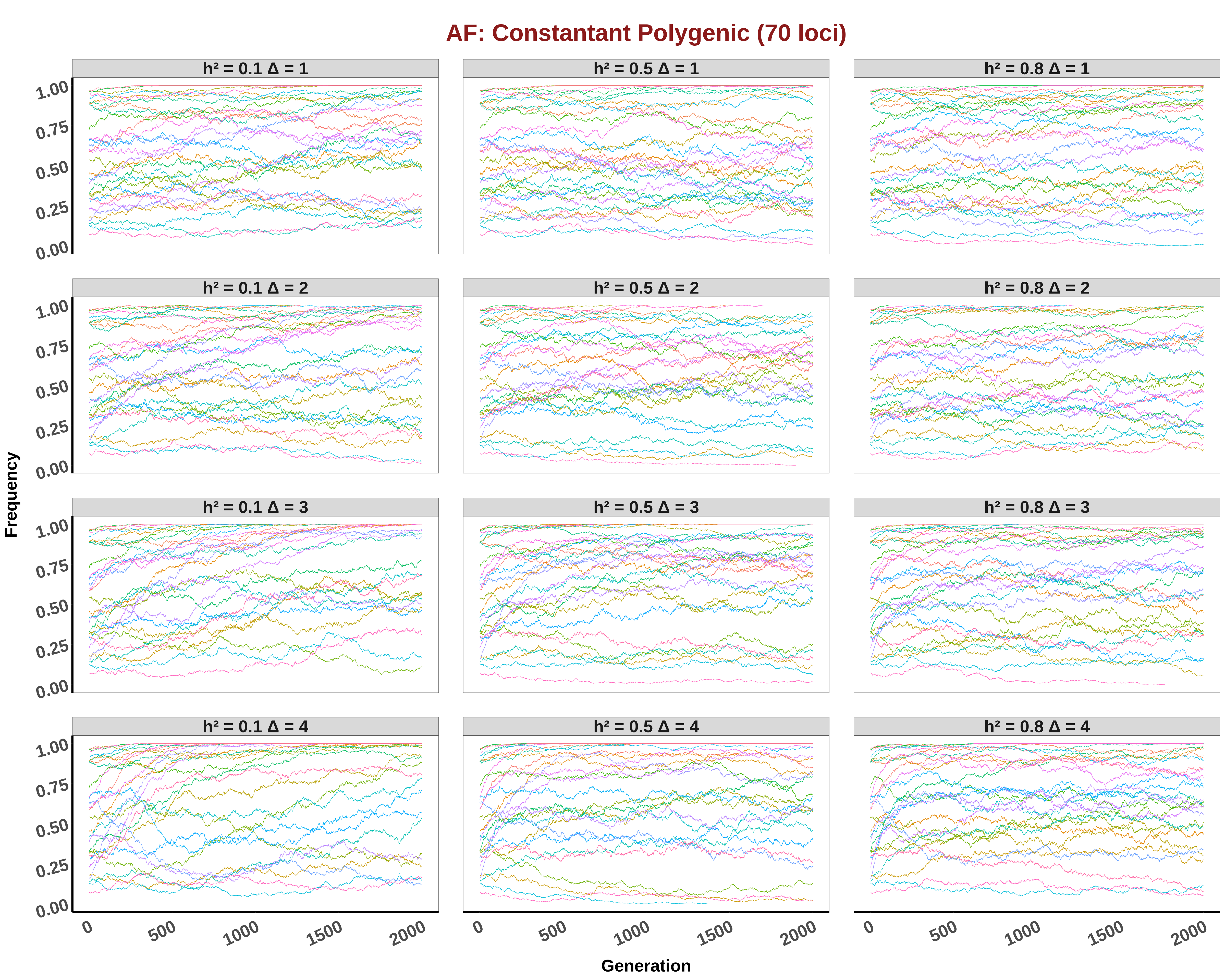

SI 2.1.3: Allele frequency dynamics for all parameter combinations from a single replicate, constant selection model. Each line represents the frequency trajectory for a locus of 30 sampled loci from polygenic trait (70 loci).

##### 2.1.4. Polygenic 300 loci

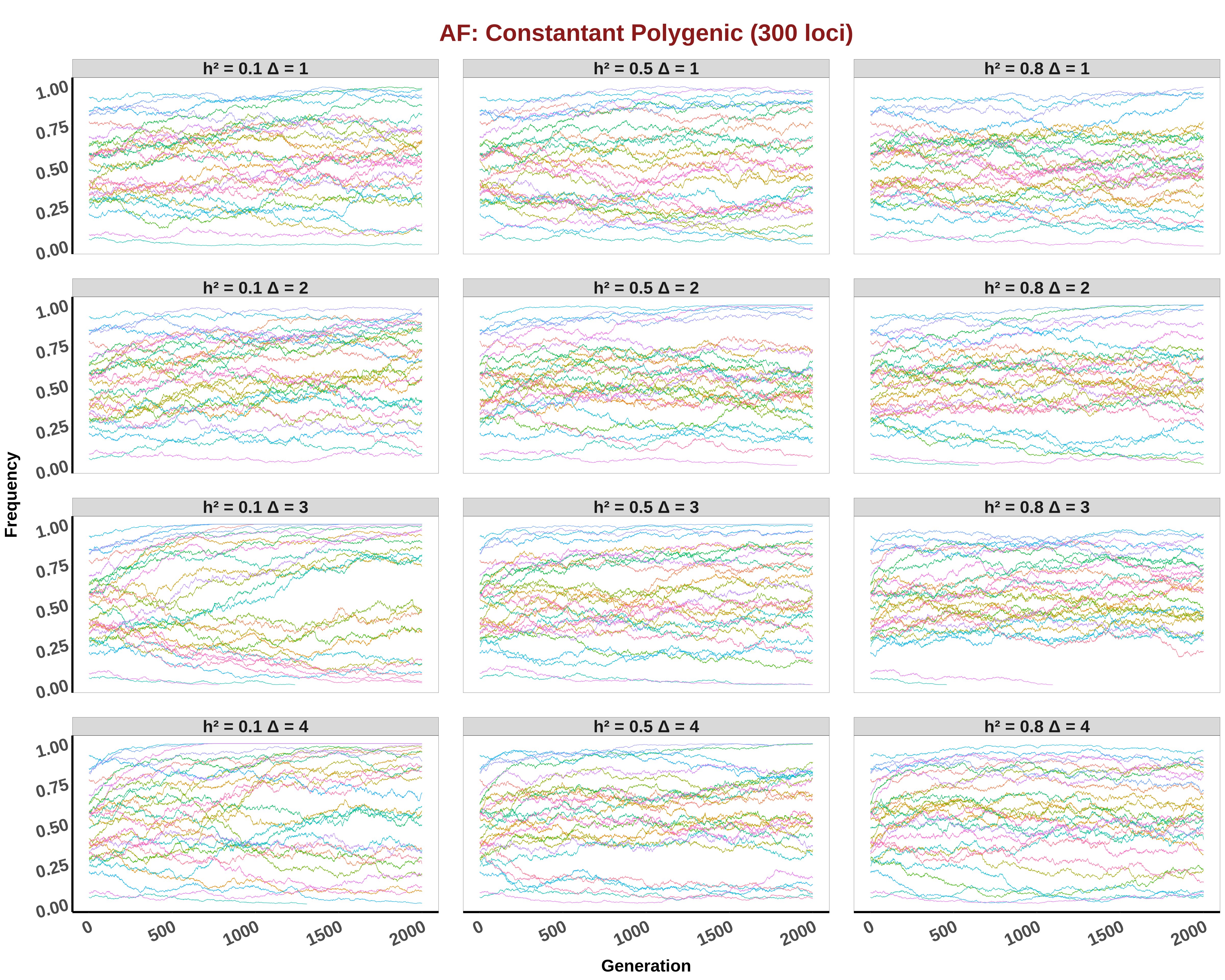

SI 2.1.4. Allele frequency dynamics for all parameter combinations from a single replicate, constant selection model. Each line represents the frequency trajectory for a locus of 30 sampled loci from polygenic trait (300 loci).

#### 2.2. Phenotypes all params

##### 2.2.1. Polygenic 70

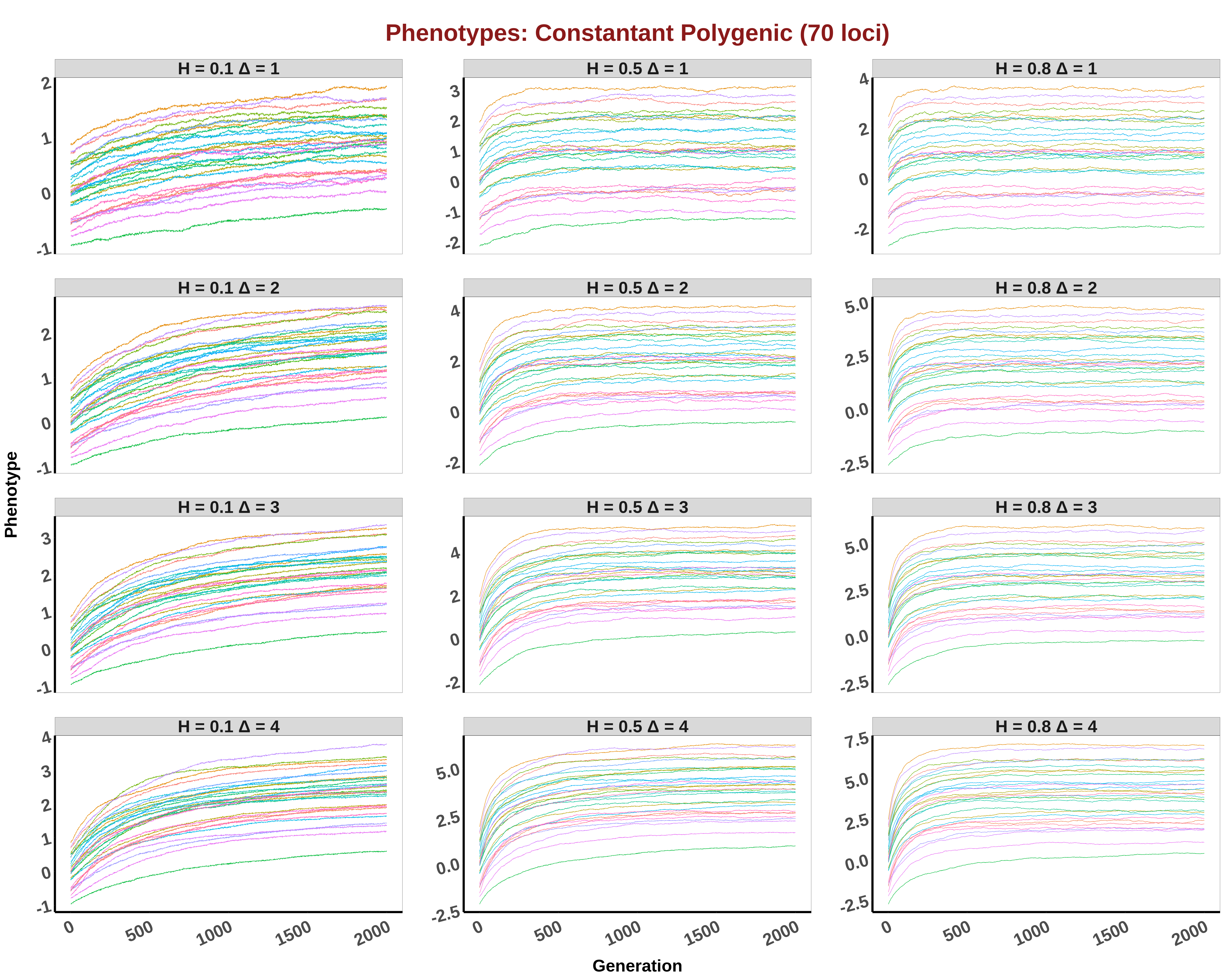

SI 2.2.1. Polygenic (70 loci) mean population phenotypic plot across 30 replicates (each line represents a single replicate) for constant selection, showing parameter interactions.

##### 2.2.2. Polygenic 100

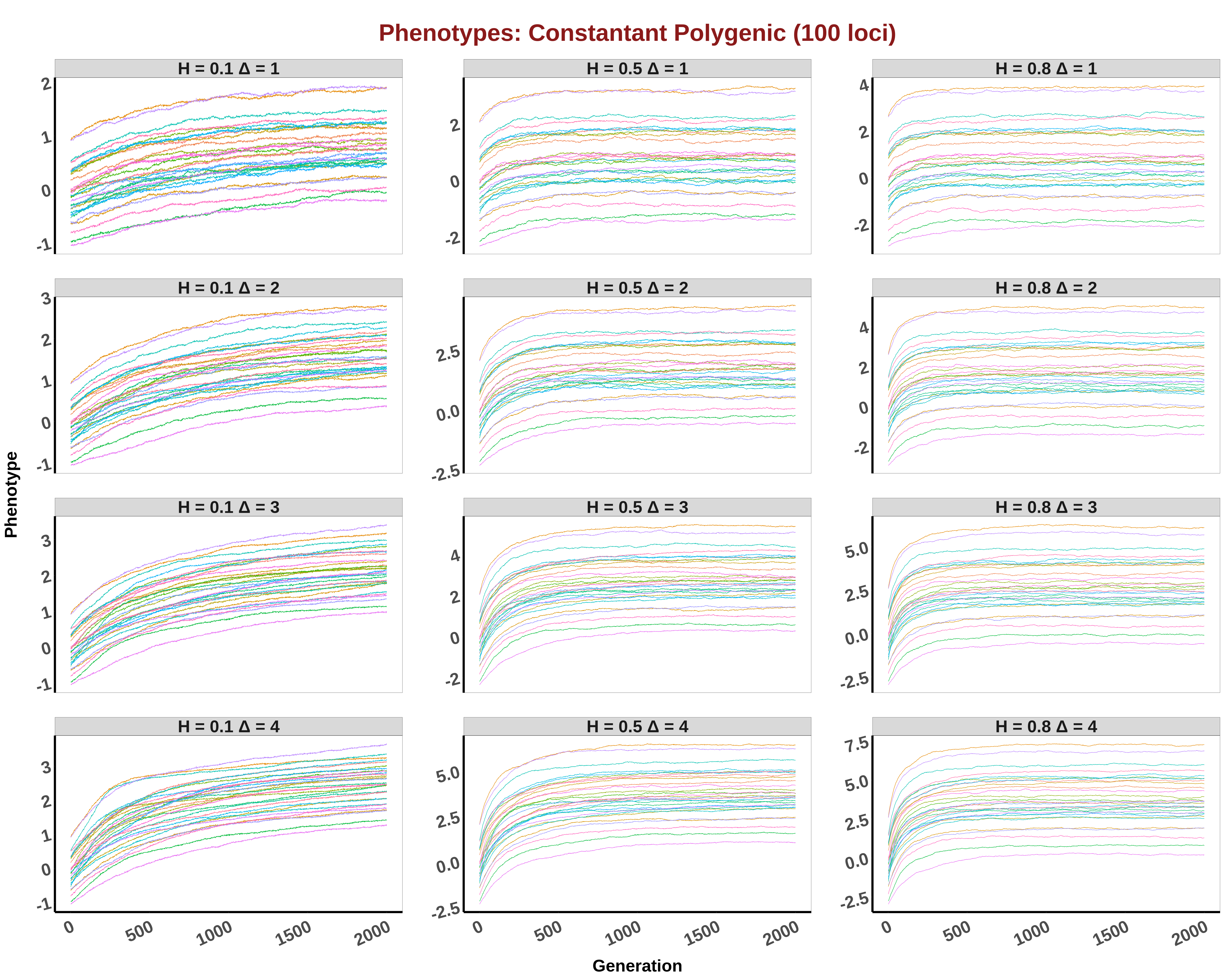

SI 2.2.2 Polygenic (100 loci) mean population phenotypic plot across 30 replicates (each line represents a single replicate) for constant selection, showing parameter interactions.

##### 2.2.3. Polygenic 300

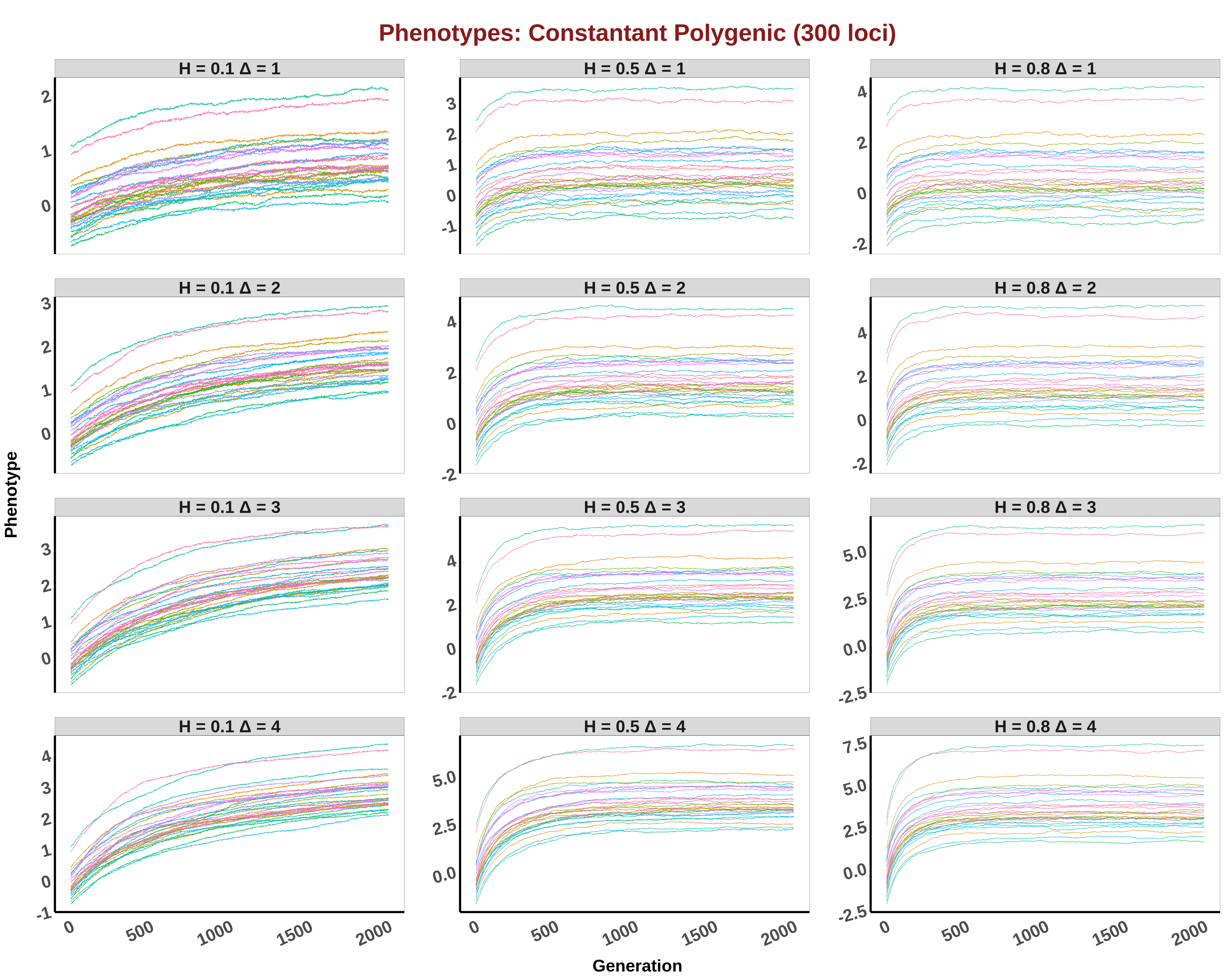

SI 2.2.3. Polygenic (300 loci) mean population phenotypic plot across 30 replicates (each line represents a single replicate) for constant selection, showing parameter interactions.

#### 2.3. Spectral analysis for all parameters

##### 2.3.1. Monogenic

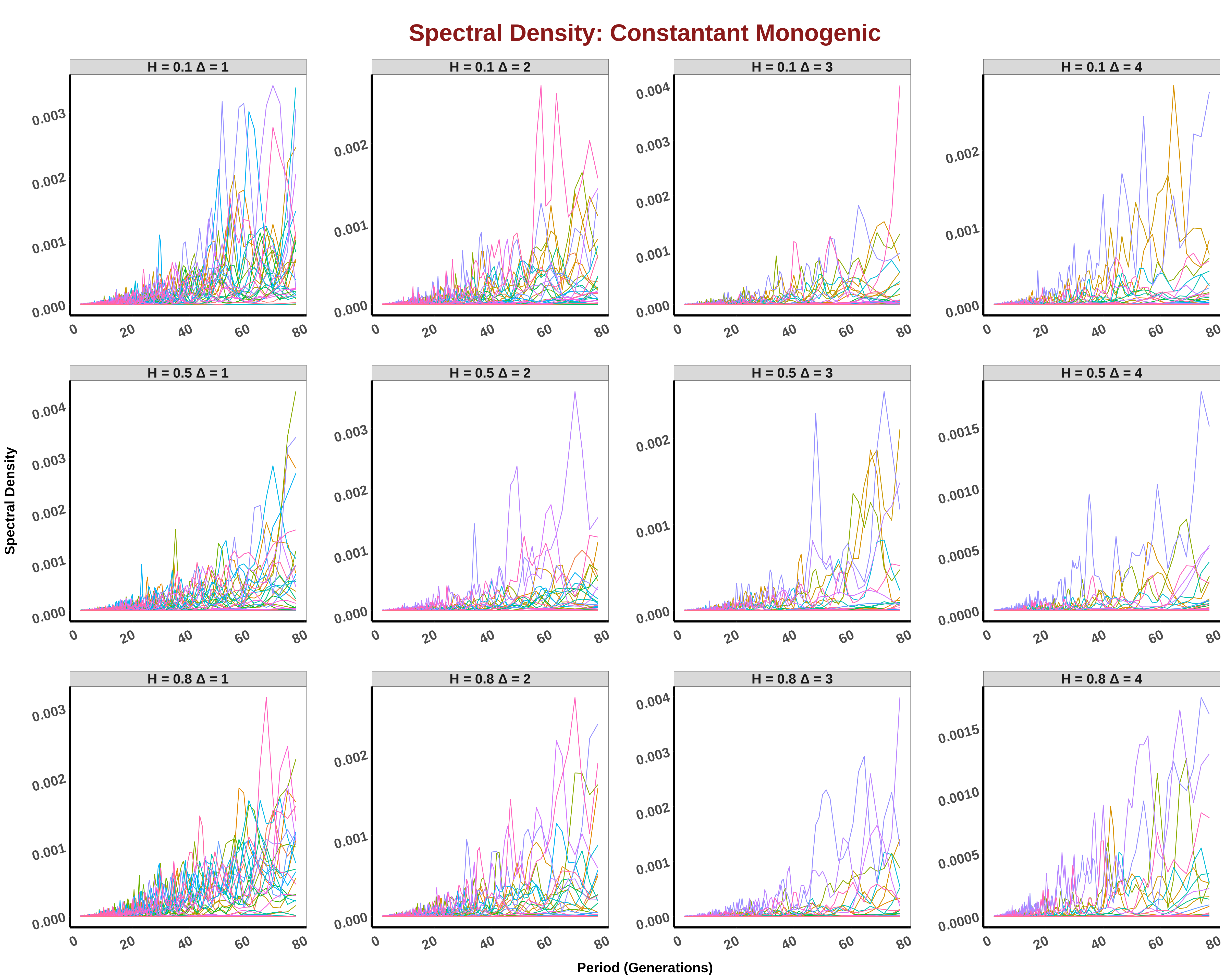

SI 2.3.1. Spectral density across 30 replicates for monogenic trait (single locus) with constant selection model, all parameter combinations. The density plots do not show any peak, and indication that there is no periodicity.

##### 2.3.2. Polygenic 300

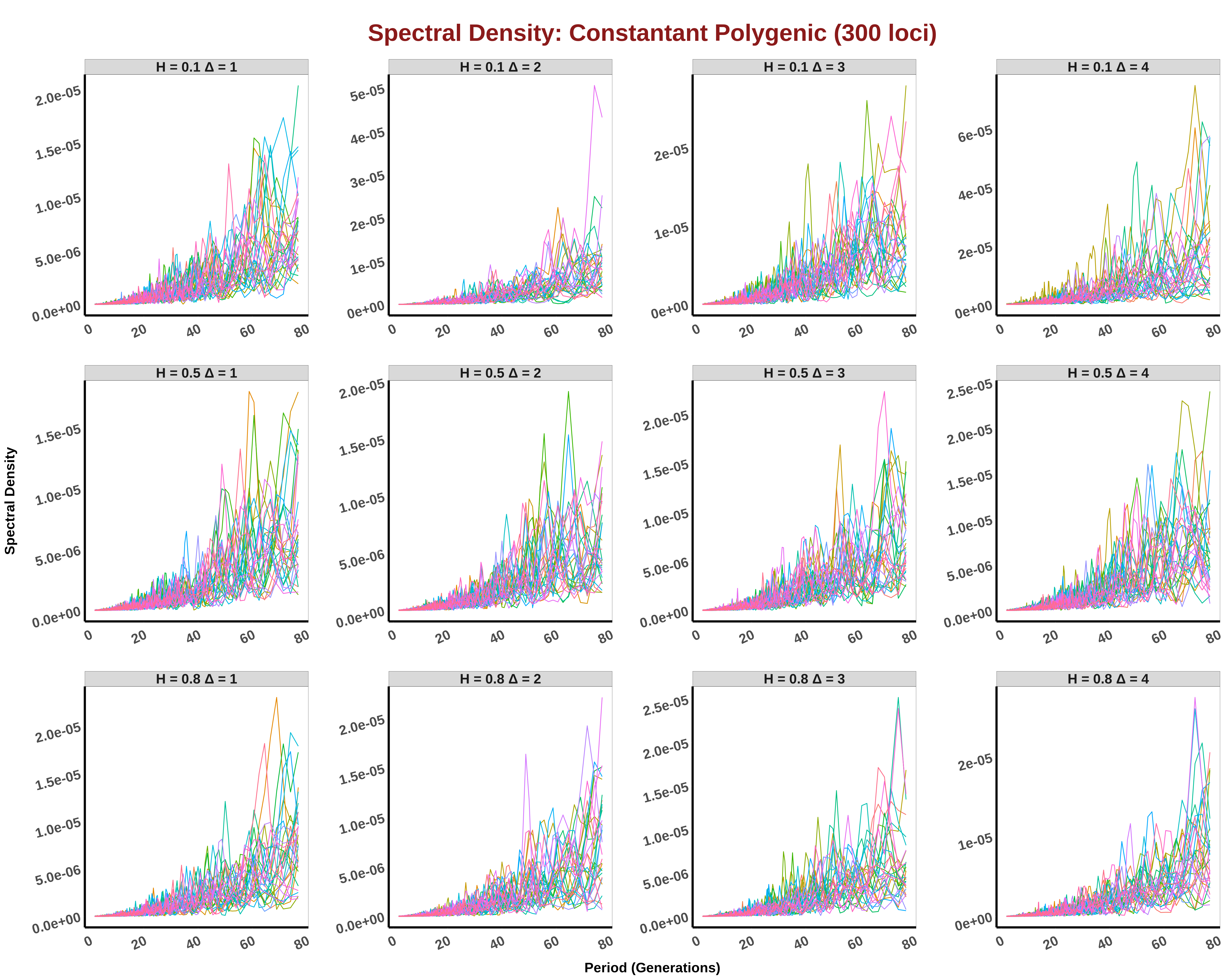

SI 2.3.2. Spectral density across 30 replicates for polygenic trait (300 loci) with constant selection model, all parameter combinations. The density plots do not show any peak, and indication that there is no periodicity.

### 3. Long-term runs with mutation rate

##### 3.1. The cumulative fixation/loss rate of the initial QTL with and without recurrent mutation

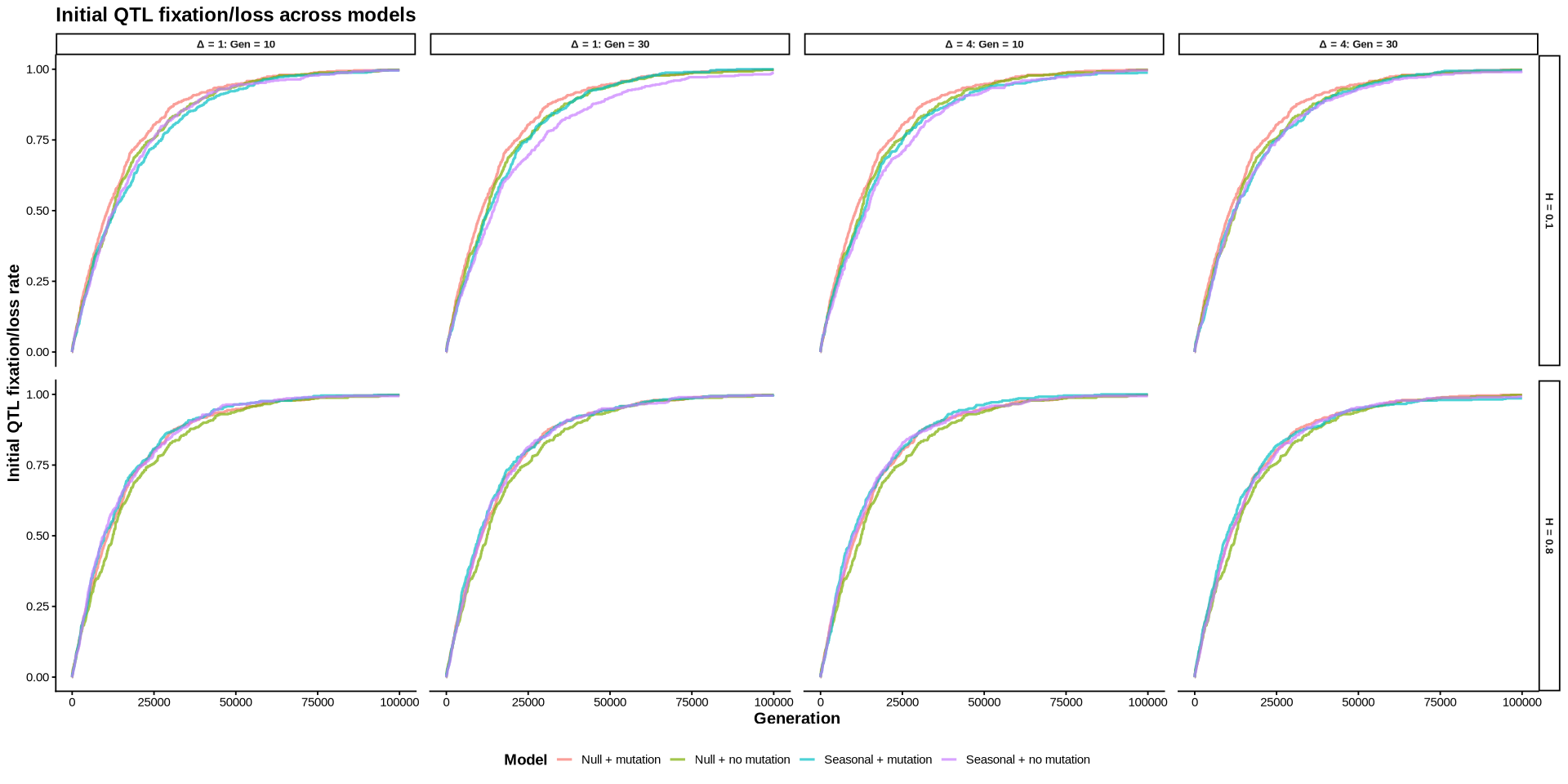

SI 3.1. The cumulative fixation/loss rate of the initial QTL (new mutation excluded) comparison for null and instantaneous fluctuating selection models for a polygenic trait of 100 loci, with and without recurrent mutation, across parameter combinations. The top panel shows the average across 5 replicates, while the bottom panel shows the rate per replicate. In some low-heritability parameter combinations, fluctuating selection seems to exhibit a lower fixation rate than the null model during earlier generations (< 50,000 generations); however, this pattern is not consistent across all parameter combinations, and trajectories converge later in the simulation.

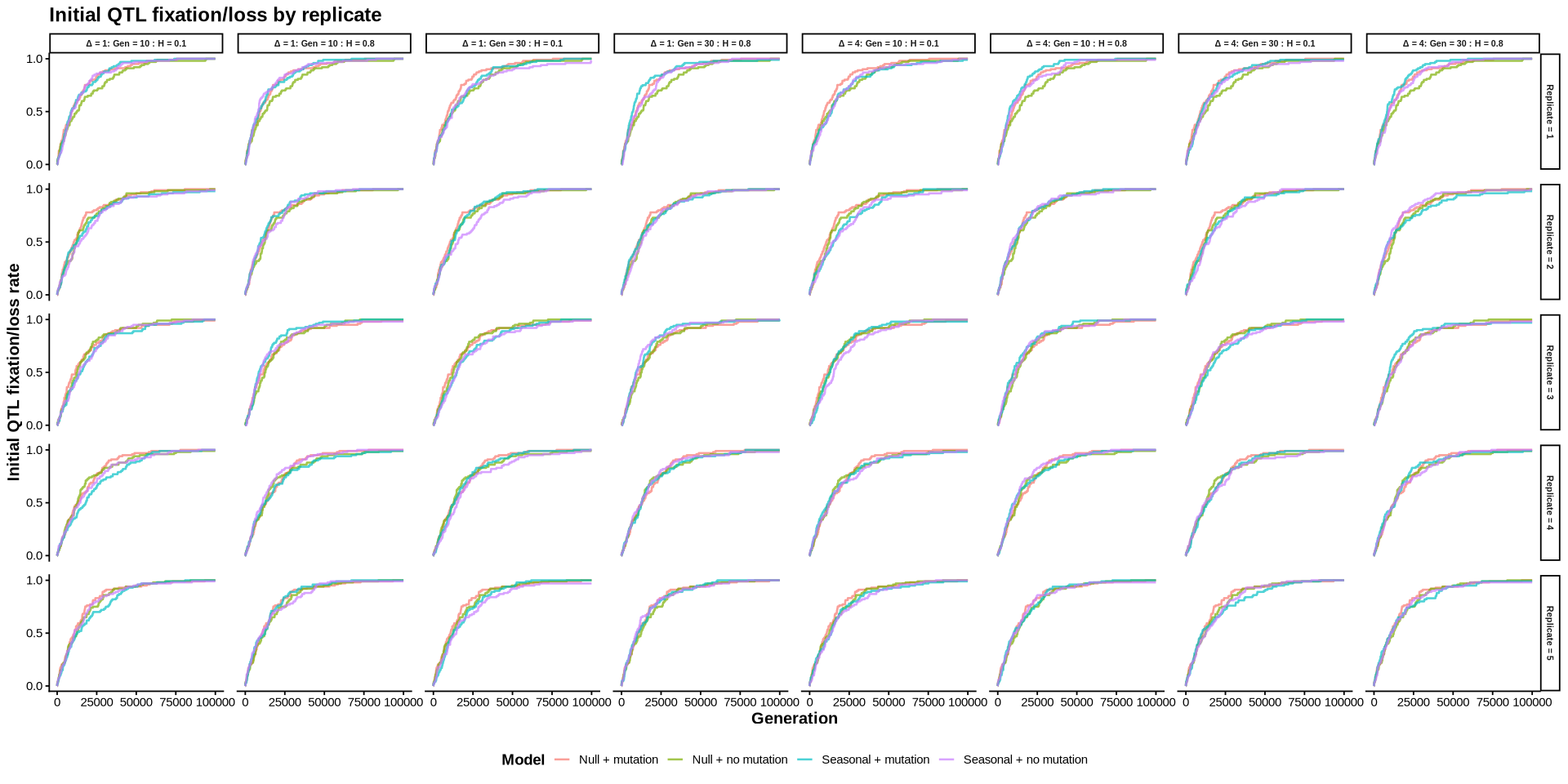

SI 3.1. The cumulative fixation/loss rate of the initial QTL (new mutation excluded) comparison for null and instantaneous fluctuating selection models for a polygenic trait of 100 loci, with and without recurrent mutation, across parameter combinations. The top panel shows the average across 5 replicates, while the bottom panel shows the rate per replicate. In some low-heritability parameter combinations, fluctuating selection seems to exhibit a lower fixation rate than the null model during earlier generations (< 50,000 generations); however, this pattern is not consistent across all parameter combinations, and trajectories converge later in the simulation.

##### 3.2. The average number of segregating sites, comparing null and instantaneous fluctuating selection models

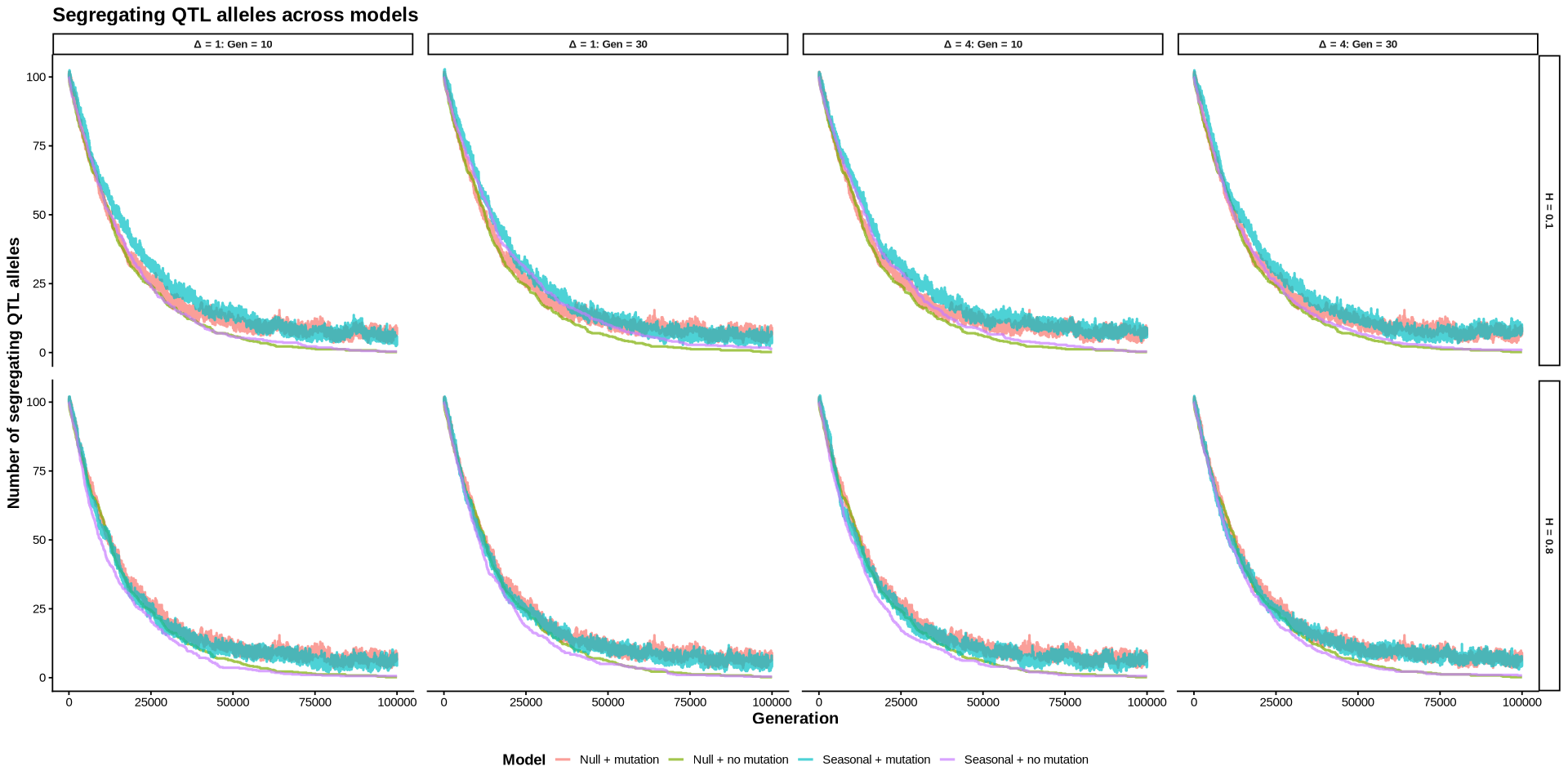

SI 3.2. The average number of segregating sites, comparing null and instantaneous fluctuating selection models for a polygenic trait with 100 loci, with and without recurrent mutation, across parameter combinations. The top panel shows the average number of segregating sites across generations, including new mutations, while the bottom panel shows per replicate changes. Likewise, in some low-heritability parameter combinations, fluctuating selection seems to maintain more segregating variation than the null model during earlier generations (< 50,000 generations), but this pattern is not consistent across all parameter combinations, as trajectories converge later in the simulation.

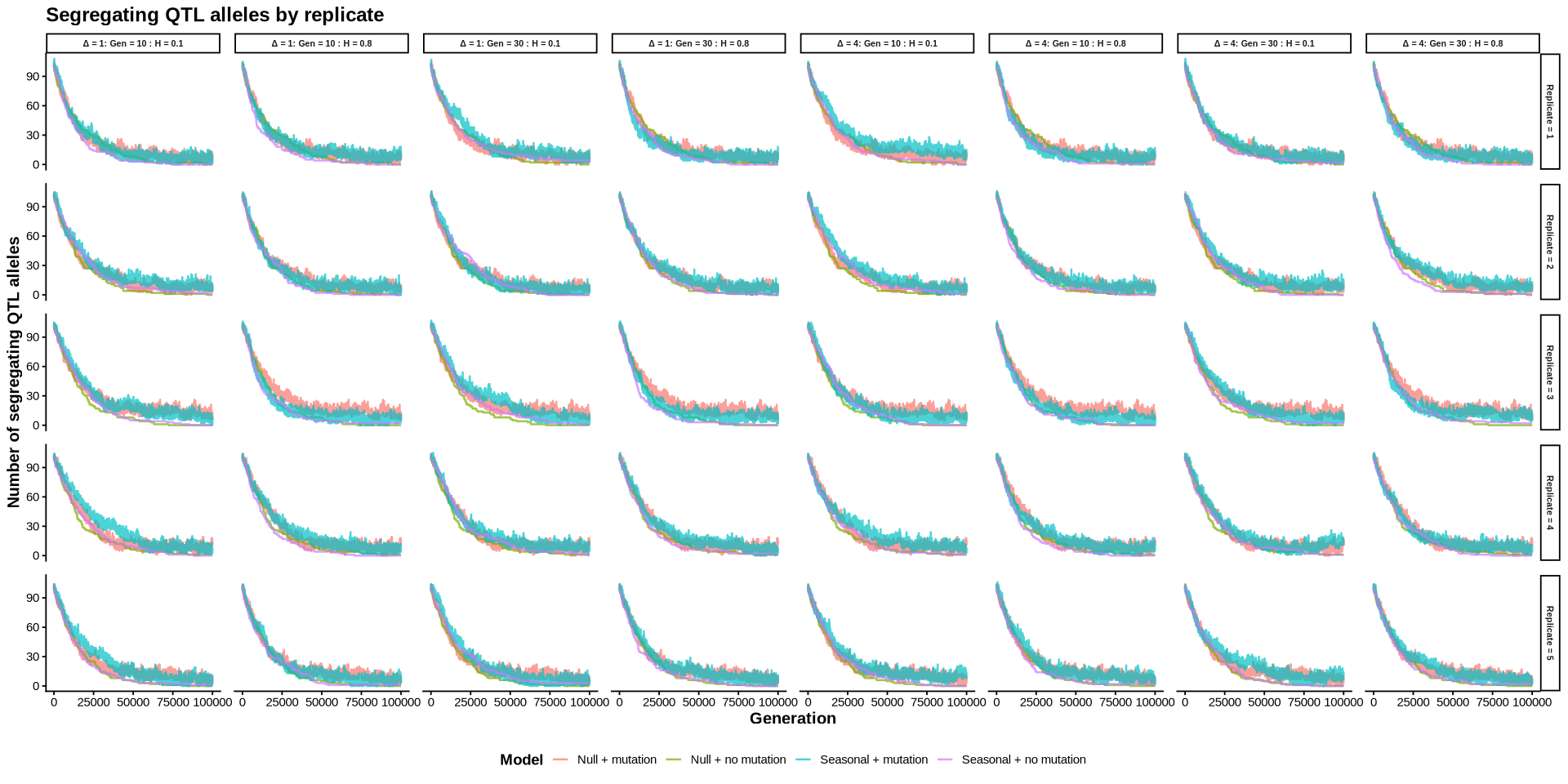

SI 3.2. The average number of segregating sites, comparing null and instantaneous fluctuating selection models for a polygenic trait with 100 loci, with and without recurrent mutation, across parameter combinations. The top panel shows the average number of segregating sites across generations, including new mutations, while the bottom panel shows per replicate changes. Likewise, in some low-heritability parameter combinations, fluctuating selection seems to maintain more segregating variation than the null model during earlier generations (< 50,000 generations), but this pattern is not consistent across all parameter combinations, as trajectories converge later in the simulation.

### 4. Instantaneous model

#### 4.1. Allele frequency all parameters

##### 4.1.1. Monogenic

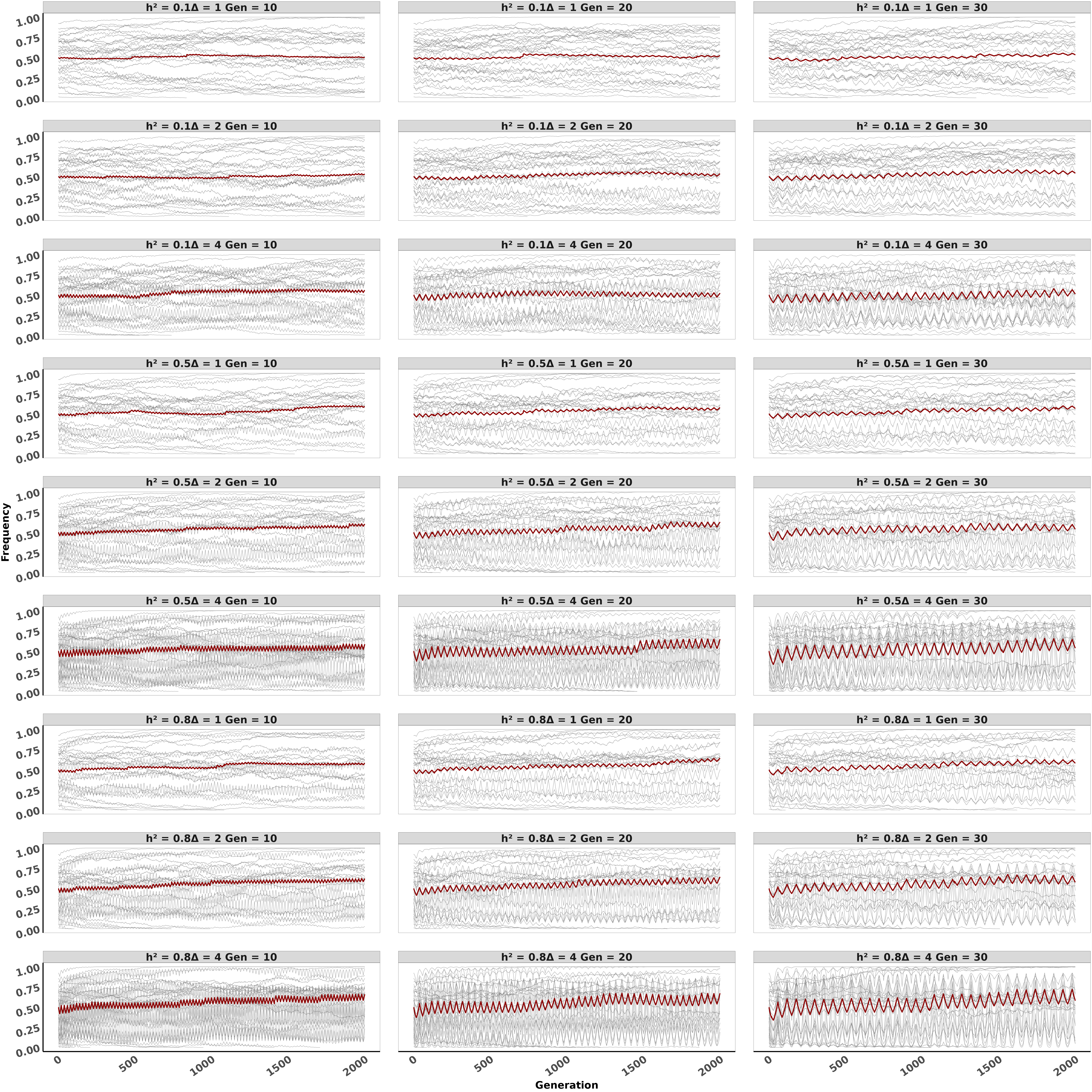

SI 4.1.1. Single locus (30 replicates) allele frequency dynamics for instantaneous selection, showing all parameter interactions. The highlighted line indicates the mean allele frequency for each generation across replicates.

##### 4.1.2. Oligogenic

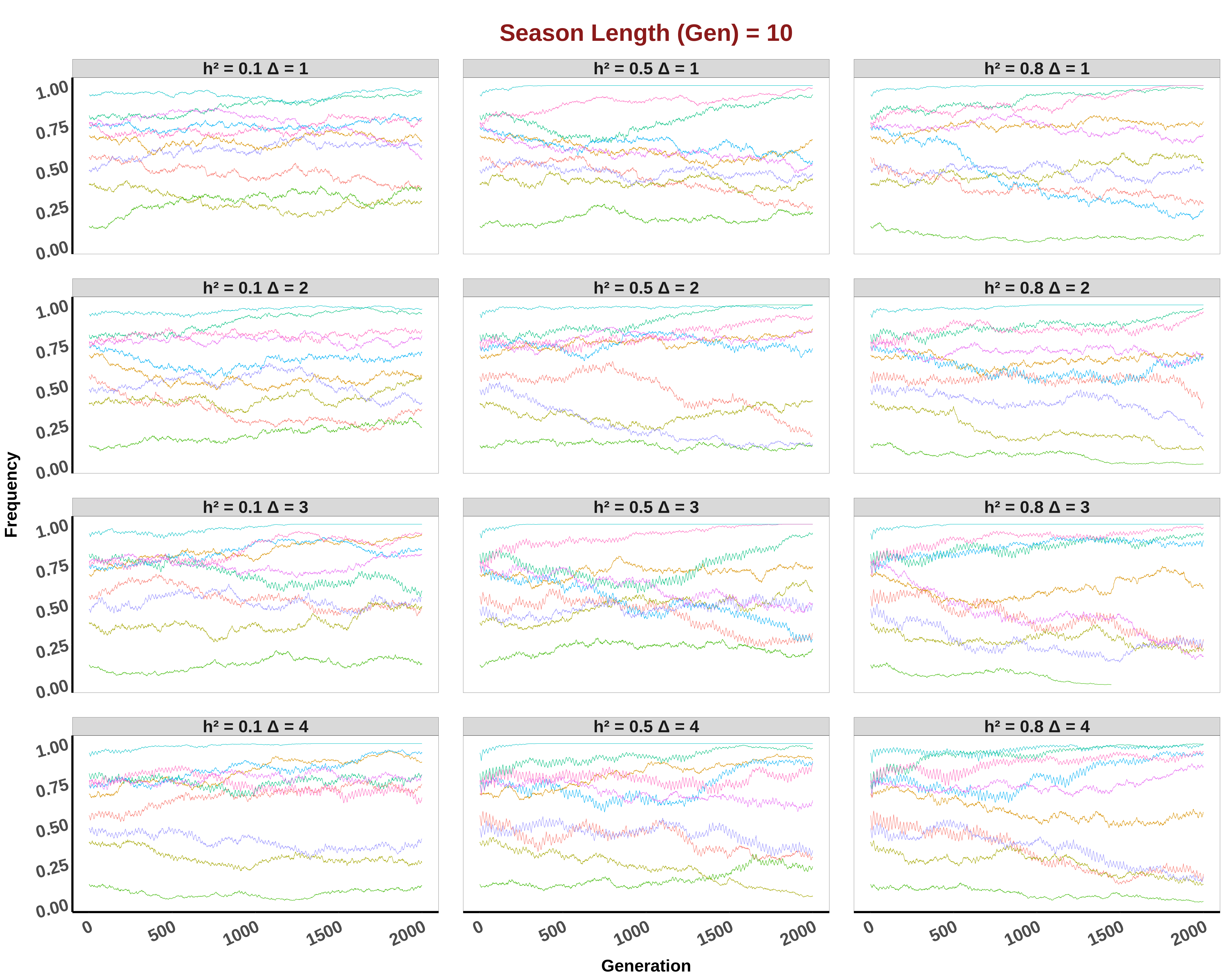

SI 4.1.2.1 Allele frequency dynamics for all parameter combinations with selection length of 10 generations from a single replicate, instantaneous selection model. Each line represents the frequency trajectory for a locus from oligogenic trait (10 loci).

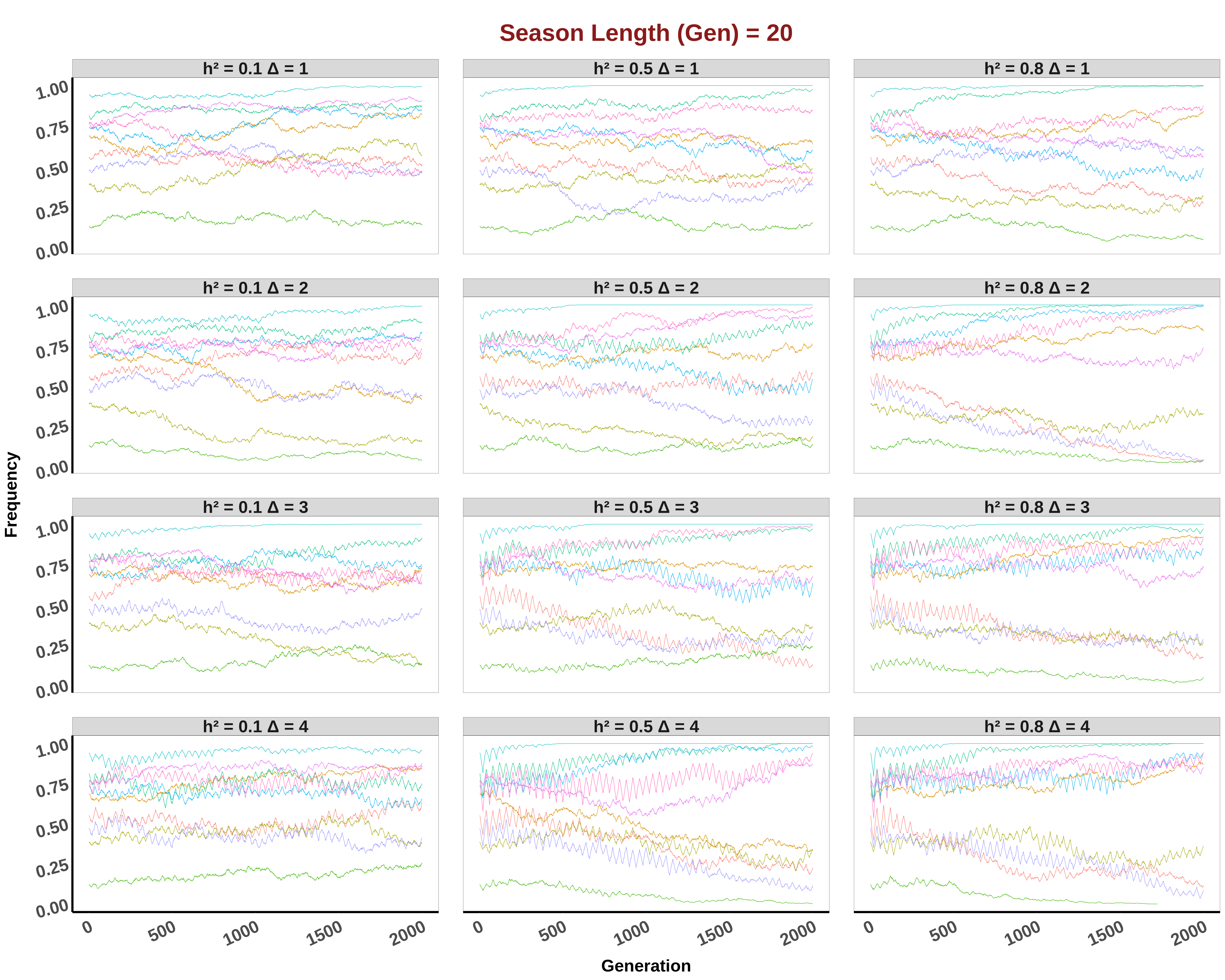

SI 4.1.2.2. Allele frequency dynamics for all parameter combinations with selection length of 20 generations from a single replicate, instantaneous selection model. Each line represents the frequency trajectory for a locus from oligogenic trait (10 loci).

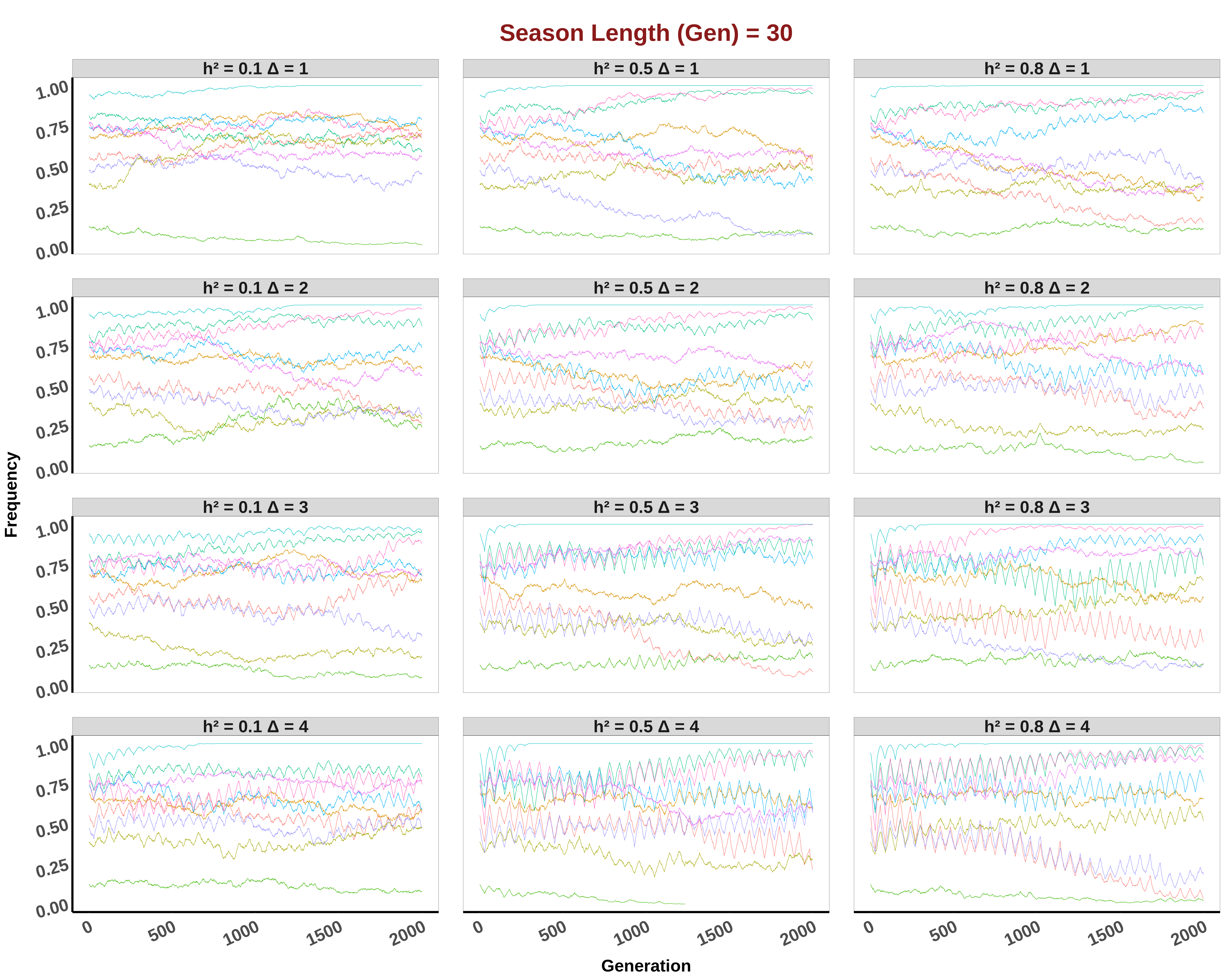

SI 4.1.2.3. Allele frequency dynamics for all parameter combinations with selection length of 30 generations from a single replicate, instantaneous selection model. Each line represents the frequency trajectory for a locus from oligogenic trait (10 loci).

##### 4.1.3. Polygenic 70 loci

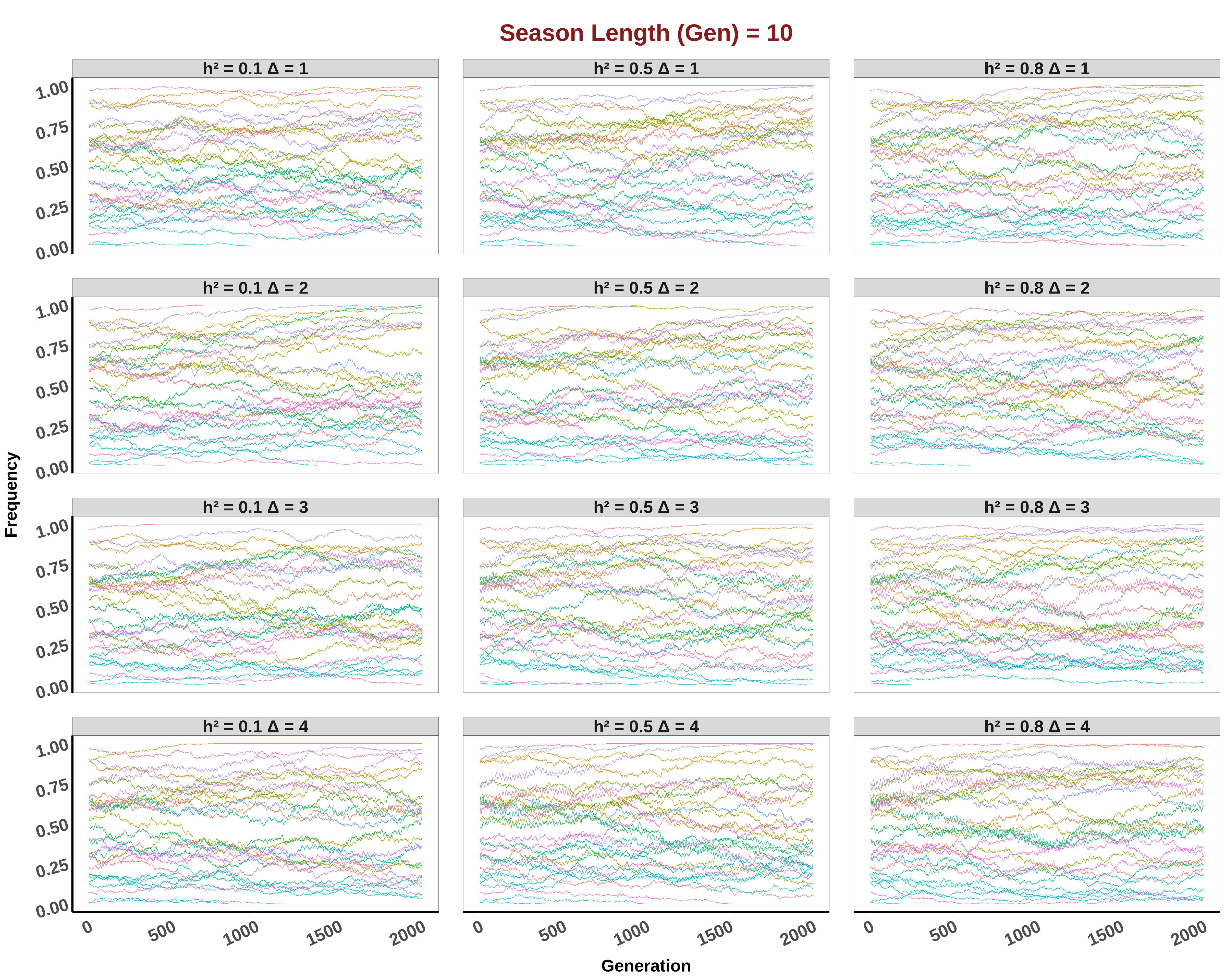

SI 4.1.3.1. Allele frequency dynamics for all parameter combinations with selection length of 10 generations from a single replicate, instantaneous selection model. Each line represents the frequency trajectory for a locus from polygenic trait (70 loci).

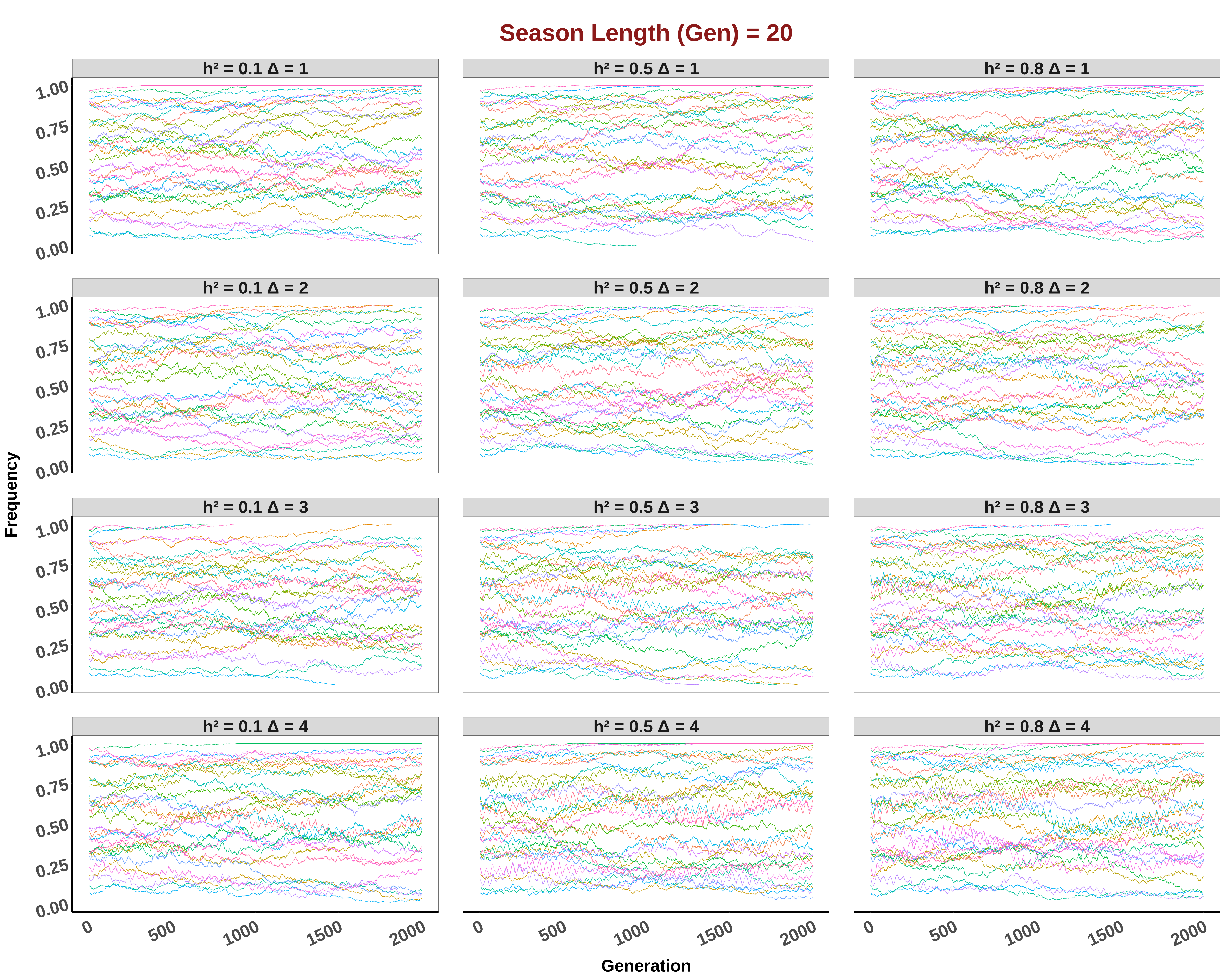

SI 4.1.3.2. Allele frequency dynamics for all parameter combinations with selection length of 20 generations from a single replicate, instantaneous selection model. Each line represents the frequency trajectory for a locus from polygenic trait (70 loci).

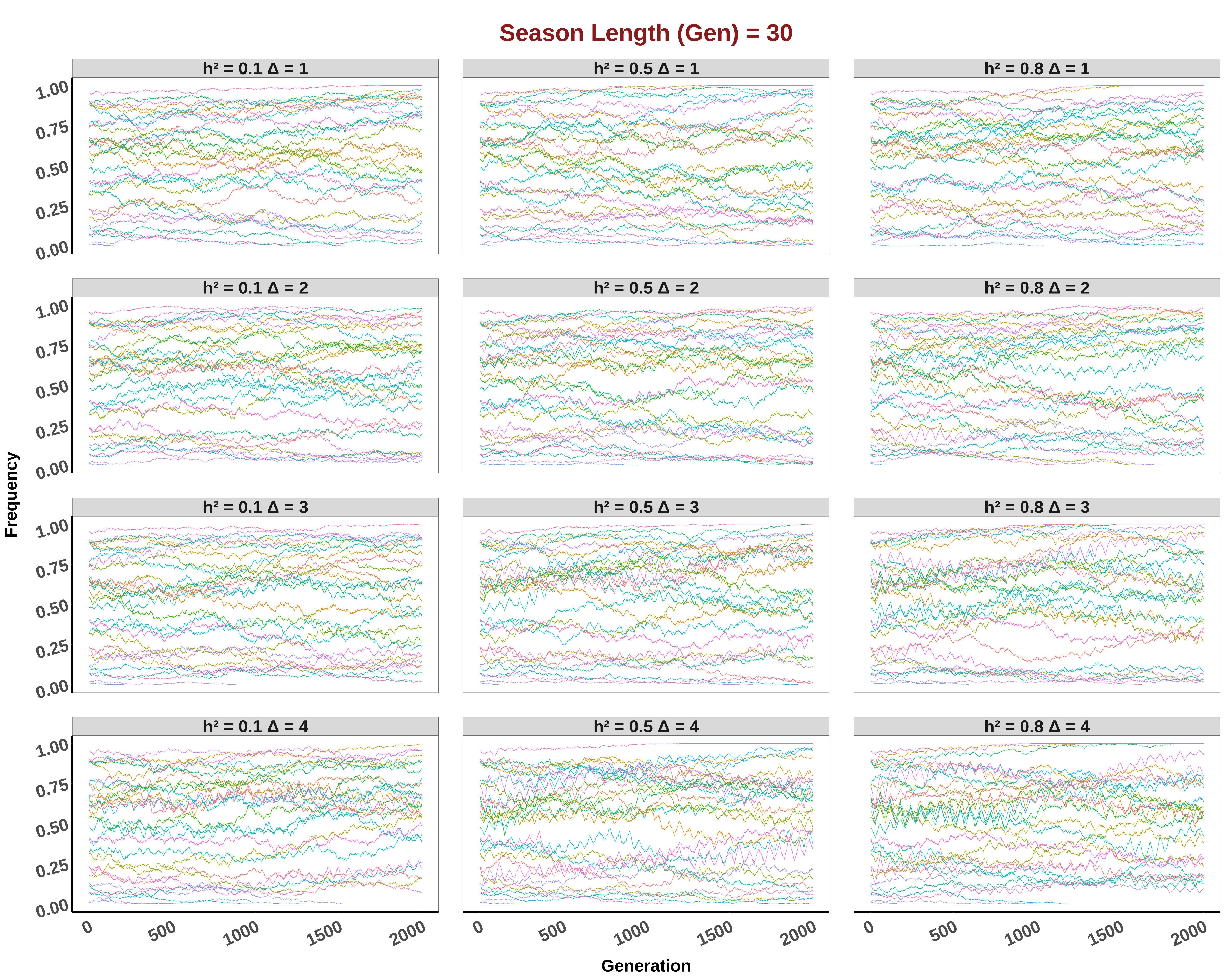

SI 4.1.3.3. Allele frequency dynamics for all parameter combinations with selection length of 30 generations from a single replicate, instantaneous selection model. Each line represents the frequency trajectory for a locus from polygenic trait (70 loci).

##### 4.1.4. Polygenic 300 loci

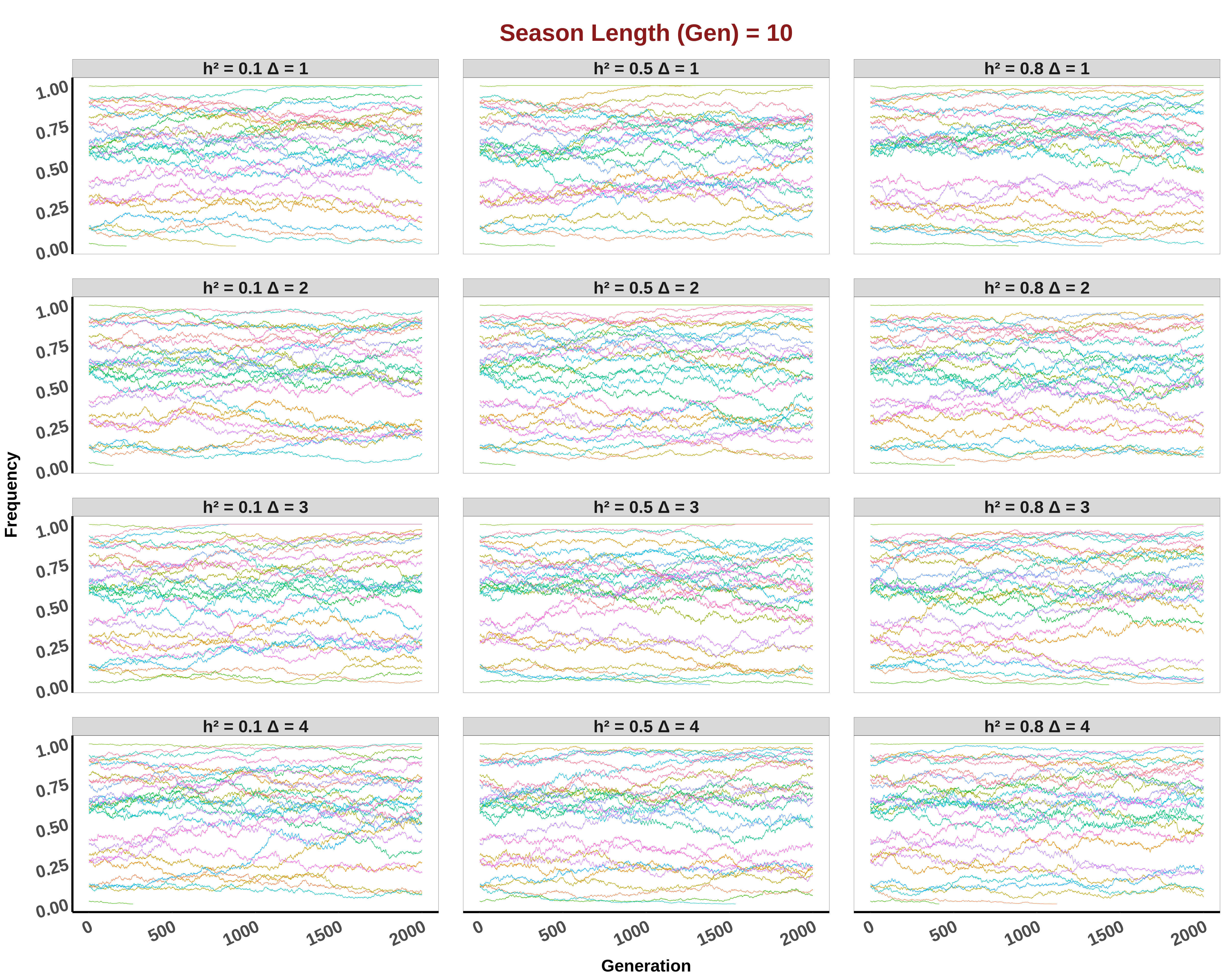

SI 4.1.4.1. Allele frequency dynamics for all parameter combinations with selection length of 10 generations from a single replicate, instantaneous selection model. Each line represents the frequency trajectory for a locus from polygenic trait (300 loci).

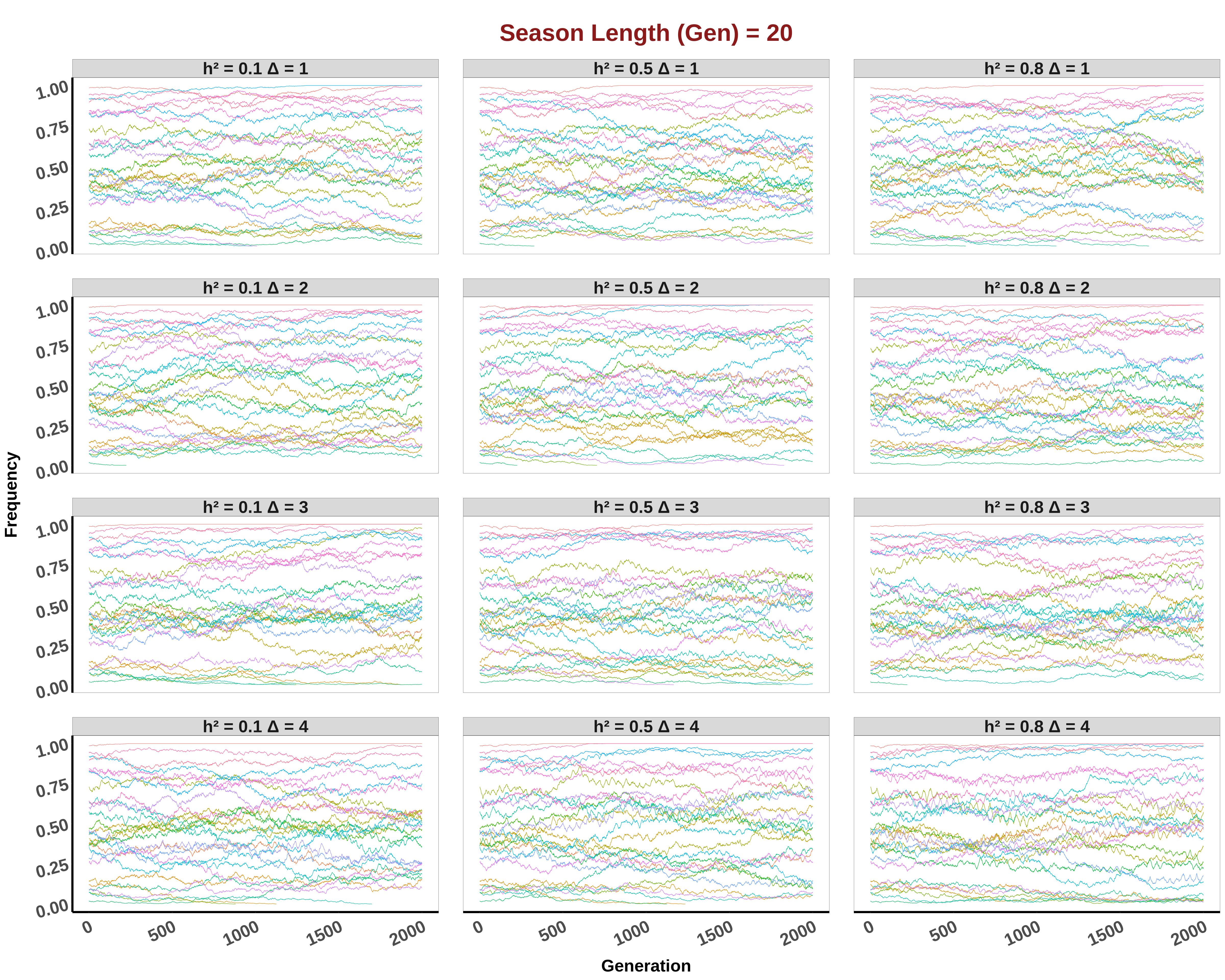

SI 4.1.4.2. Allele frequency dynamics for all parameter combinations with selection length of 20 generations from a single replicate, instantaneous selection model. Each line represents the frequency trajectory for a locus from polygenic trait (300 loci).

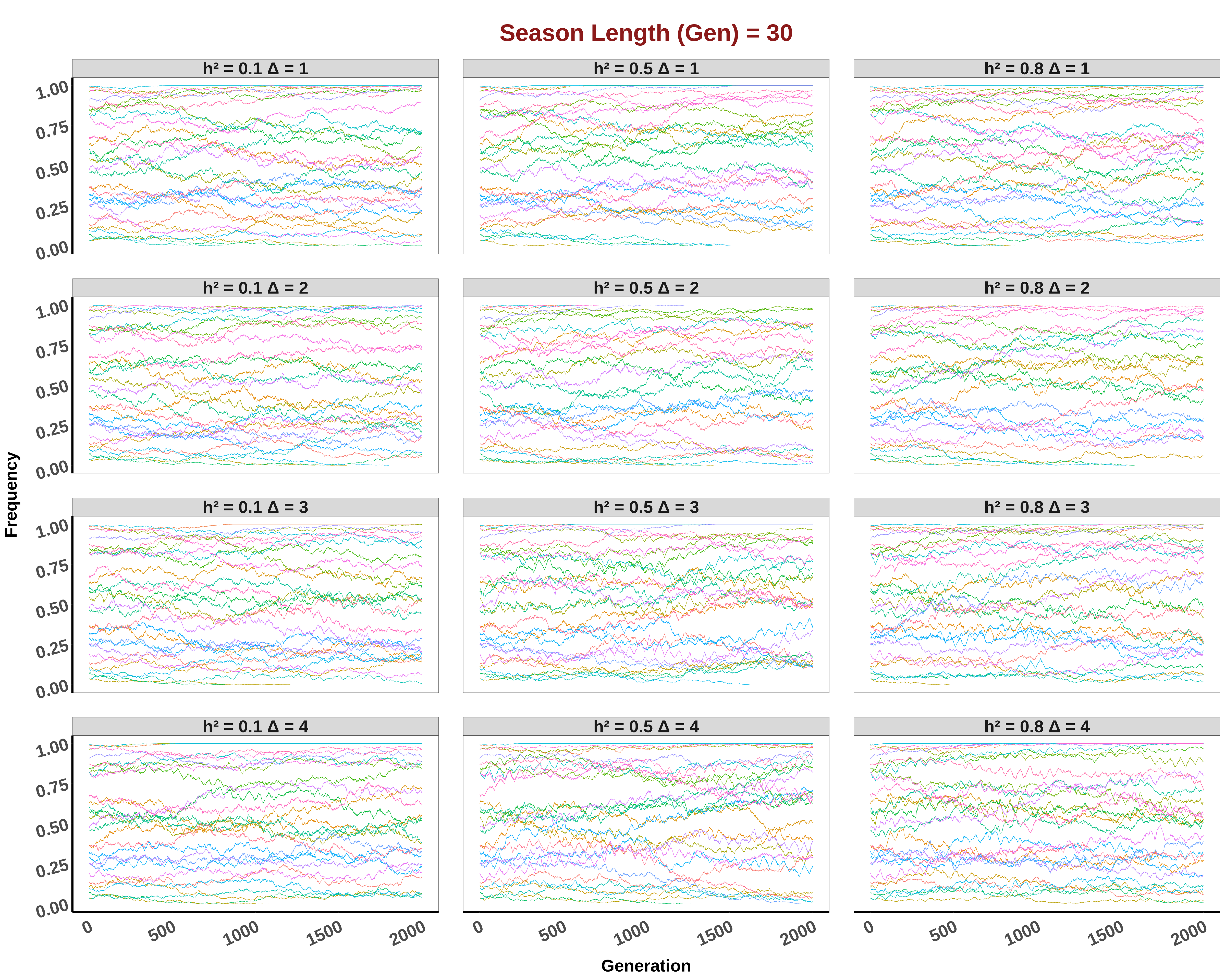

SI 4.1.4.3. Allele frequency dynamics for all parameter combinations with selection length of 30 generations from a single replicate, instantaneous selection model. Each line represents the frequency trajectory for a locus from polygenic trait (300 loci).

#### 4.2. Spectral analysis for all parameters

###### 4.2.1. Monogenic

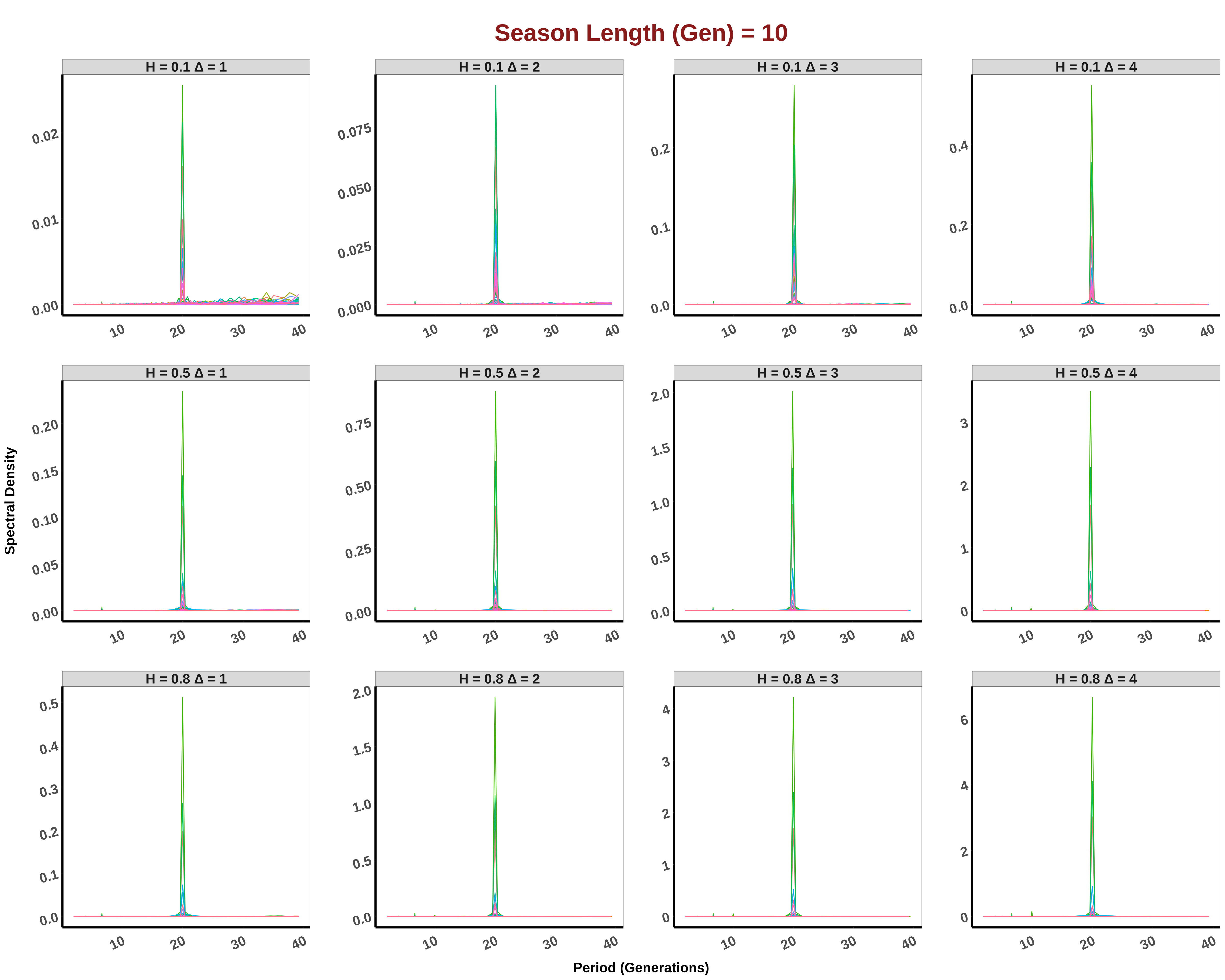

SI 4.2.1.1. Spectral density across 30 replicates for monogenic trait (single locus) with instantaneous selection model, all parameter combinations with selection length of 10 generations. Each peak represents a full cycle of selection period.

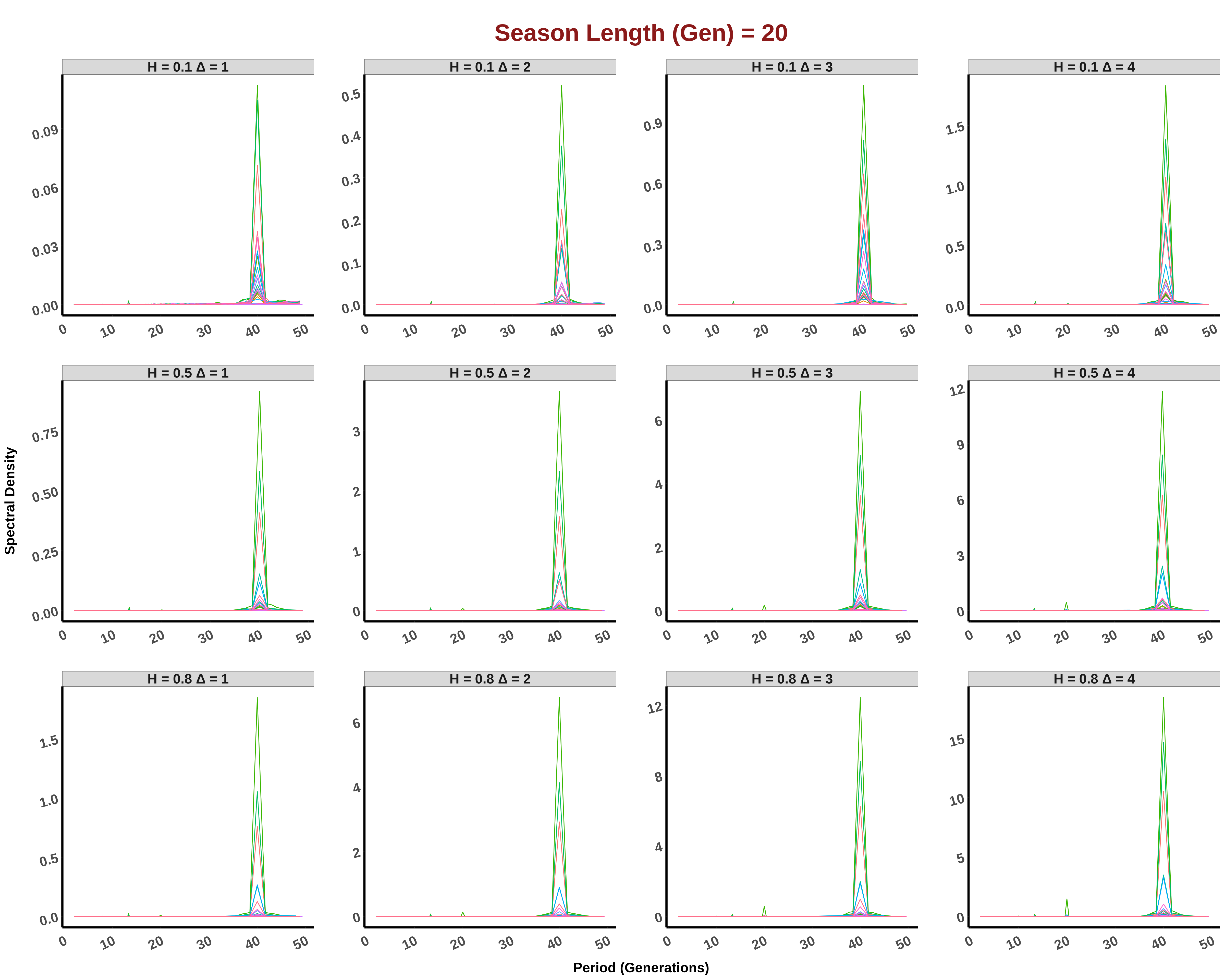

SI 4.2.1.2. Spectral density across 30 replicates for monogenic trait (single locus) with instantaneous selection model, all parameter combinations with selection length of 20 generations. Each peak represents a full cycle of selection period.

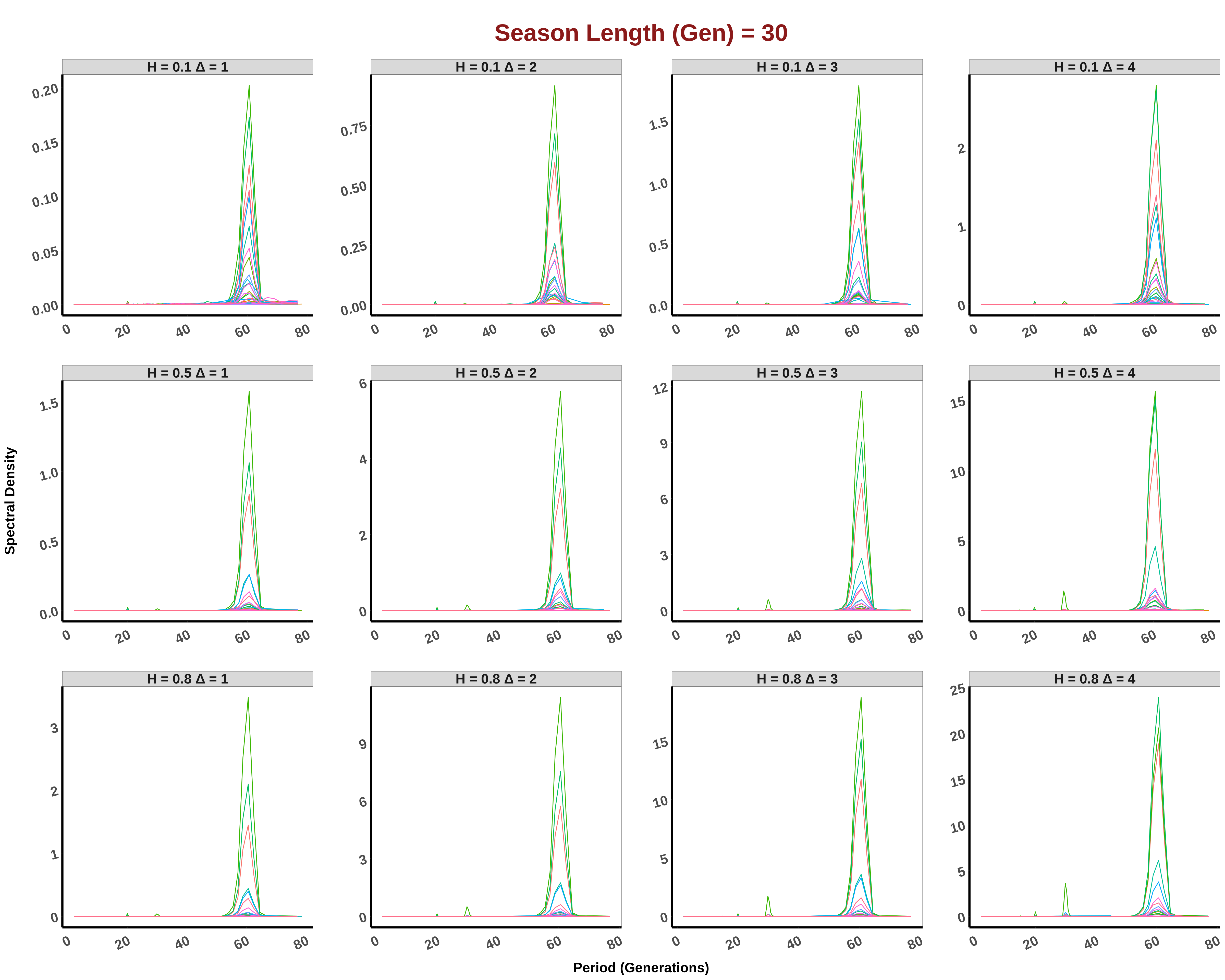

SI 4.2.1.3. Spectral density across 30 replicates for monogenic trait (single locus) with instantaneous selection model, all parameter combinations with selection length of 30 generations. Each peak represents a full cycle of selection period.

##### 4.2.2. Oligogenic

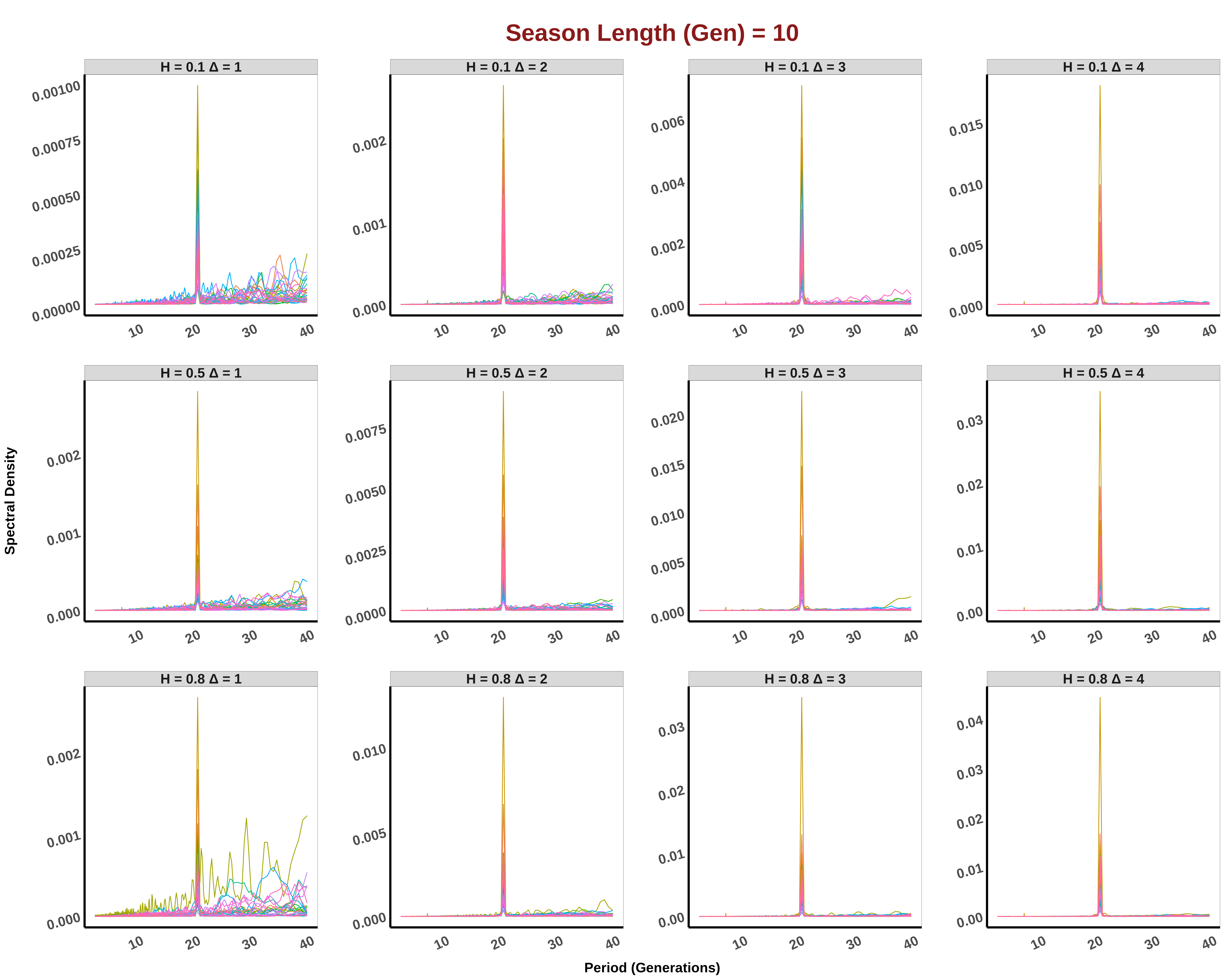

SI 4.2.2.1. Spectral density across 30 replicates for oligogenic trait (10 loci) with instantaneous selection model, all parameter combinations with selection length of 10 generations. Each peak represents a full cycle of selection period.

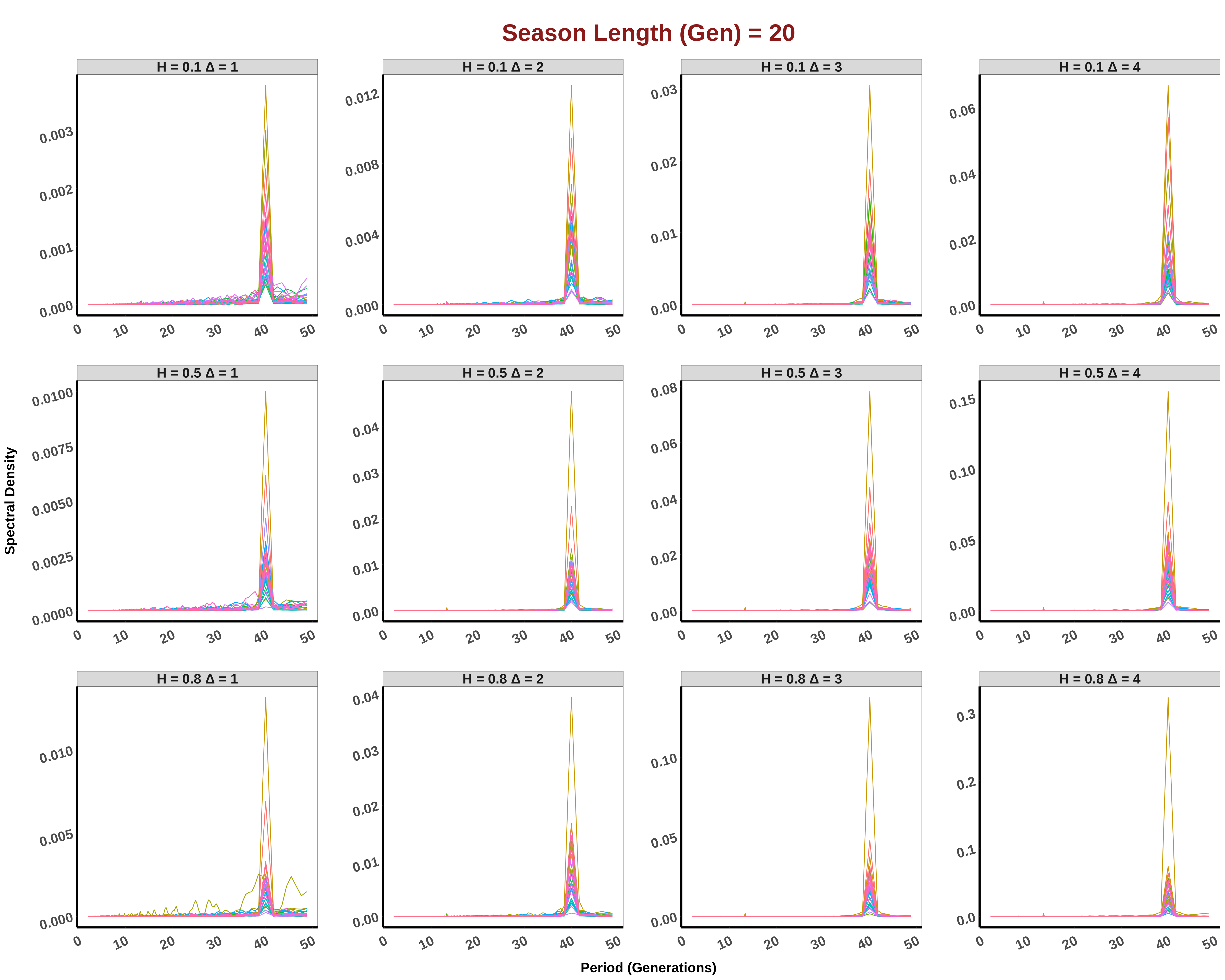

SI 4.2.2.2. Spectral density across 30 replicates for oligogenic trait (10 loci) with instantaneous selection model, all parameter combinations with selection length of 20 generations. Each peak represents a full cycle of selection period.

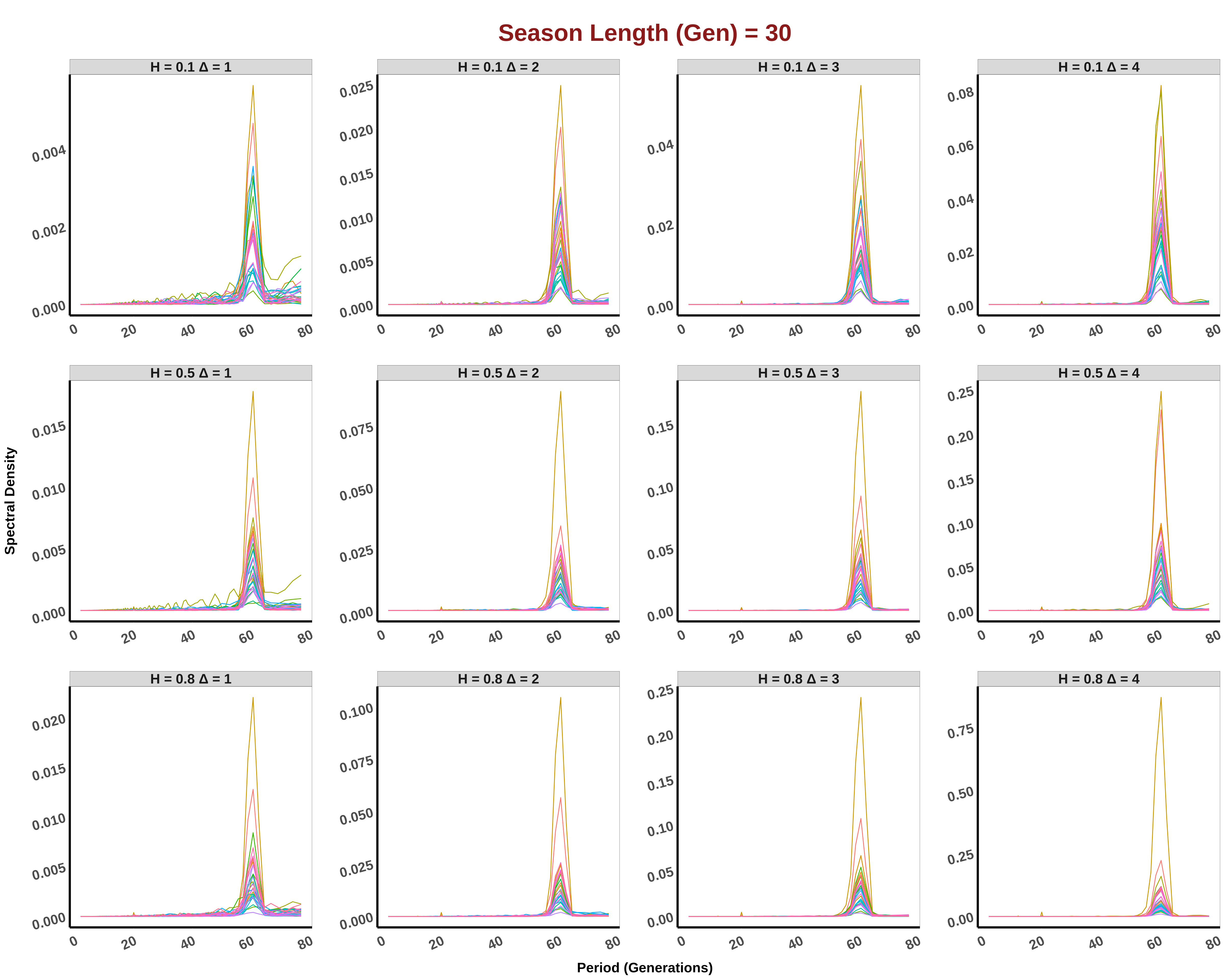

SI 4.2.2.3. Spectral density across 30 replicates for oligogenic trait (10 loci) with instantaneous selection model, all parameter combinations with selection length of 30 generations. Each peak represents a full cycle of selection period.

##### 4.2.3. Polygenic 70

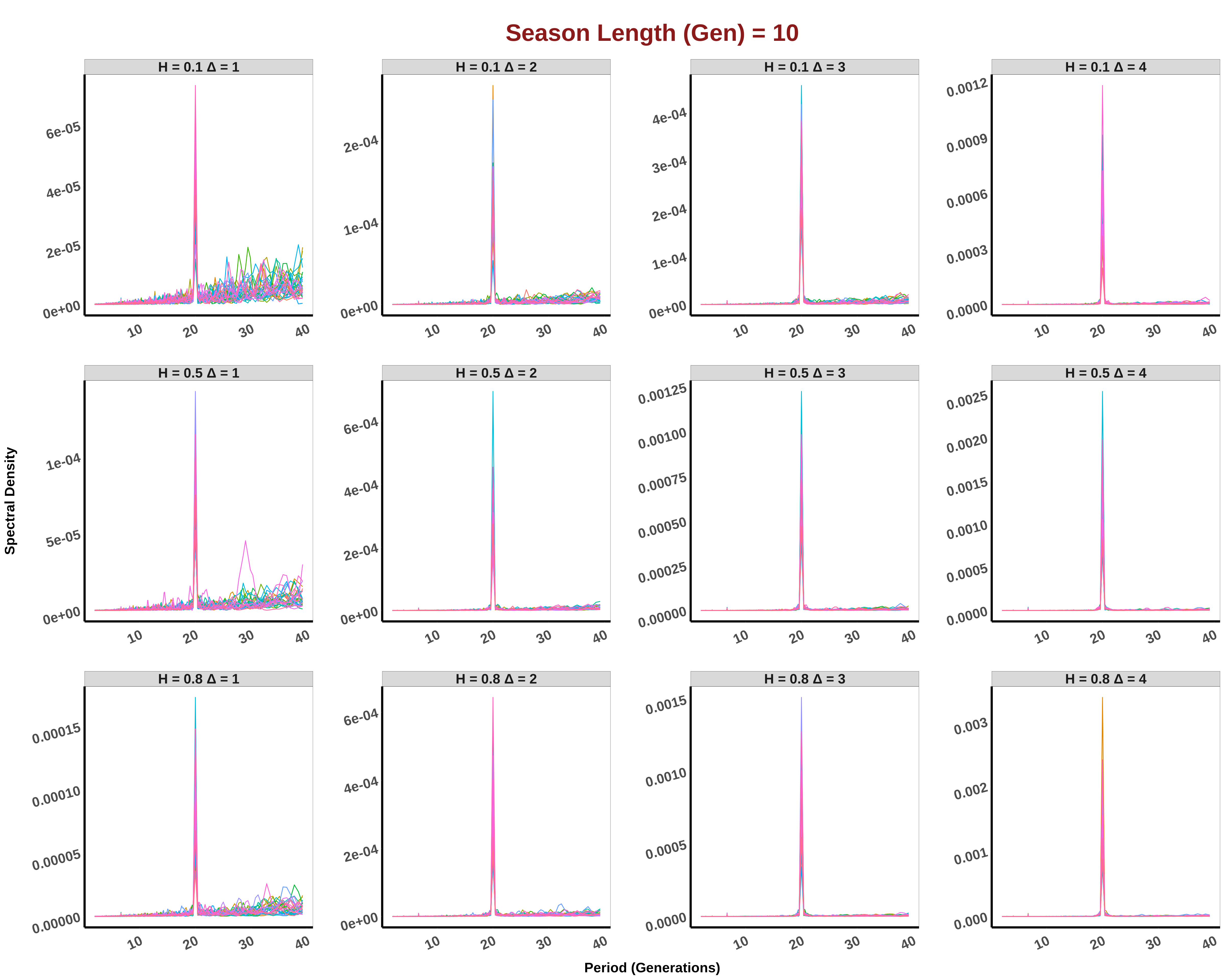

SI 4.2.3.1. Spectral density across 30 replicates for polygenic trait (70 loci) with instantaneous selection model, all parameter combinations with selection length of 10 generations. Each peak represents a full cycle of selection period.

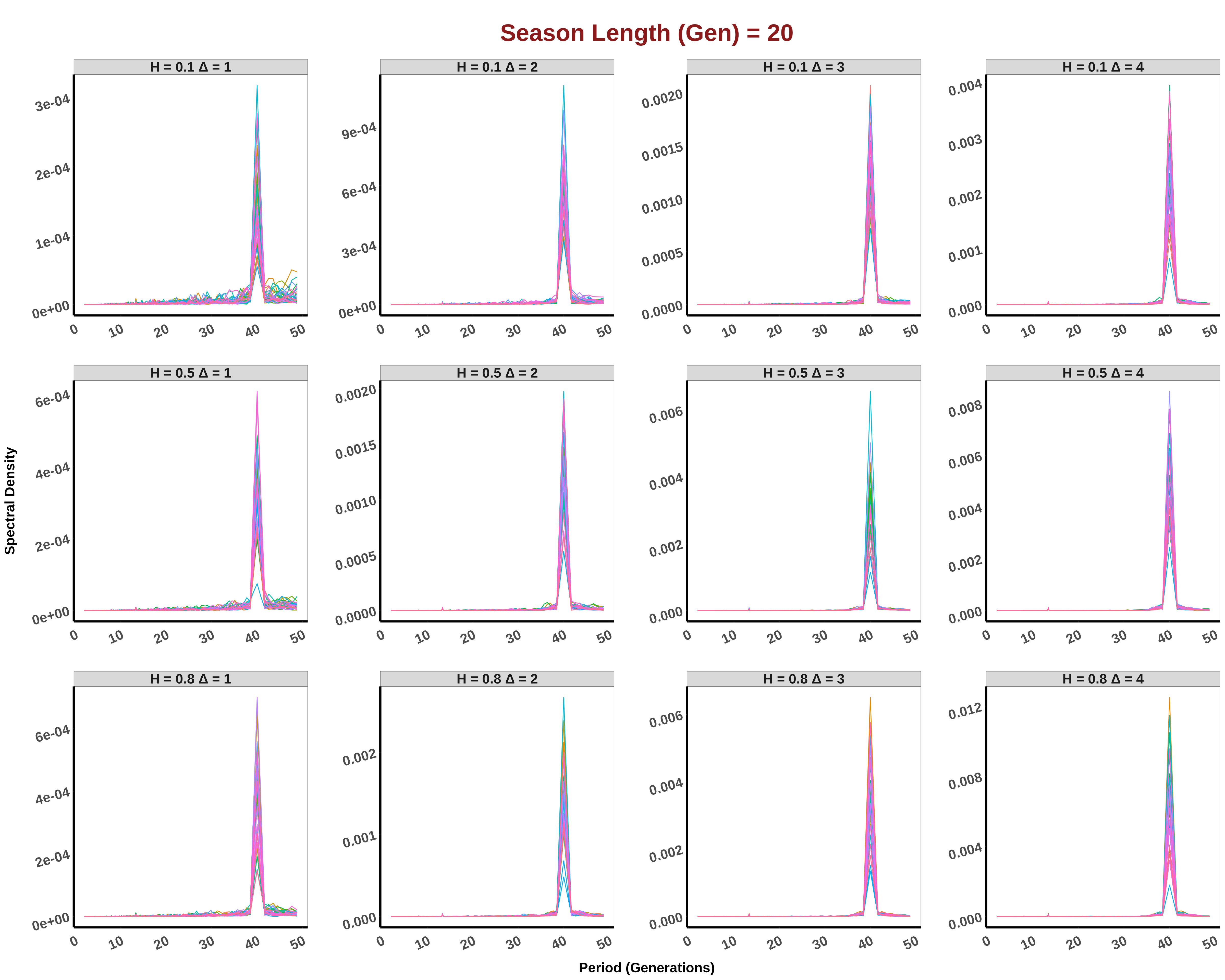

SI 4.2.3.2. Spectral density across 30 replicates for polygenic trait (70 loci) with instantaneous selection model, all parameter combinations with selection length of 20 generations. Each peak represents a full cycle of selection period.

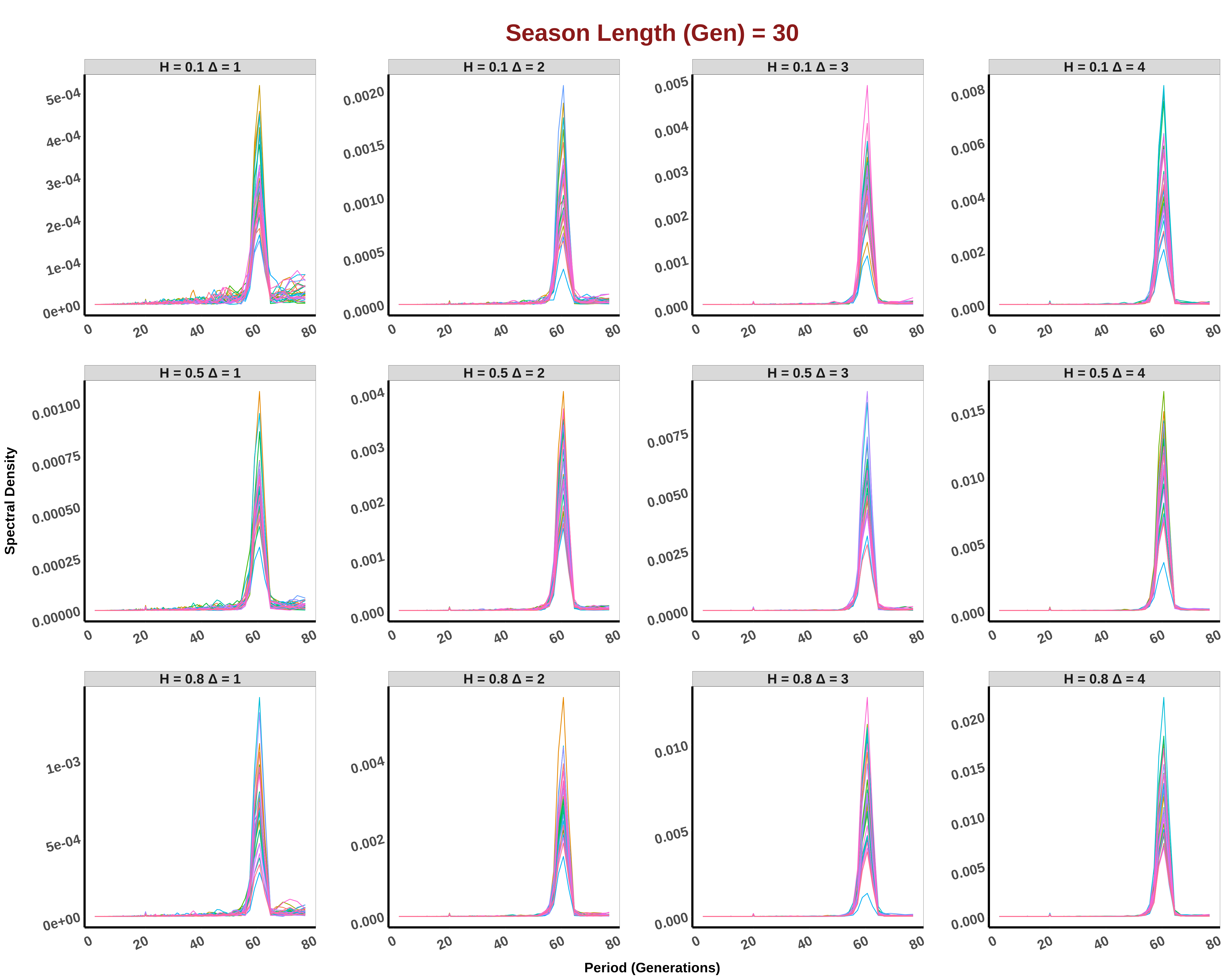

SI 4.2.3.3. Spectral density across 30 replicates for polygenic trait (70 loci) with instantaneous selection model, all parameter combinations with selection length of 30 generations. Each peak represents a full cycle of selection period.

##### 4.2.4. Polygenic 300

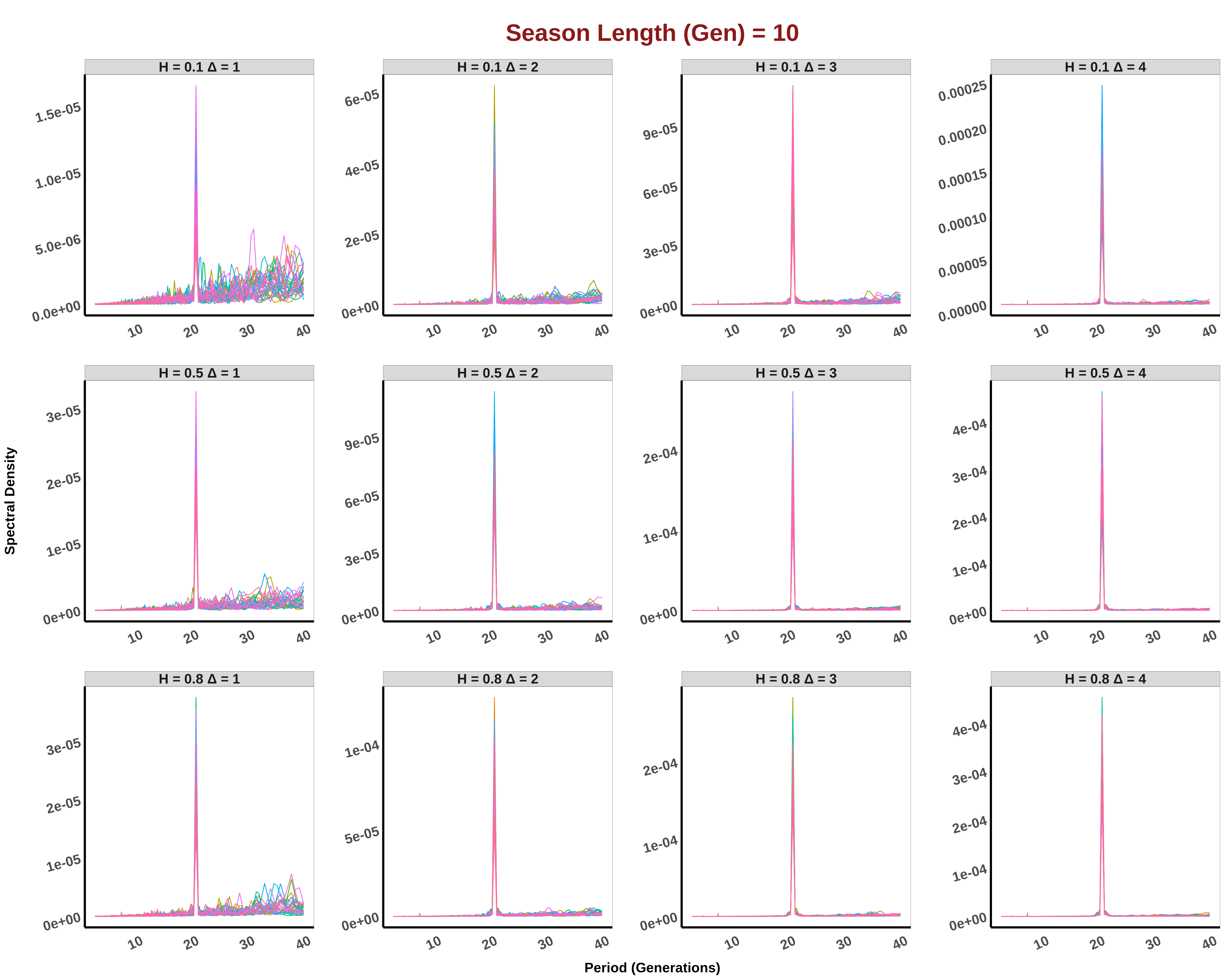

SI 4.2.4.1. Spectral density across 30 replicates for polygenic trait (300 loci) with instantaneous selection model, all parameter combinations with selection length of 10 generations. Each peak represents a full cycle of selection period.

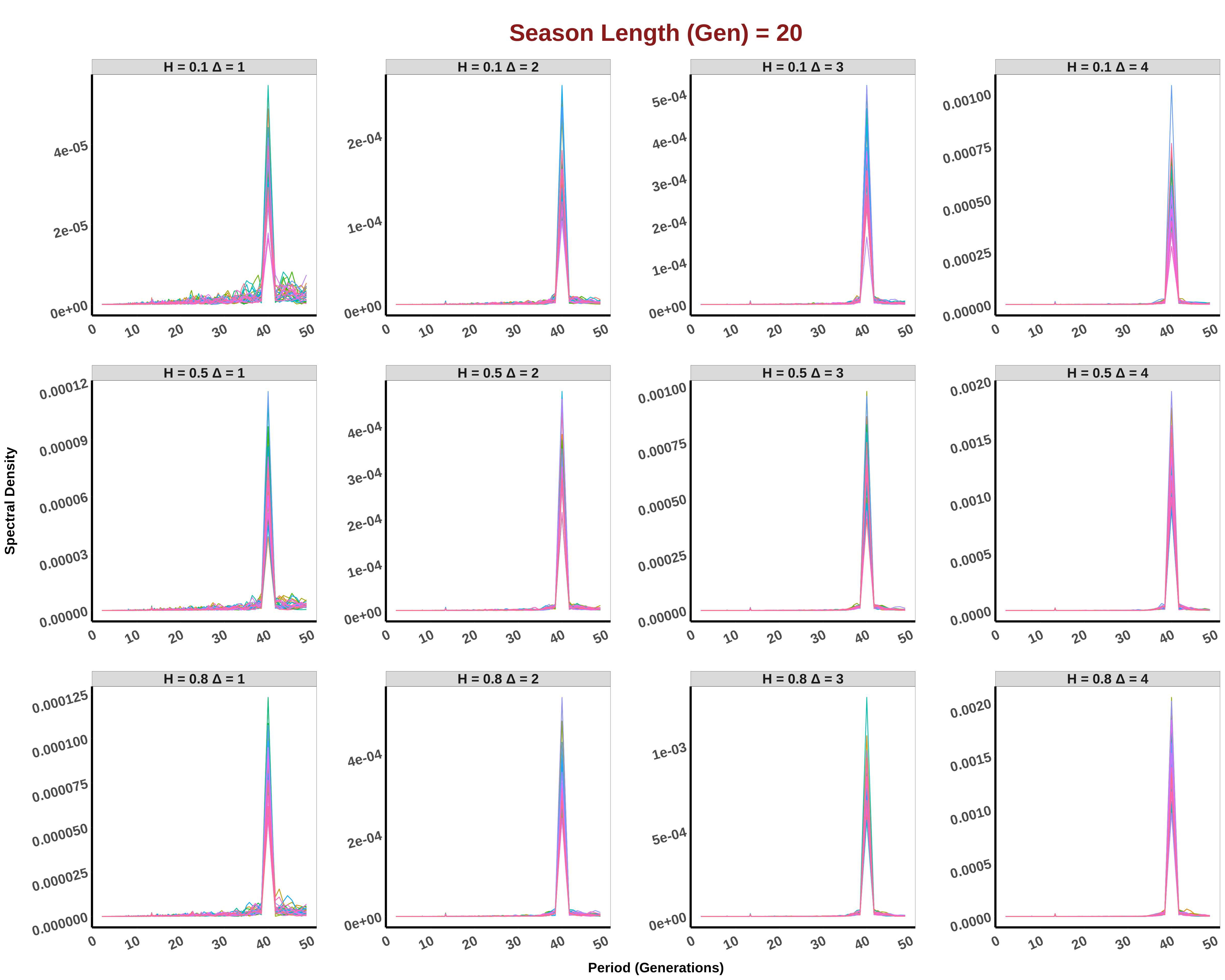

SI 4.2.4.2. Spectral density across 30 replicates for polygenic trait (300 loci) with instantaneous selection model, all parameter combinations with selection length of 20 generations. Each peak represents a full cycle of selection period.

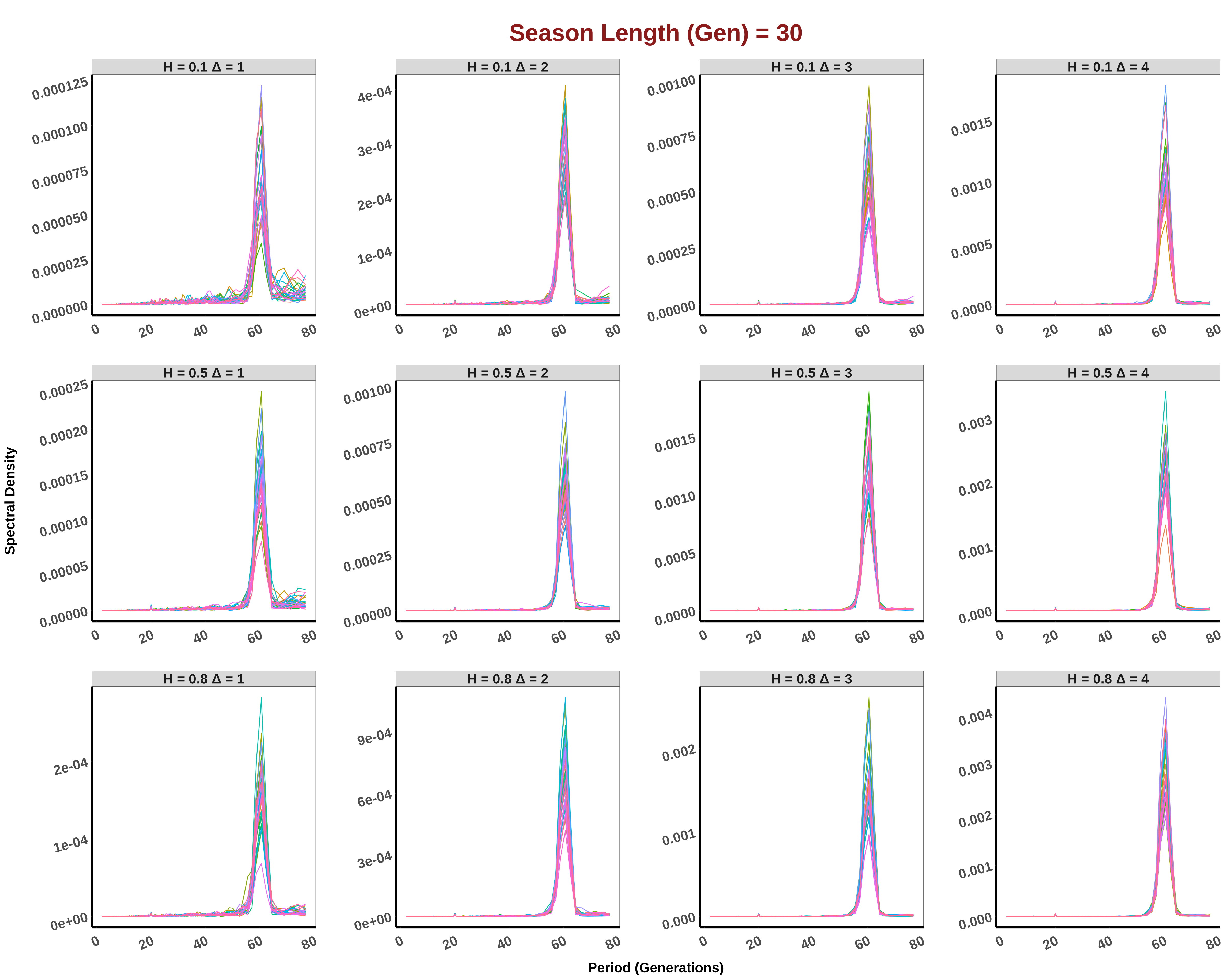

SI 4.2.4.3. Spectral density across 30 replicates for polygenic trait (300 loci) with instantaneous selection model, all parameter combinations with selection length of 30 generations. Each peak represents a full cycle of selection period.

### 5. Fixed percentages for each architecture

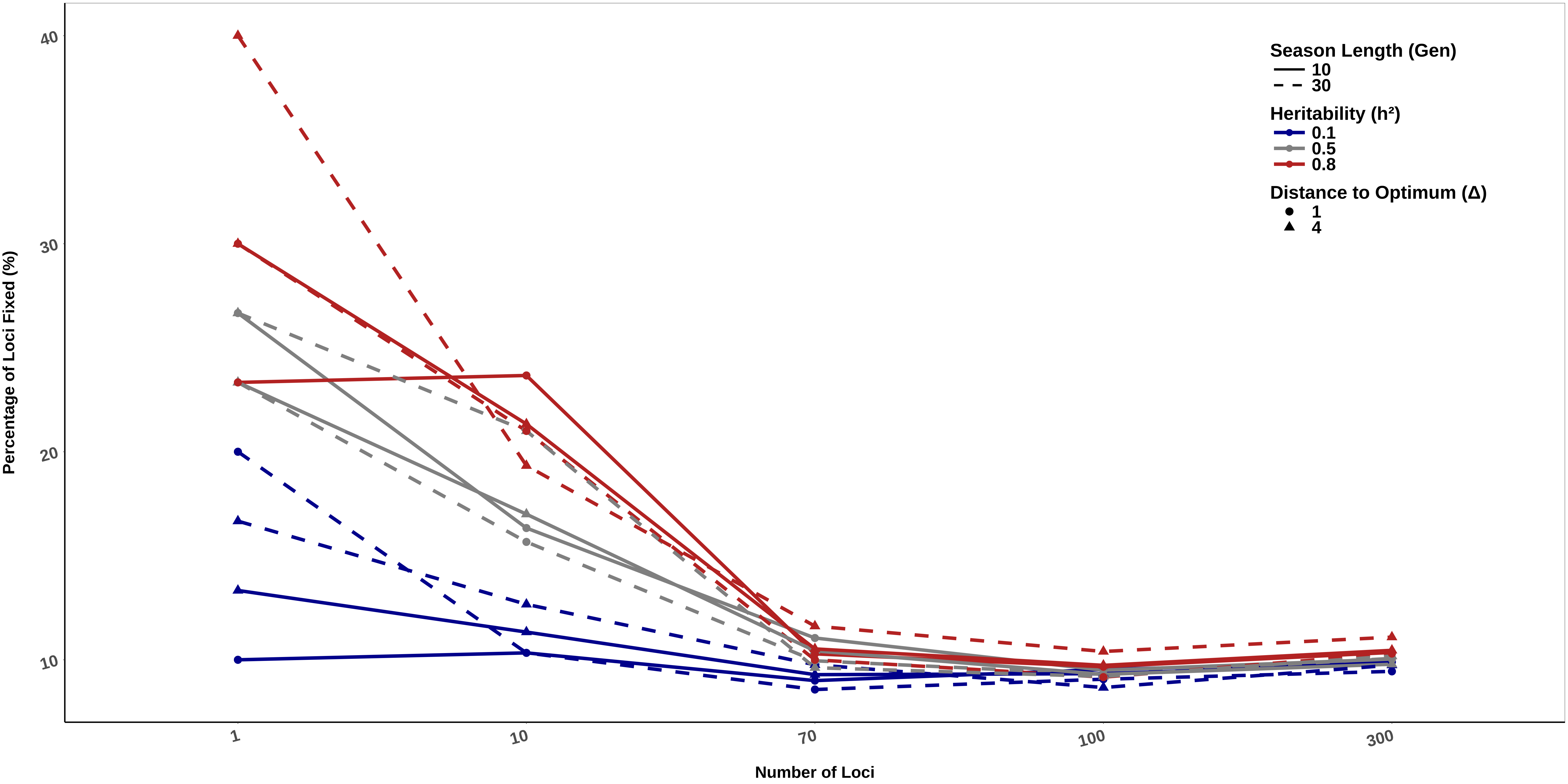

### 6. Sampling Points for spectral analysis

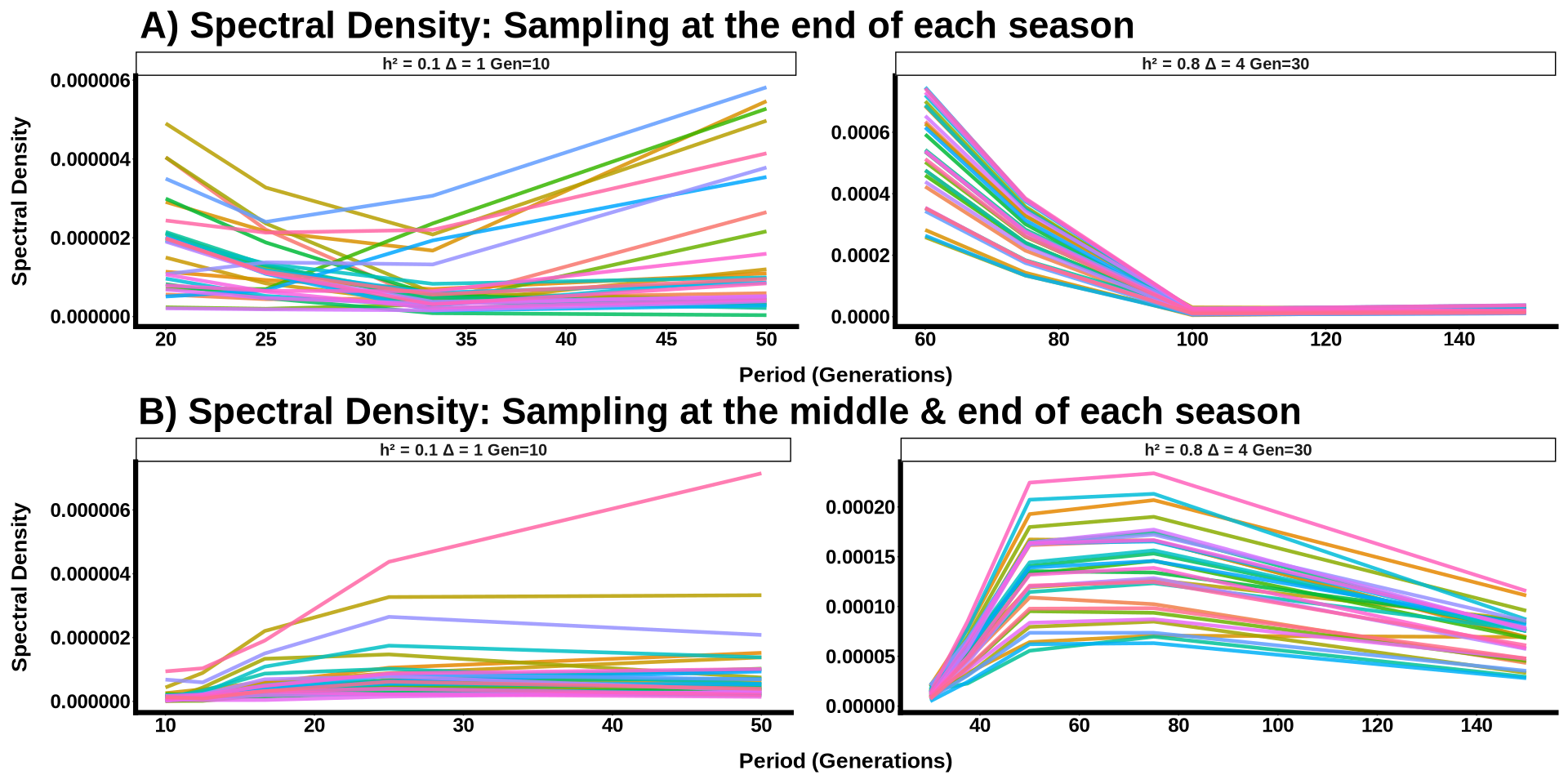

SI 6. Spectral analysis of instantaneous selection for a 100 loci model using 10 sampling points to compare the identification of spectral peaks under high versus low parameter combinations. Although seasonal signatures of selection can be identified with 10 samples under the high parameter combination, recovering the spectral peak remains challenging under the low parameter combination, which may require increasing the number of sampling points.

### 7. Simple Gradual Model

#### 7.1. Allele frequency all parameters

##### 7.1.1. Monogenic

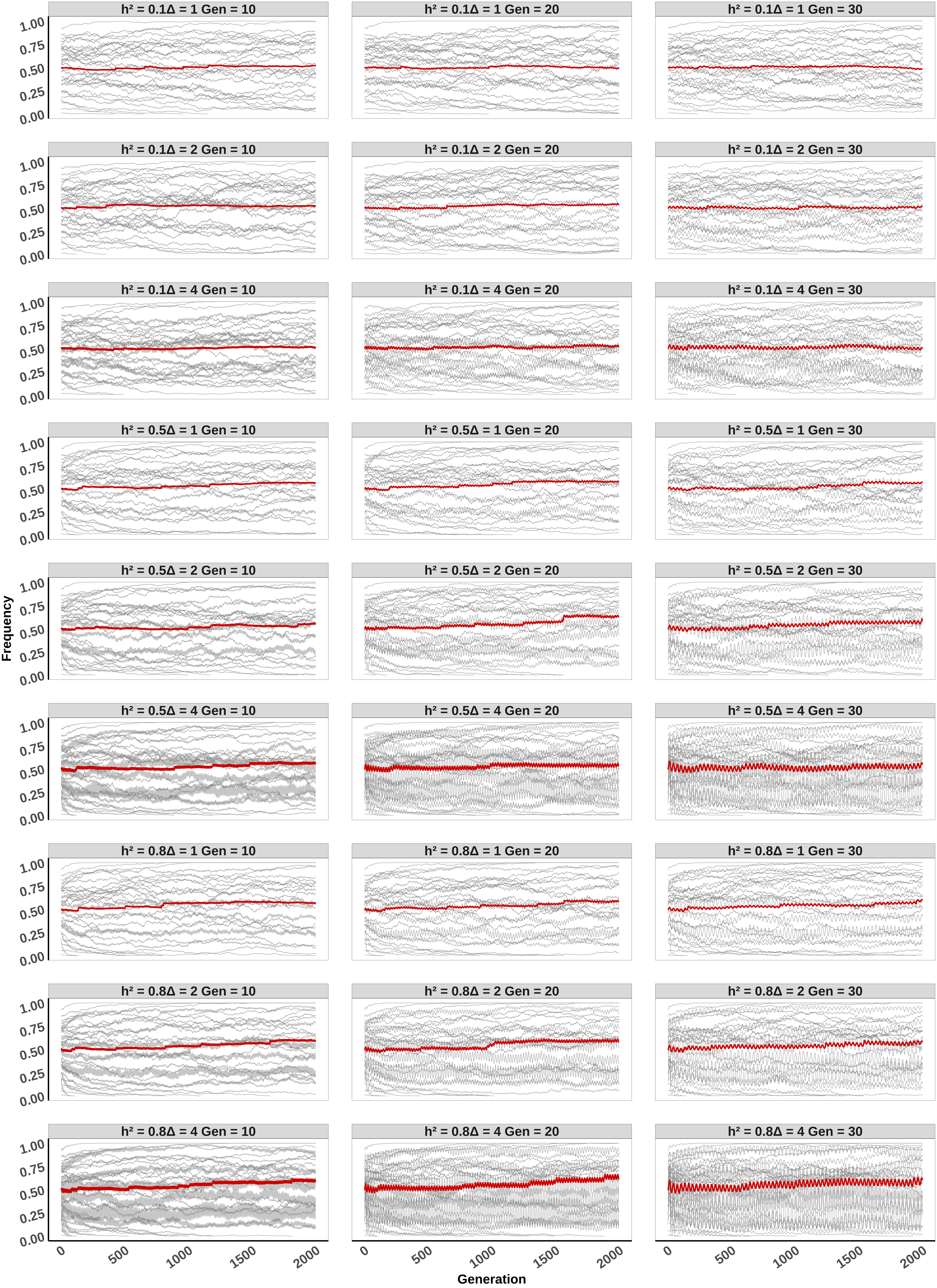

SI 7.1.1. Single locus 30 replicates allele frequency dynamics for gradual two equal seasons selection model, showing all parameter interactions. The highlighted line indicates the mean allele frequency for each generation.

##### 7.1.2. Oligogenic

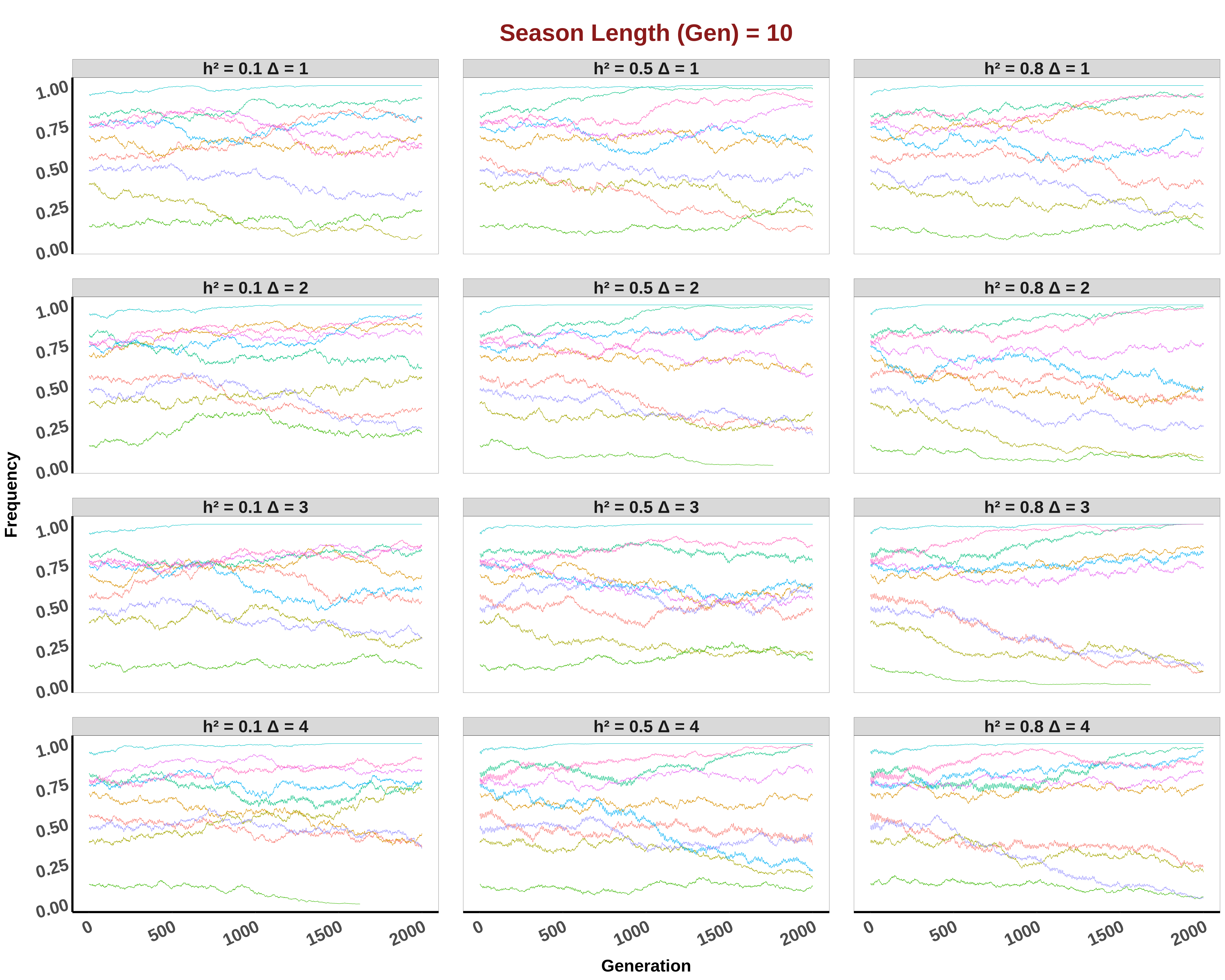

SI 7.1.2.1. Allele frequency dynamics for all parameter combinations with selection length of 10 generations from a single replicate, gradual two equal seasons selection model. Each line represents the frequency trajectory for a locus from oligogenic trait (10 loci).

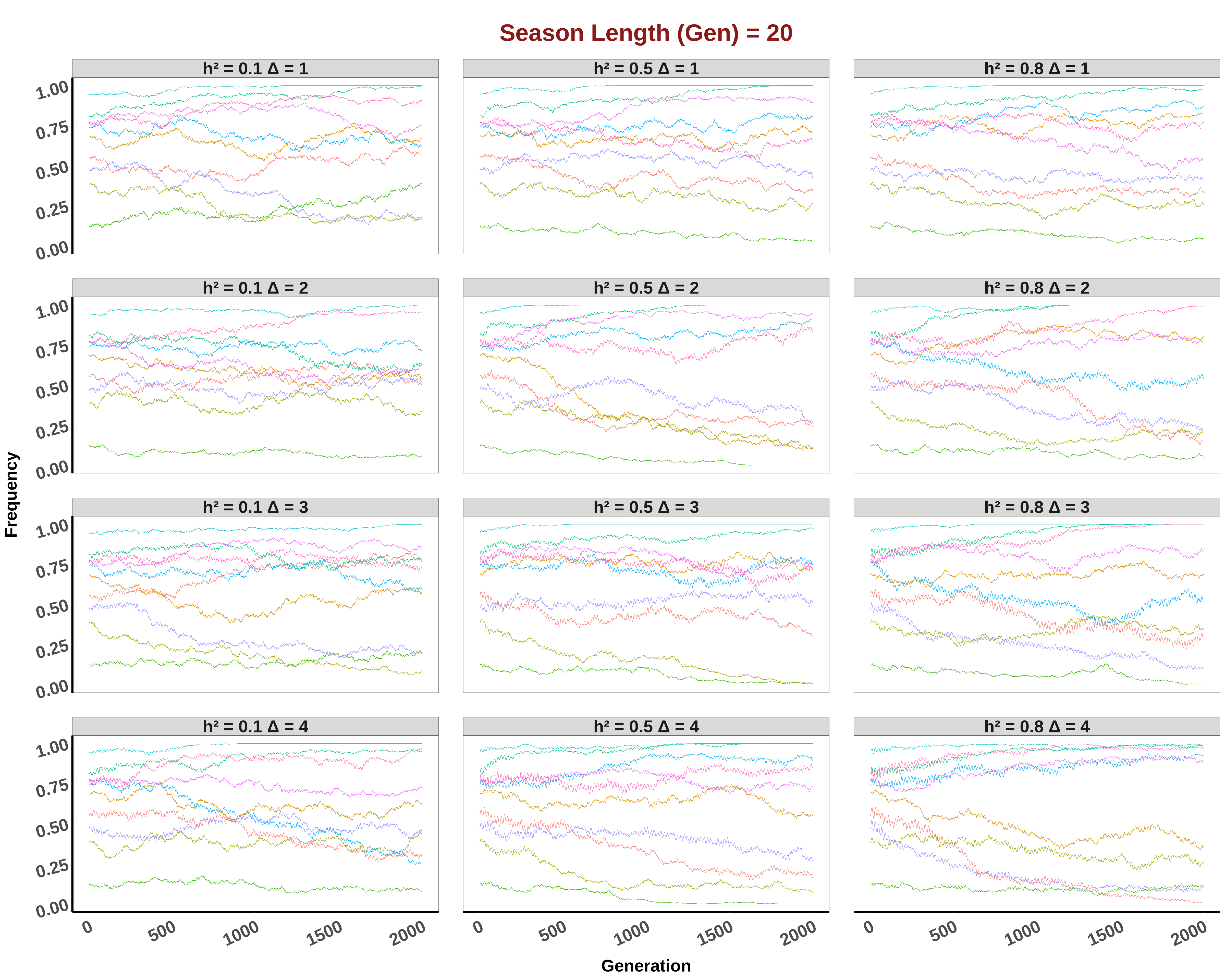

SI 7.1.2.2. Allele frequency dynamics for all parameter combinations with selection length of 20 generations from a single replicate, gradual two equal seasons selection model. Each line represents the frequency trajectory for a locus from oligogenic trait (10 loci).

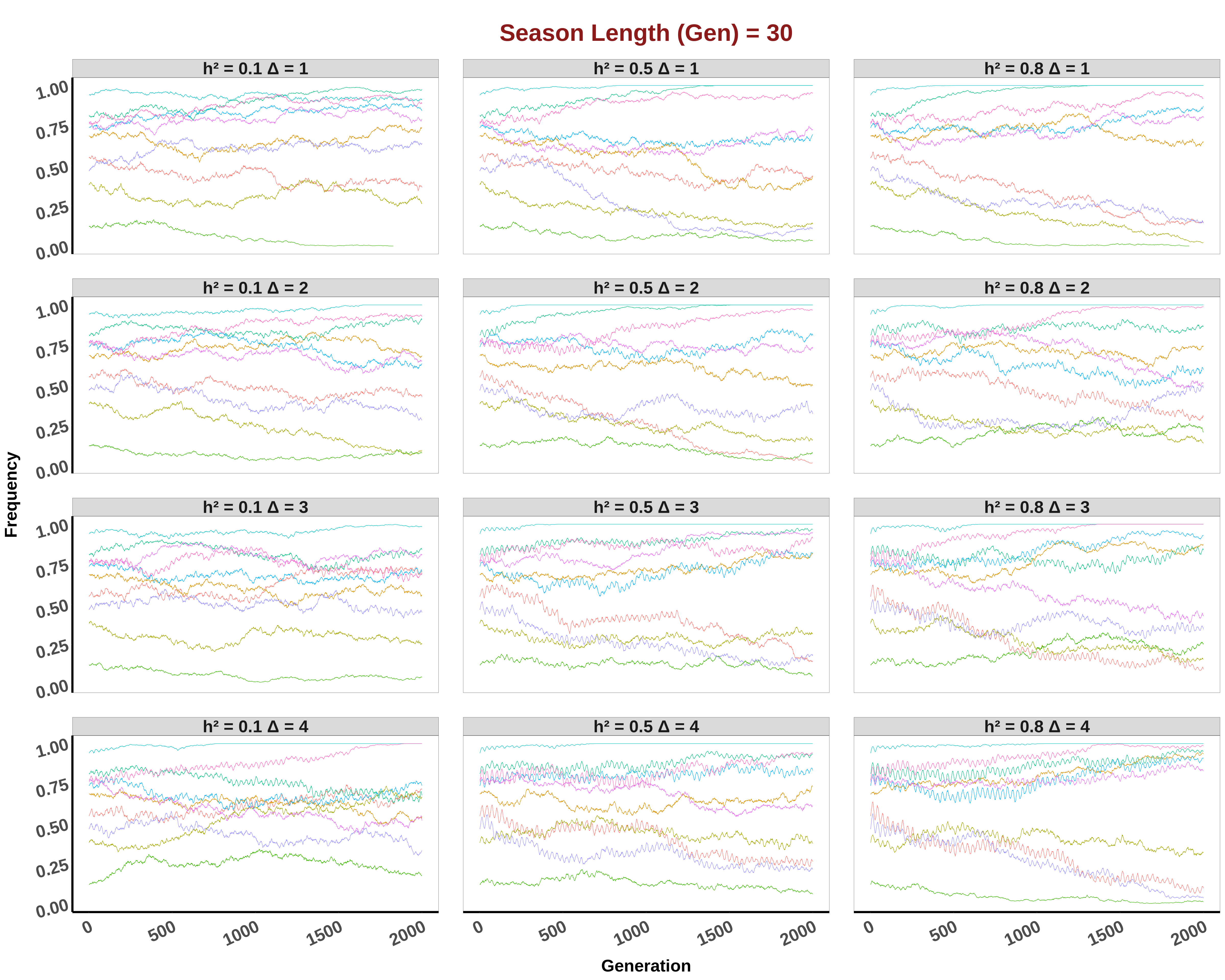

SI 7.1.2.3. Allele frequency dynamics for all parameter combinations with selection length of 30 generations from a single replicate, gradual two equal seasons selection model. Each line represents the frequency trajectory for a locus from oligogenic trait (10 loci).

##### 7.1.3. Polygenic 70 loci

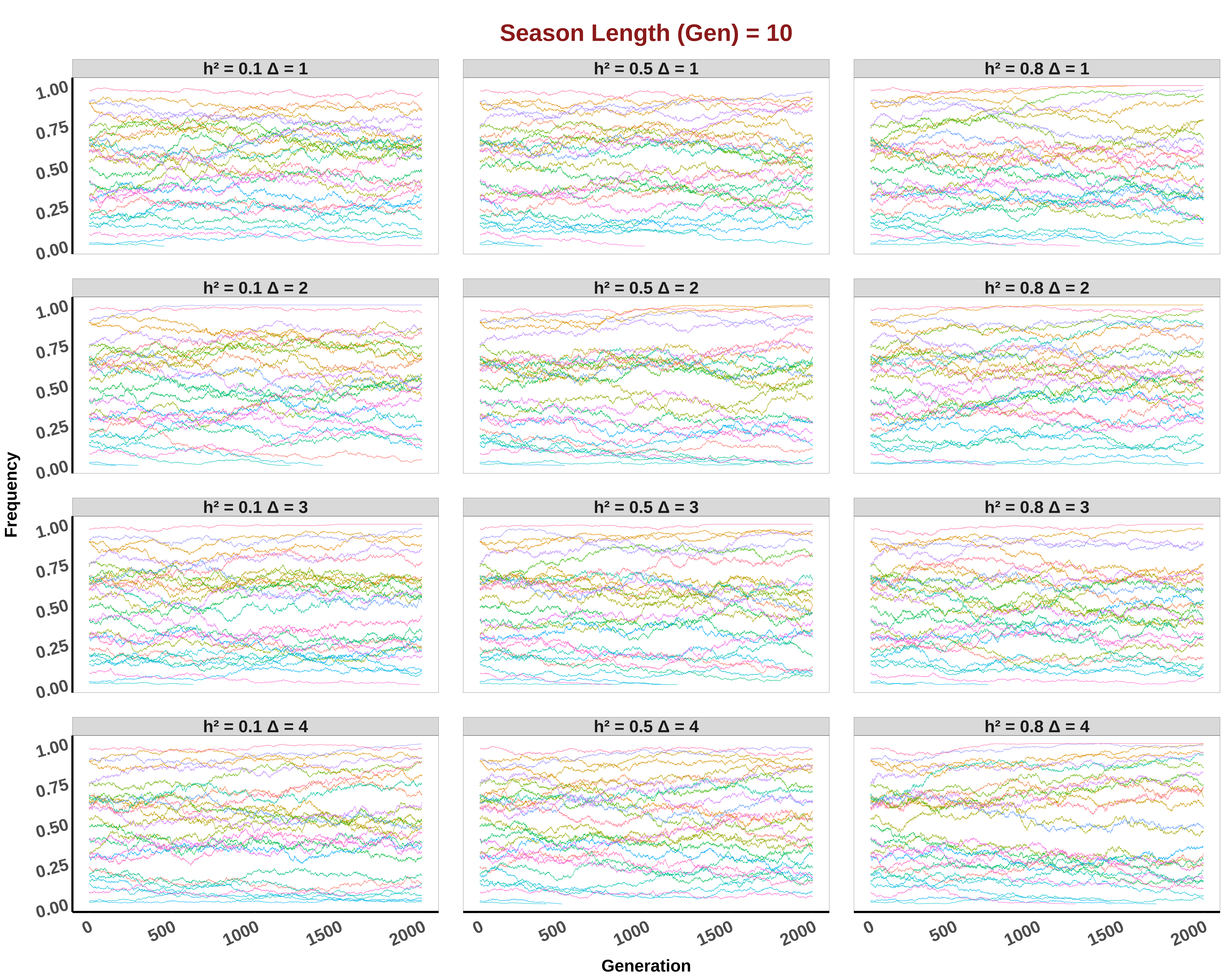

SI 7.1.3.1. Allele frequency dynamics for all parameter combinations with selection length of 10 generations from a single replicate, gradual two equal seasons selection model. Each line represents the frequency trajectory for a locus from polygenic trait (70 loci).

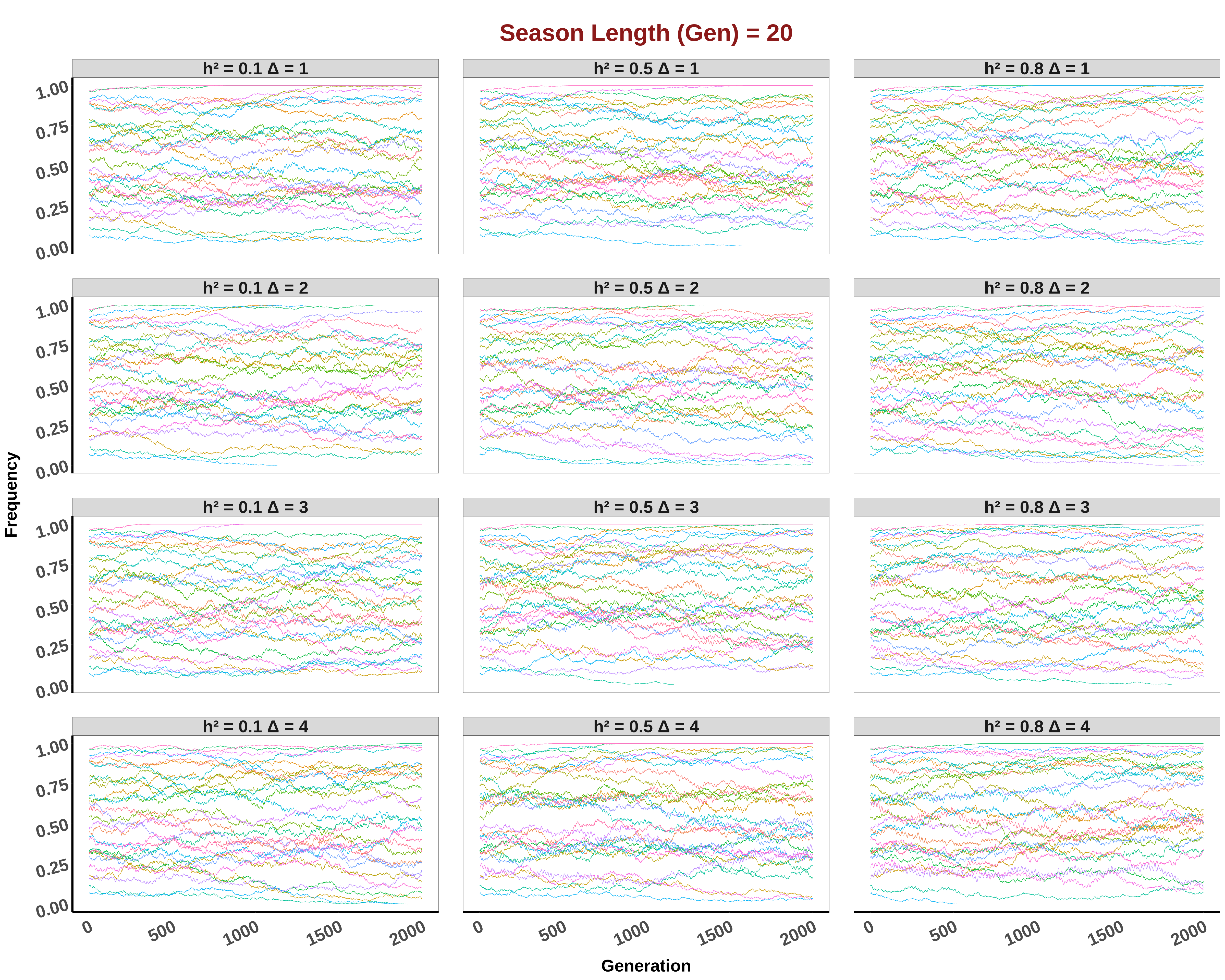

SI 7.1.3.2. Allele frequency dynamics for all parameter combinations with selection length of 20 generations from a single replicate, gradual two equal seasons selection model. Each line represents the frequency trajectory for a locus from polygenic trait (70 loci).

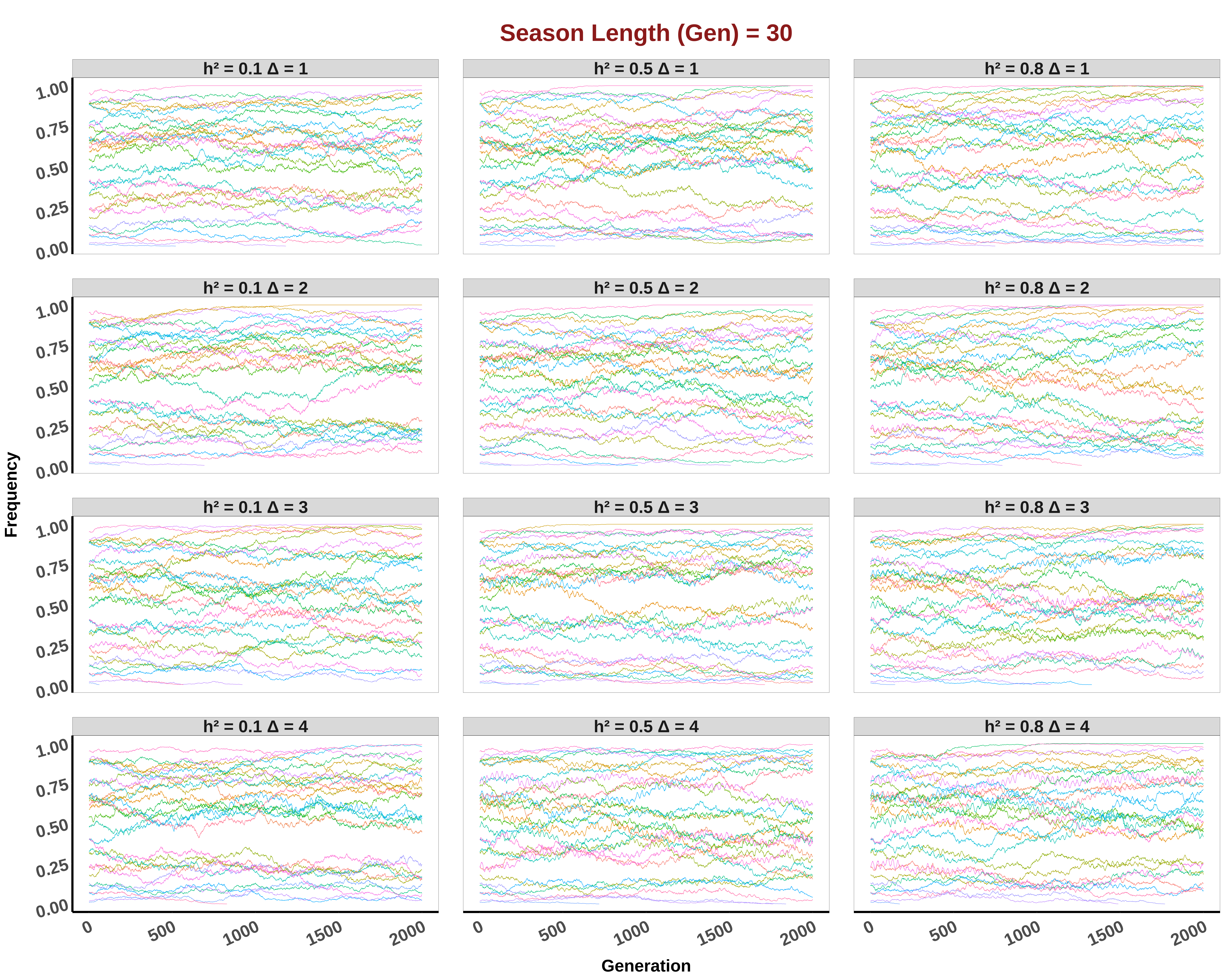

SI 7.1.3.3. Allele frequency dynamics for all parameter combinations with selection length of 30 generations from a single replicate, gradual two equal seasons selection model. Each line represents the frequency trajectory for a locus from polygenic trait (70 loci).

##### 7.1.4. Polygenic 300 loci

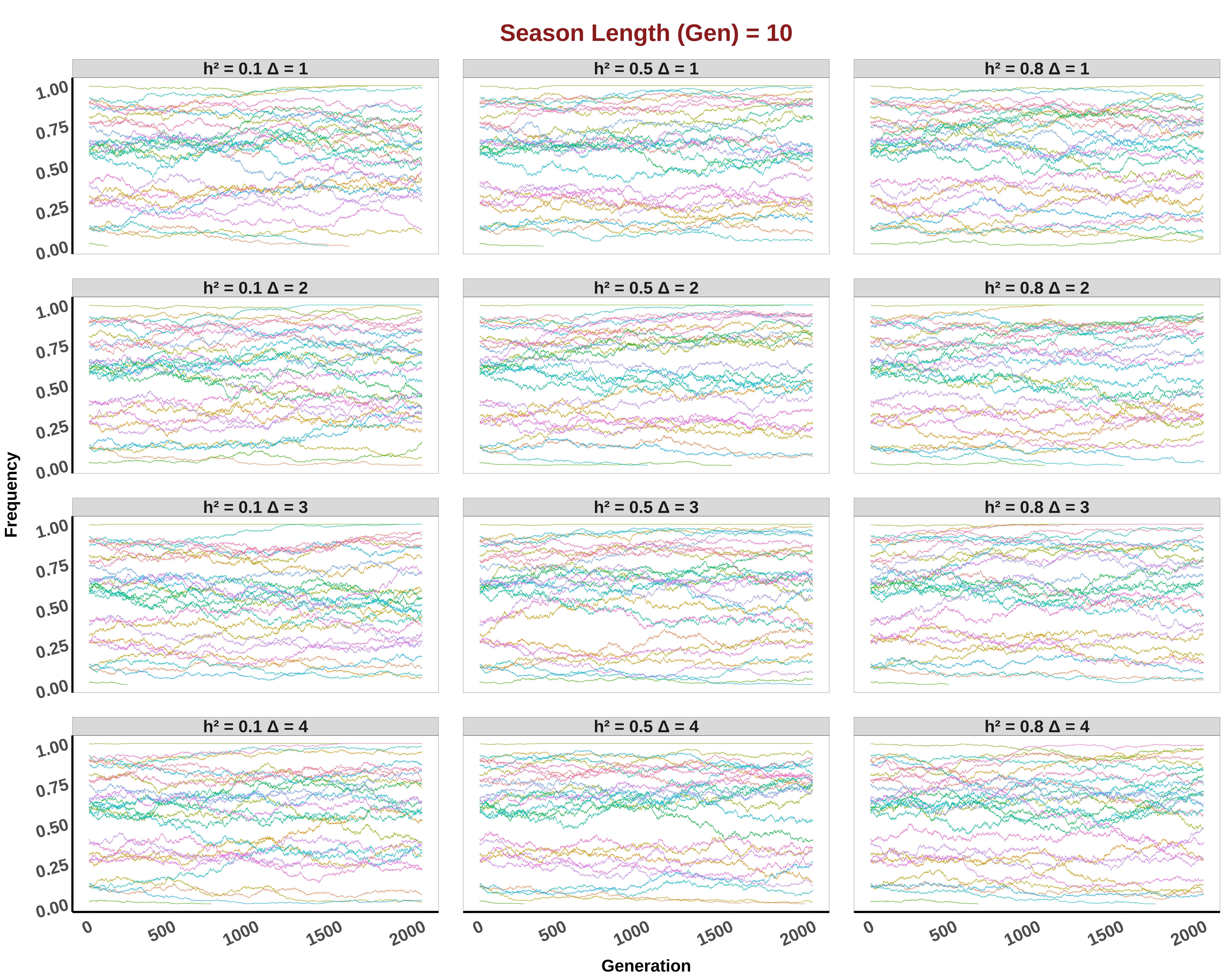

SI 7.1.4.1. Allele frequency dynamics for all parameter combinations with selection length of 10 generations from a single replicate, gradual two equal seasons selection model. Each line represents the frequency trajectory for a locus from polygenic trait (300 loci).

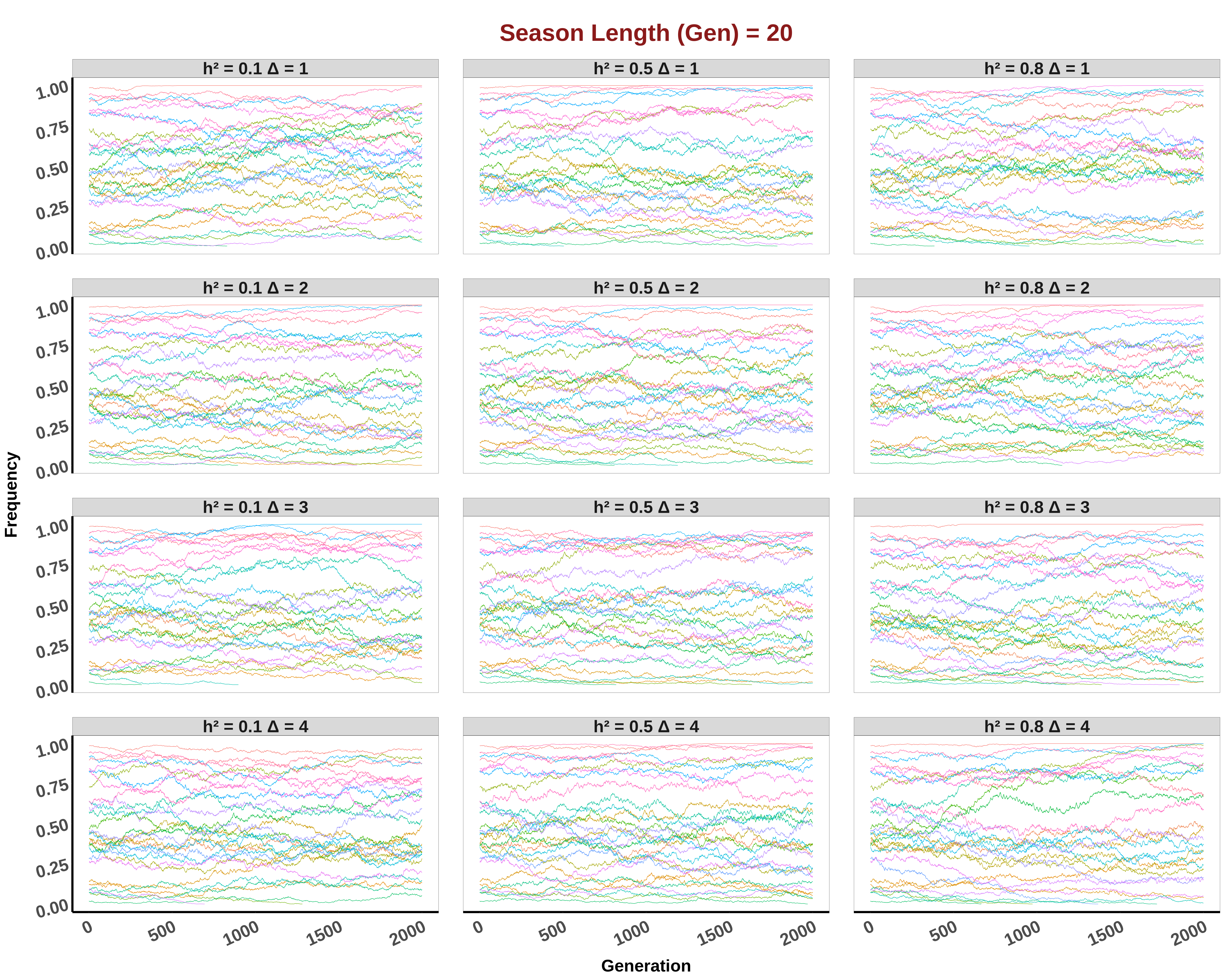

SI 7.1.4.2. Allele frequency dynamics for all parameter combinations with selection length of 20 generations from a single replicate, gradual two equal seasons selection model. Each line represents the frequency trajectory for a locus from polygenic trait (300 loci).

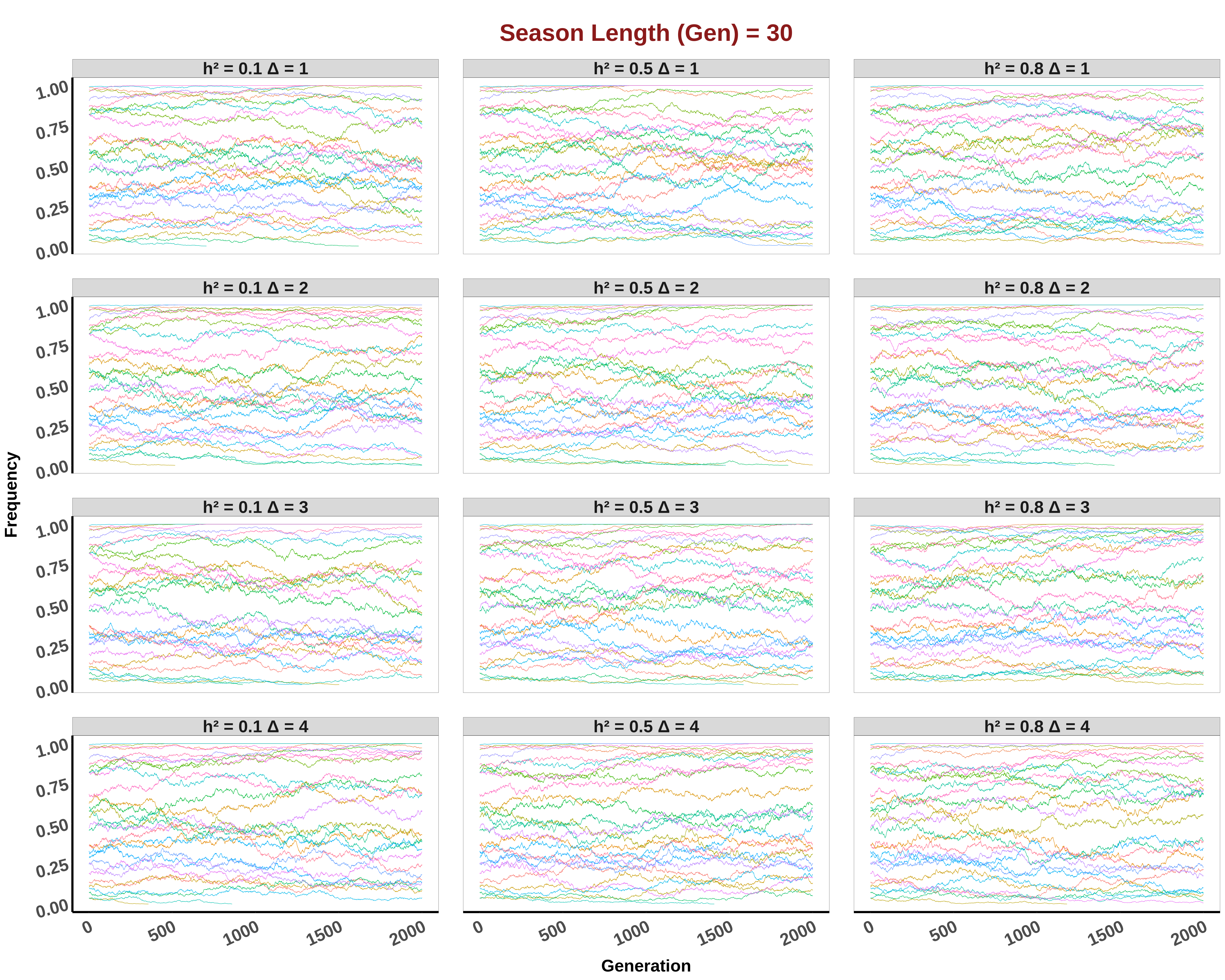

SI 7.1.4.3. Allele frequency dynamics for all parameter combinations with selection length of 30 generations from a single replicate, gradual two equal seasons selection model. Each line represents the frequency trajectory for a locus from polygenic trait (300 loci).

#### 7.2. Spectral analysis for all parameters

##### 7.2.1. Monogenic

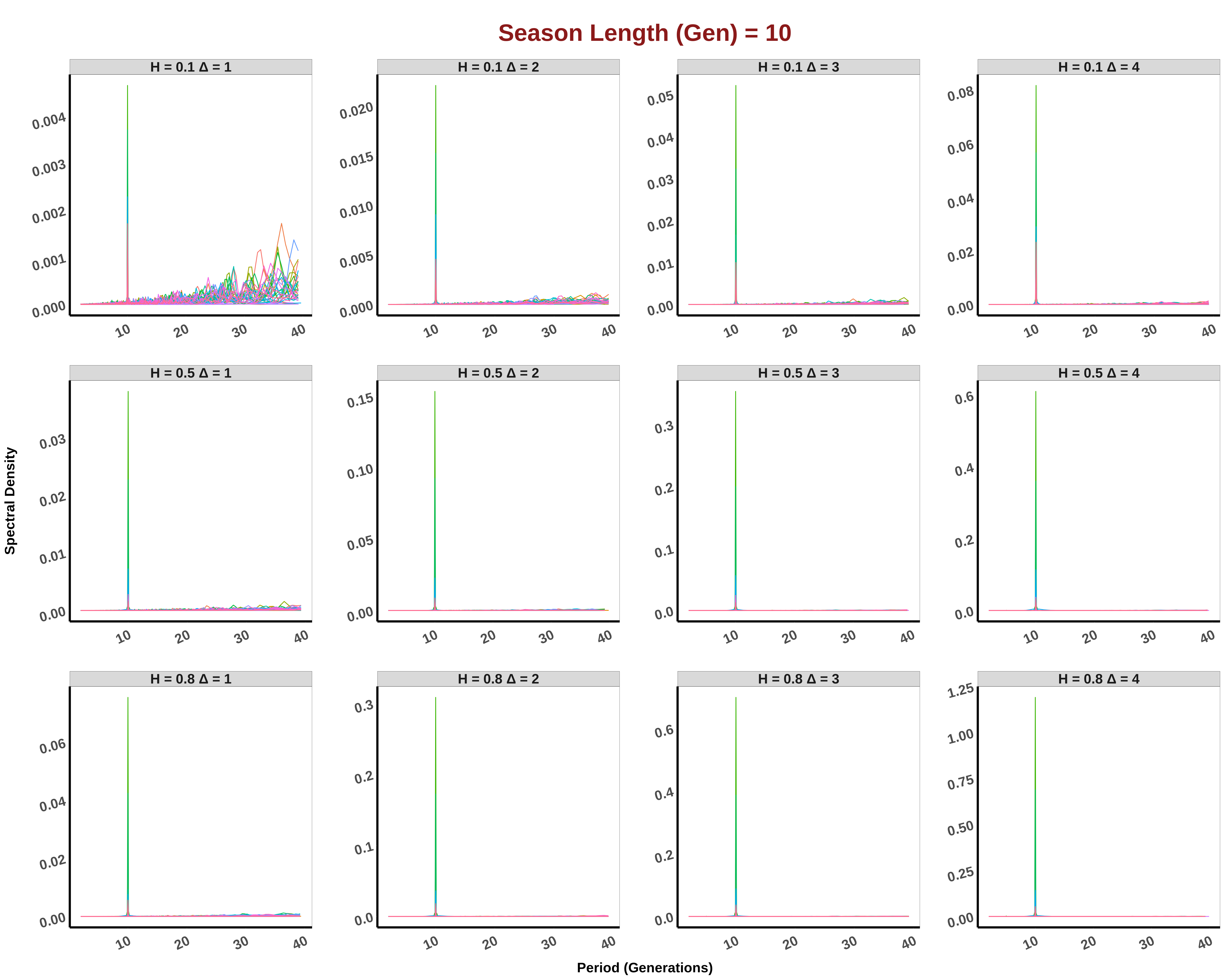

SI 7.2.1.1. Spectral density across 30 replicates for monogenic trait (1 locus) with gradual optima change two equal seasons, all parameter combinations with selection length of 10 generations. Each peak represents a full cycle of selection period.

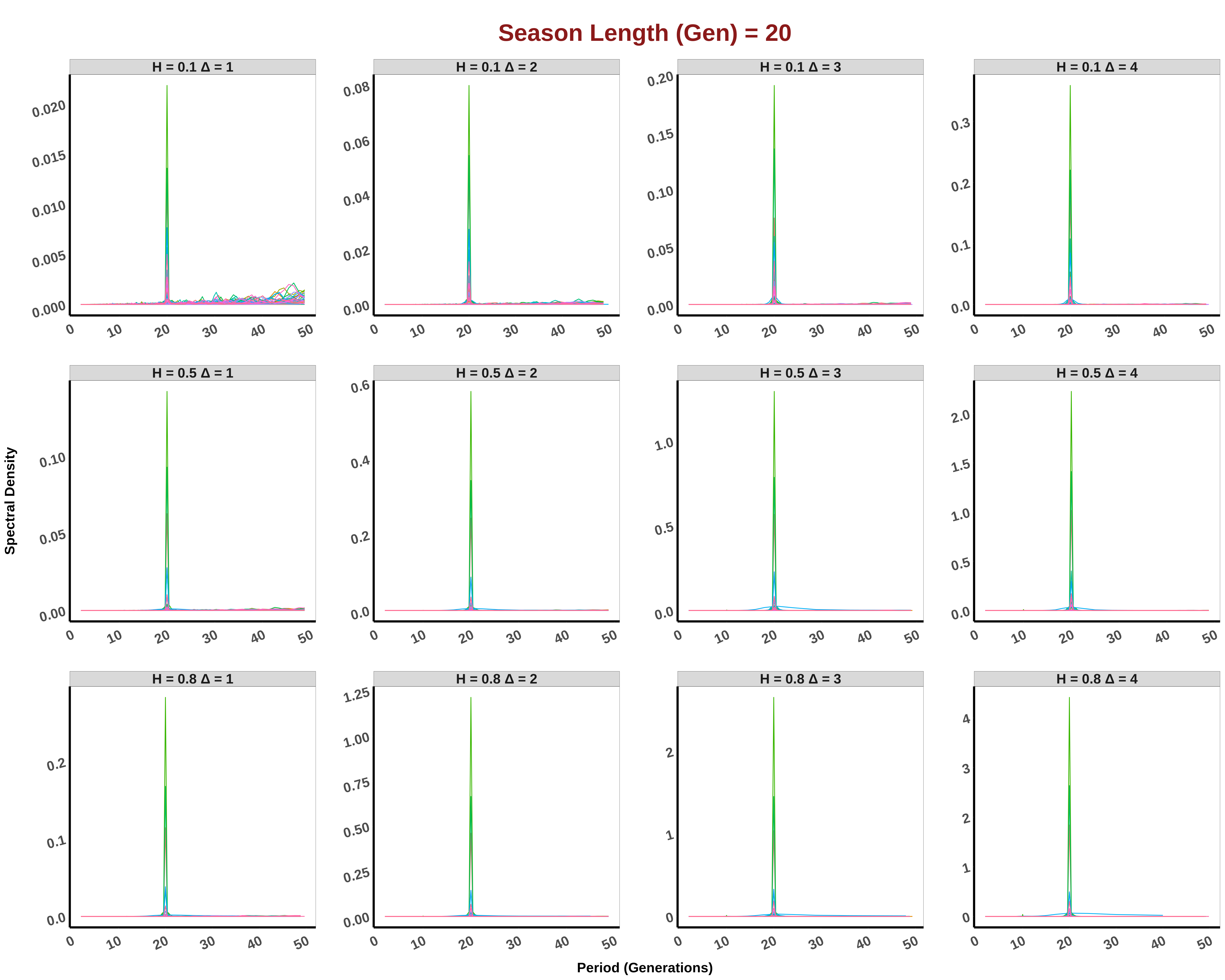

SI 7.2.1.2. Spectral density across 30 replicates for monogenic trait (1 locus) with gradual optima change two equal seasons, all parameter combinations with selection length of 20 generations. Each peak represents a full cycle of selection period.

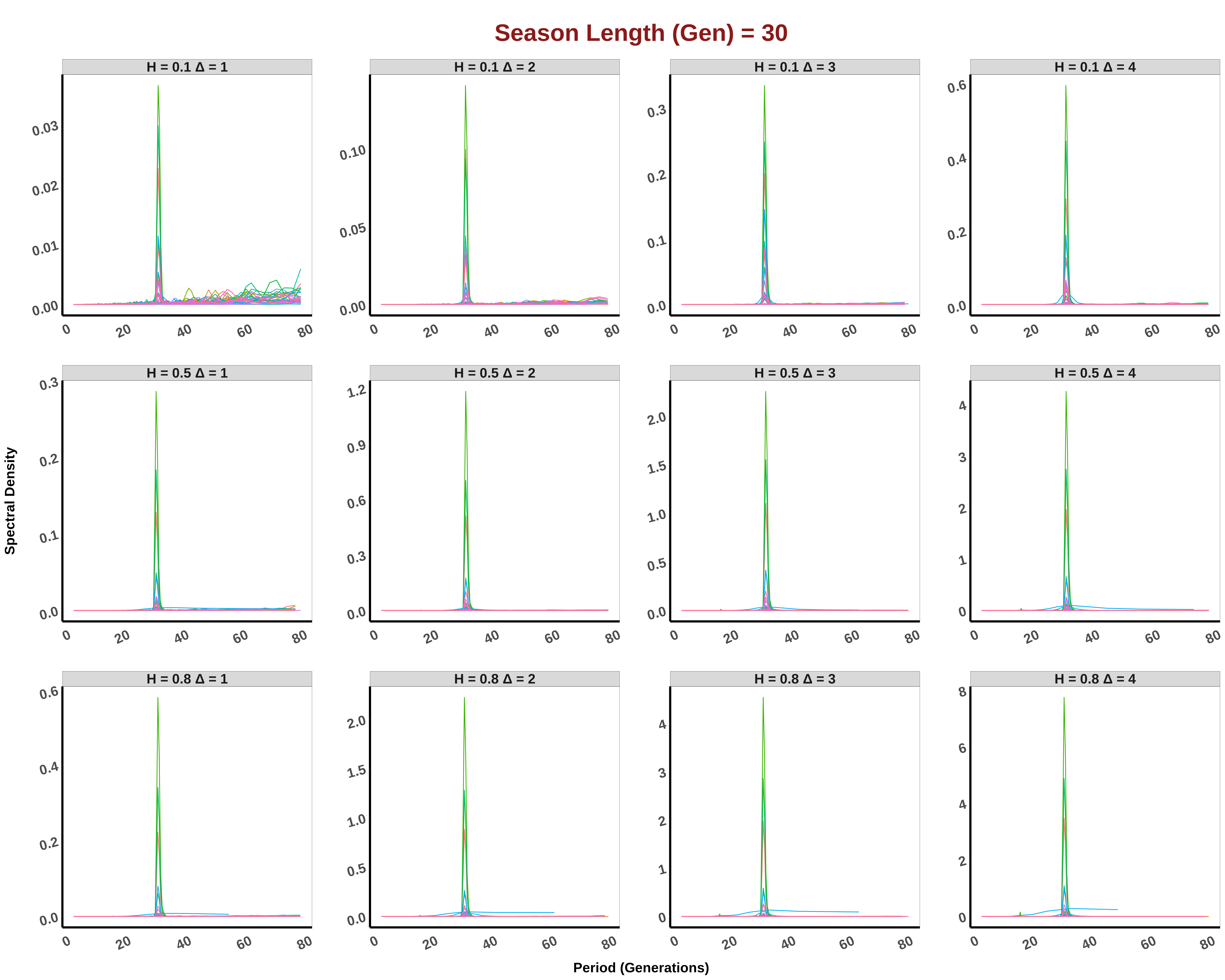

SI 7.2.1.3. Spectral density across 30 replicates for monogenic trait (1 locus) with gradual optima change two equal seasons, all parameter combinations with selection length of 30 generations. Each peak represents a full cycle of selection period.

##### 7.2.2. Oligogenic

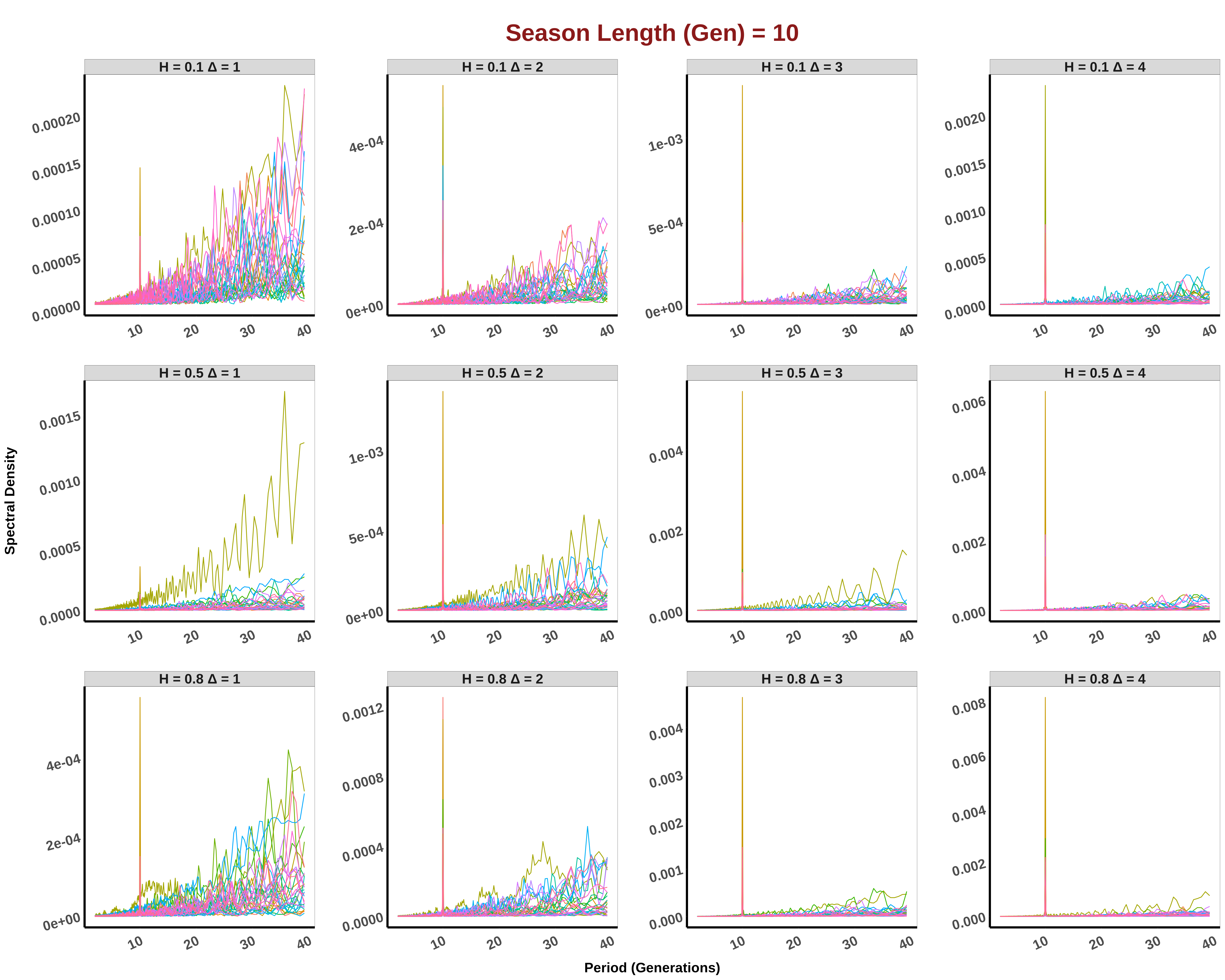

SI 7.2.2.1. Spectral density across 30 replicates for oligogenic trait (10 loci) with gradual optima change two equal seasons, all parameter combinations with selection length of 10 generations. Each peak represents a full cycle of selection period.

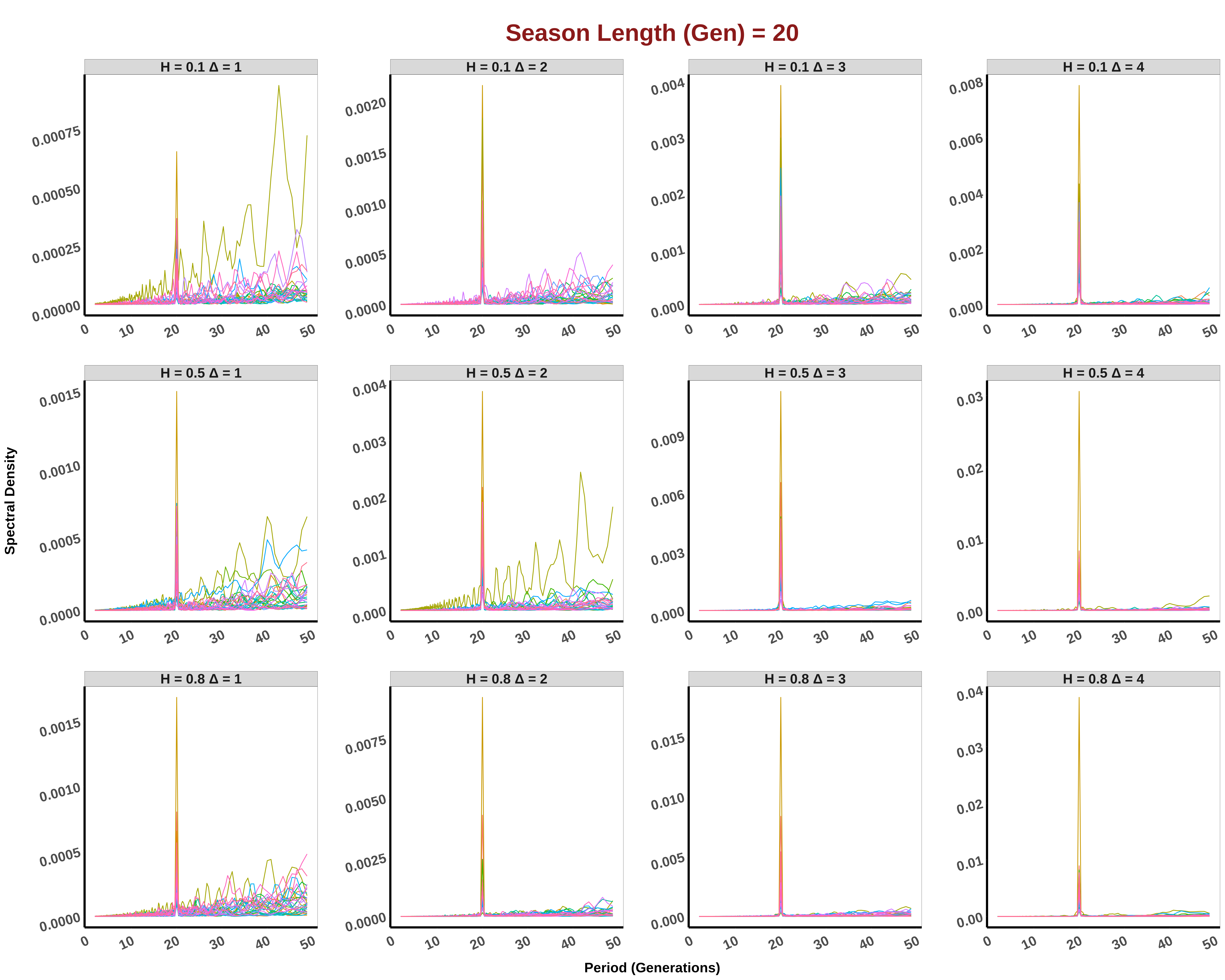

SI 7.2.2.2. Spectral density across 30 replicates for oligogenic trait (10 loci) with gradual optima change two equal seasons, all parameter combinations with selection length of 20 generations. Each peak represents a full cycle of selection period.

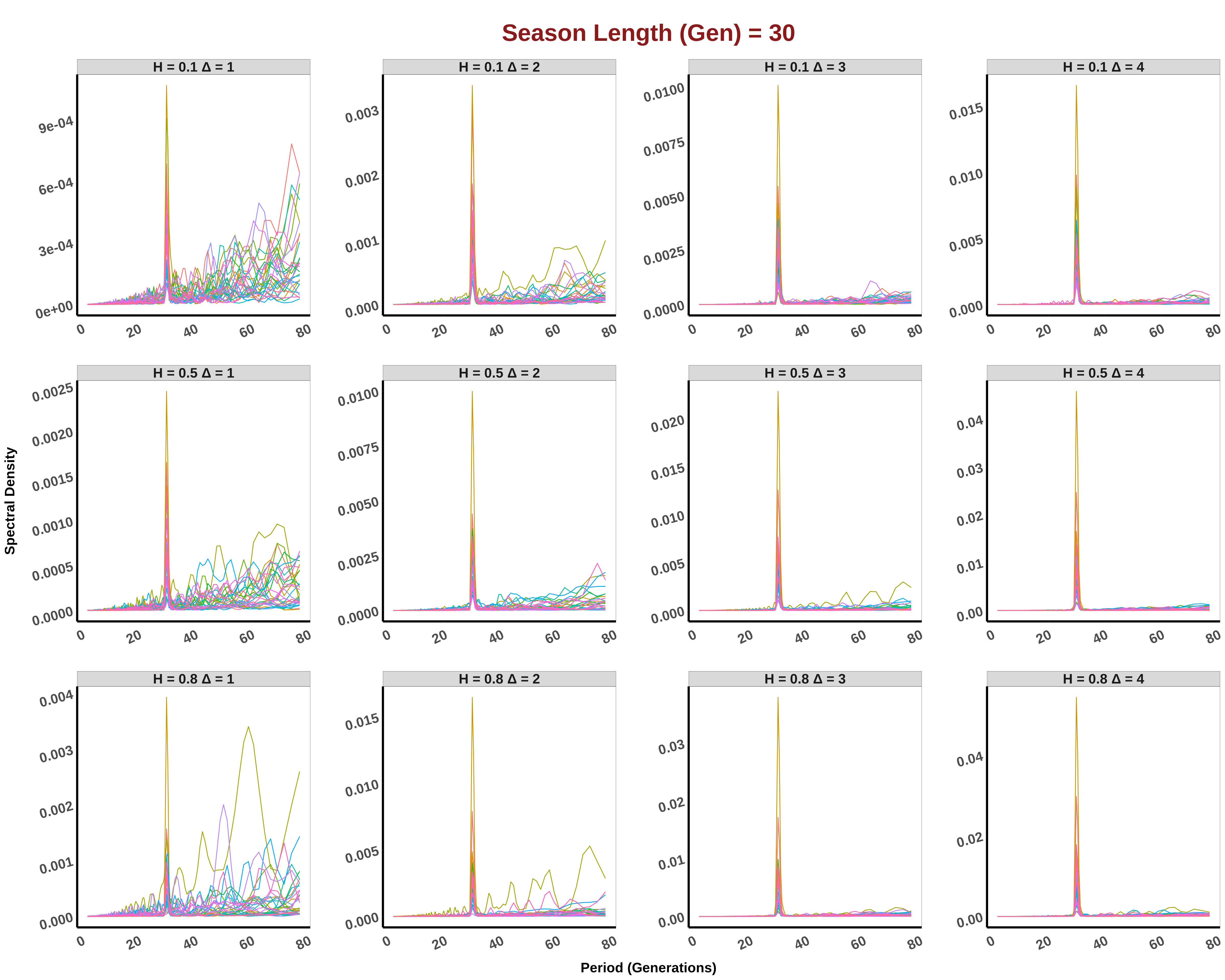

SI 7.2.2.3. Spectral density across 30 replicates for oligogenic trait (10 loci) with gradual optima change two equal seasons, all parameter combinations with selection length of 30 generations. Each peak represents a full cycle of selection period.

##### 7.2.3. Polygenic 70

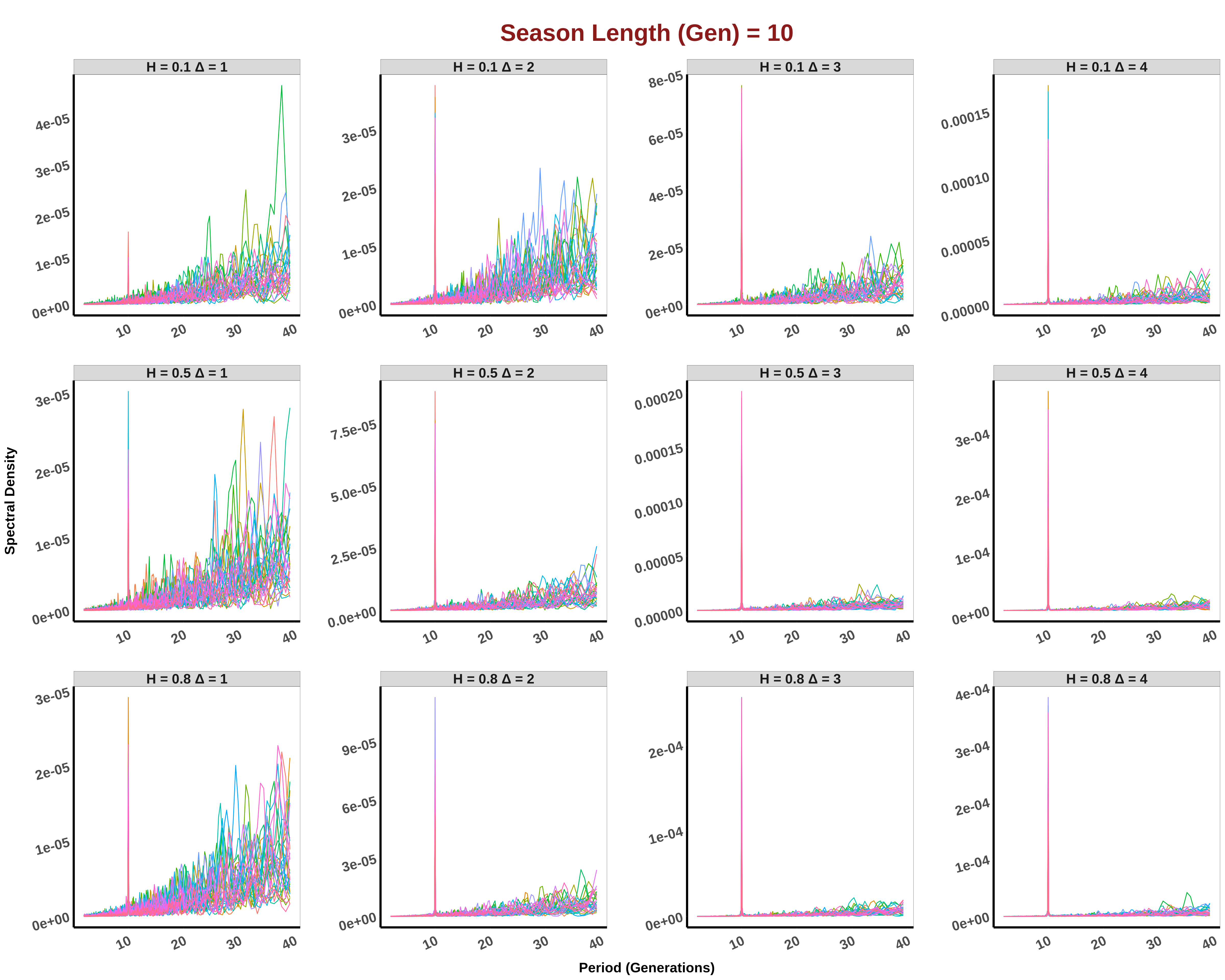

SI 7.2.3.1. Spectral density across 30 replicates for polygenic trait (70 loci) with gradual optima change two equal seasons, all parameter combinations with selection length of 10 generations. Each peak represents a full cycle of selection period.

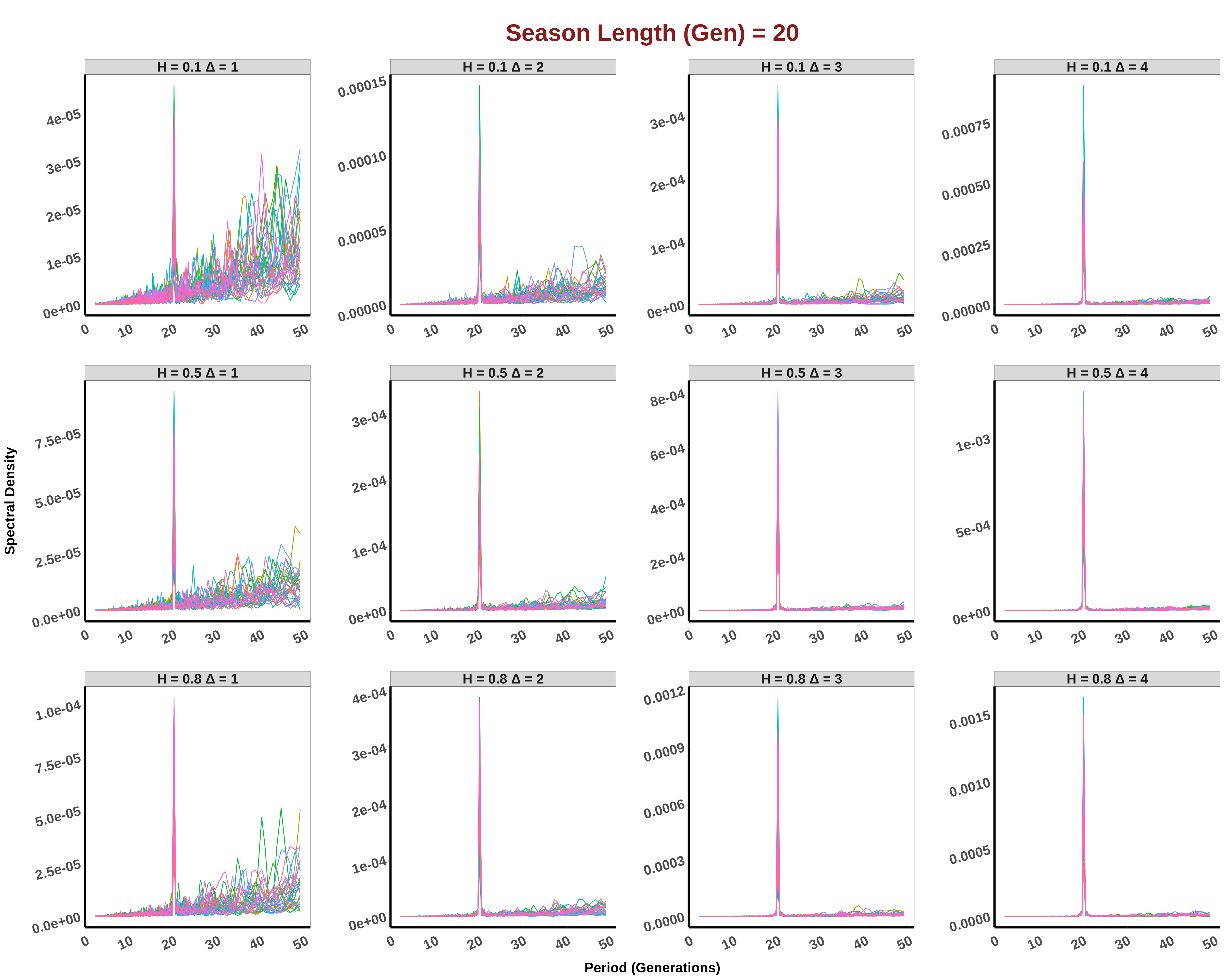

SI 7.2.3.2. Spectral density across 30 replicates for polygenic trait (70 loci) with gradual optima change two equal seasons, all parameter combinations with selection length of 20 generations. Each peak represents a full cycle of selection period.

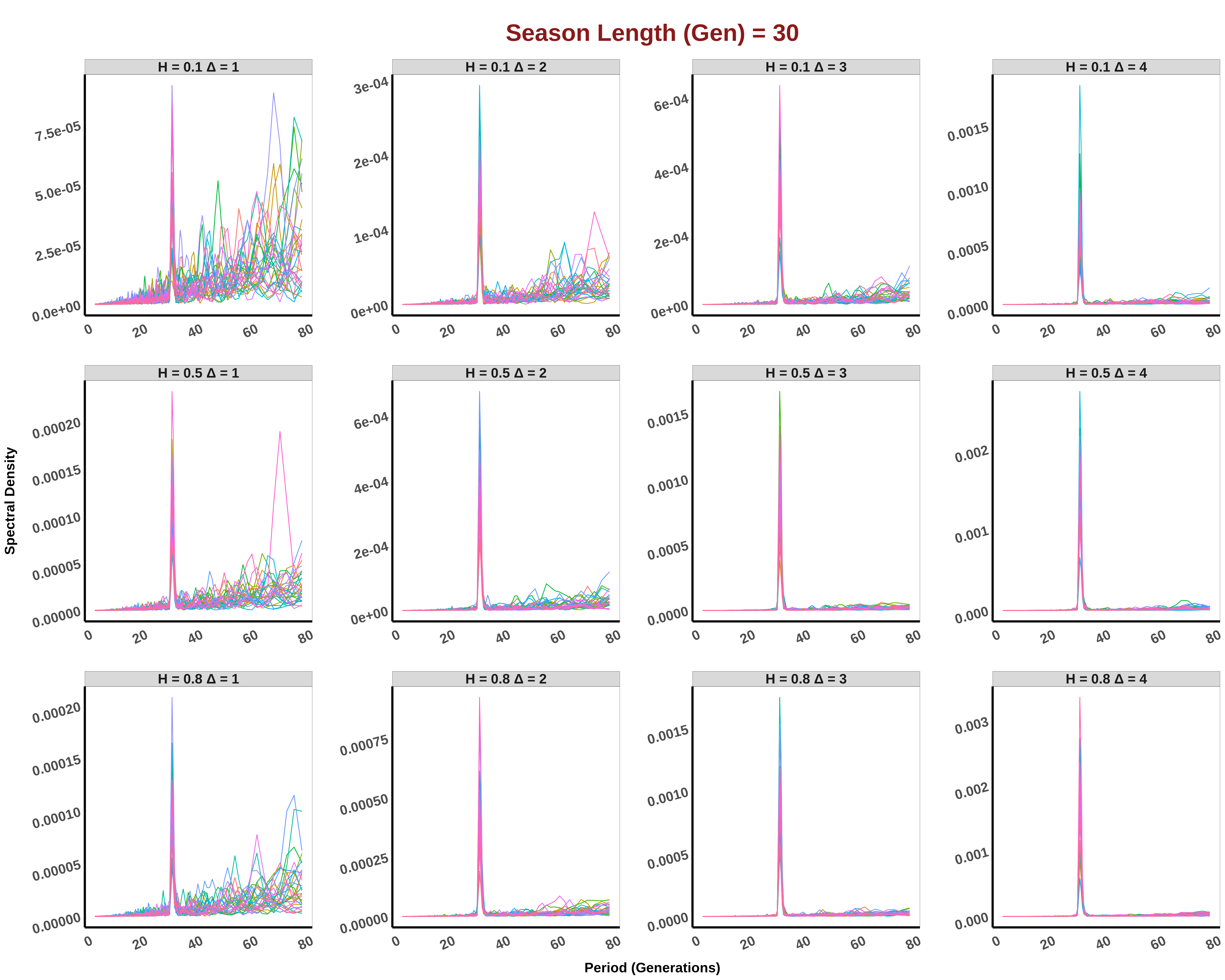

SI 7.2.3.3. Spectral density across 30 replicates for polygenic trait (70 loci) with gradual optima change two equal seasons, all parameter combinations with selection length of 30 generations. Each peak represents a full cycle of selection period.

##### 7.2.4. Polygenic 300

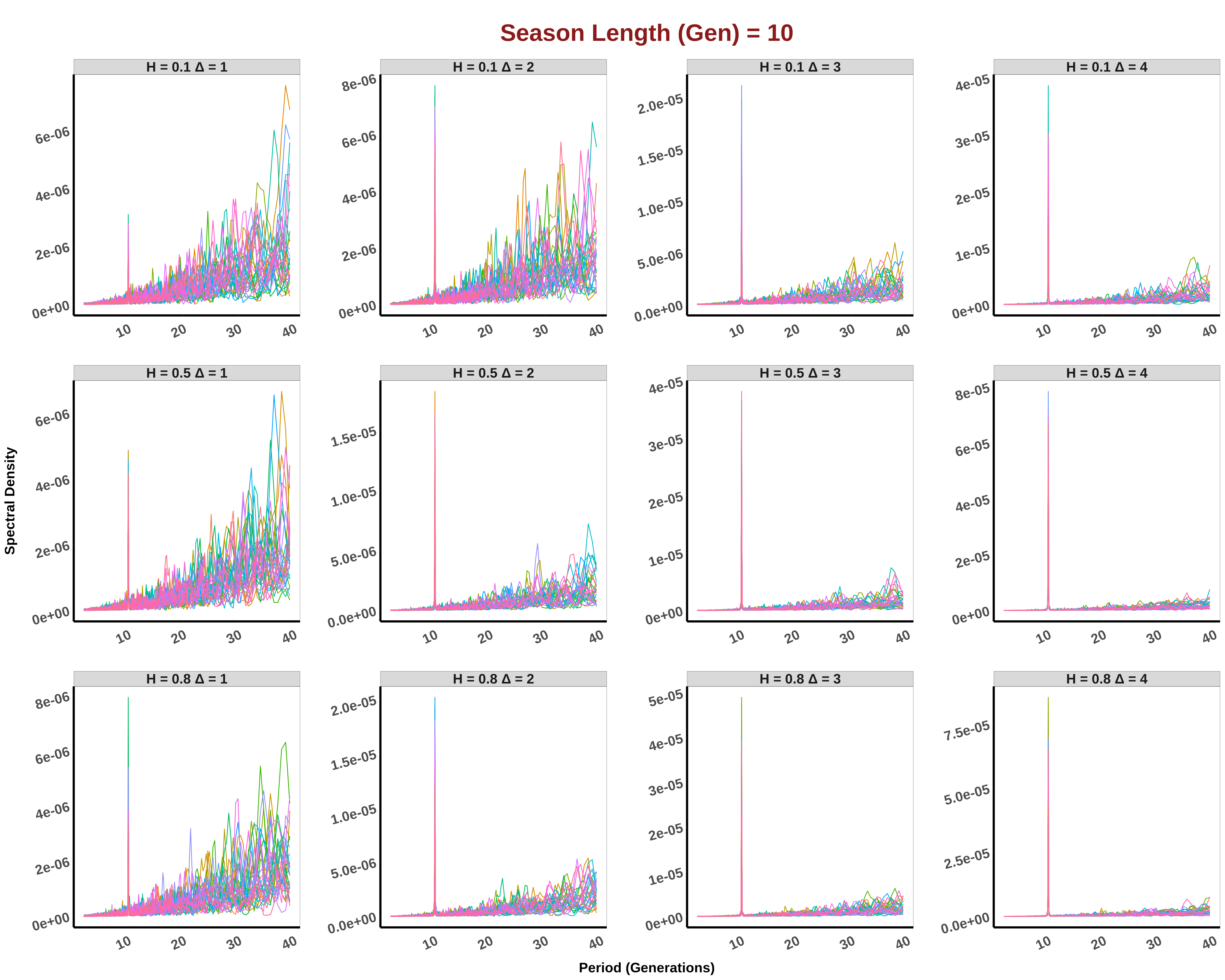

SI 7.2.4.1. Spectral density across 30 replicates for polygenic trait (300 loci) with gradual optima change two equal seasons, all parameter combinations with selection length of 10 generations. Each peak represents a full cycle of selection period.

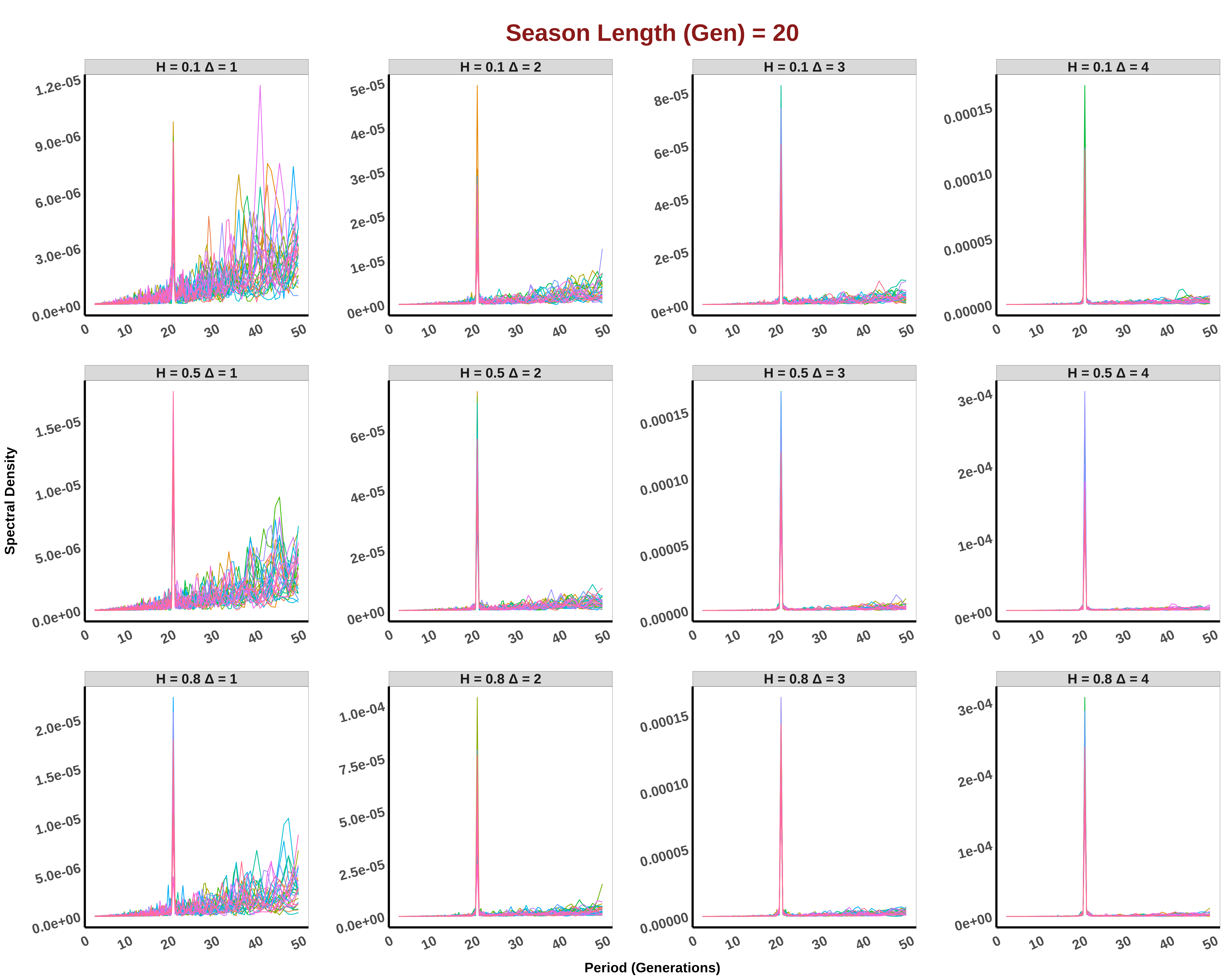

SI 2.2.4.2. Spectral density across 30 replicates for polygenic trait (300 loci) with gradual optima change two equal seasons, all parameter combinations with selection length of 20 generations. Each peak represents a full cycle of selection period.

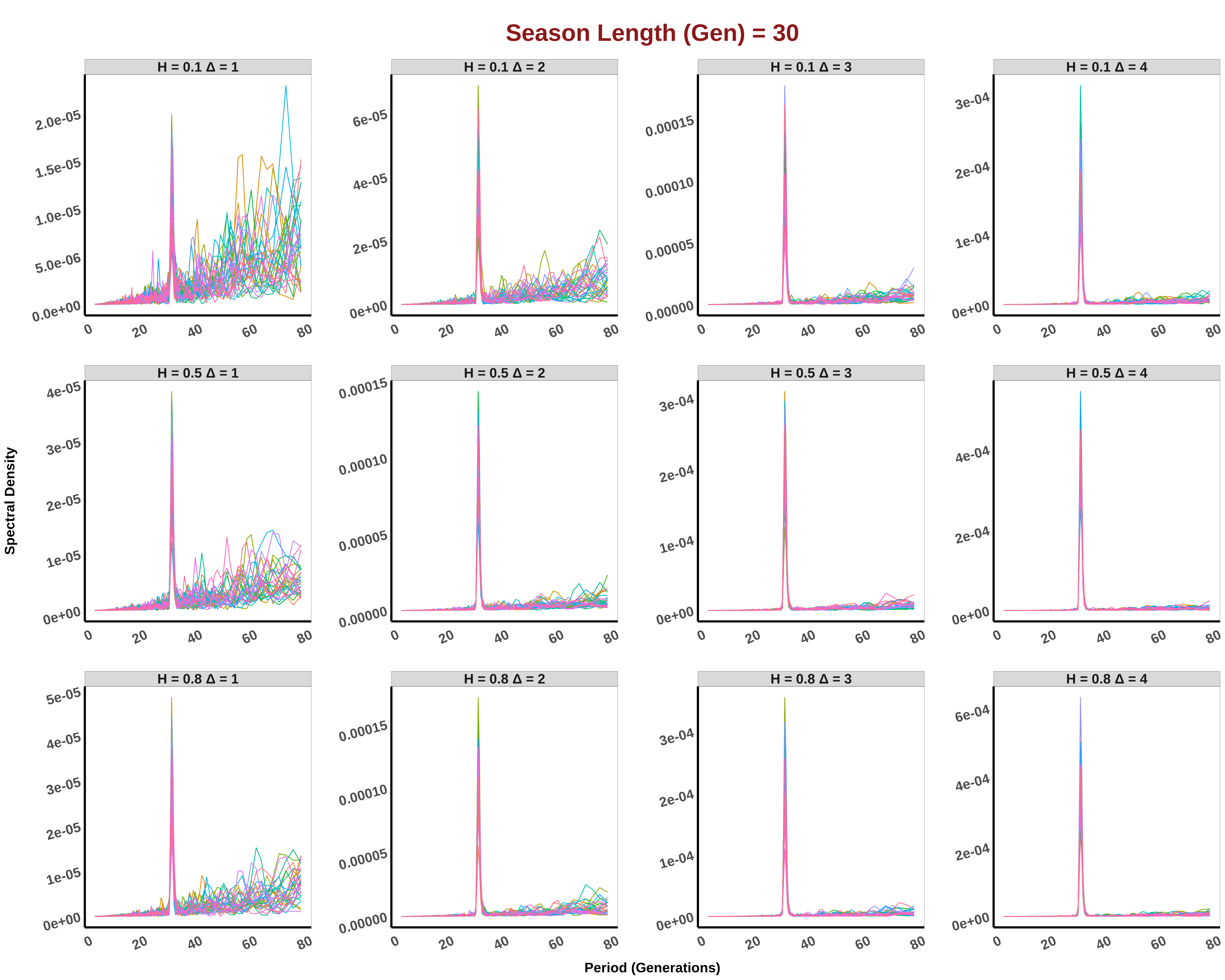

SI 7.2.4.3. Spectral density across 30 replicates for polygenic trait (300 loci) with gradual optima change two equal seasons, all parameter combinations with selection length of 30 generations. Each peak represents a full cycle of selection period.

### 8. Complex model I

#### 8.1. Allele frequency all parameters for a single replicate

##### 8.1.1. Monogenic

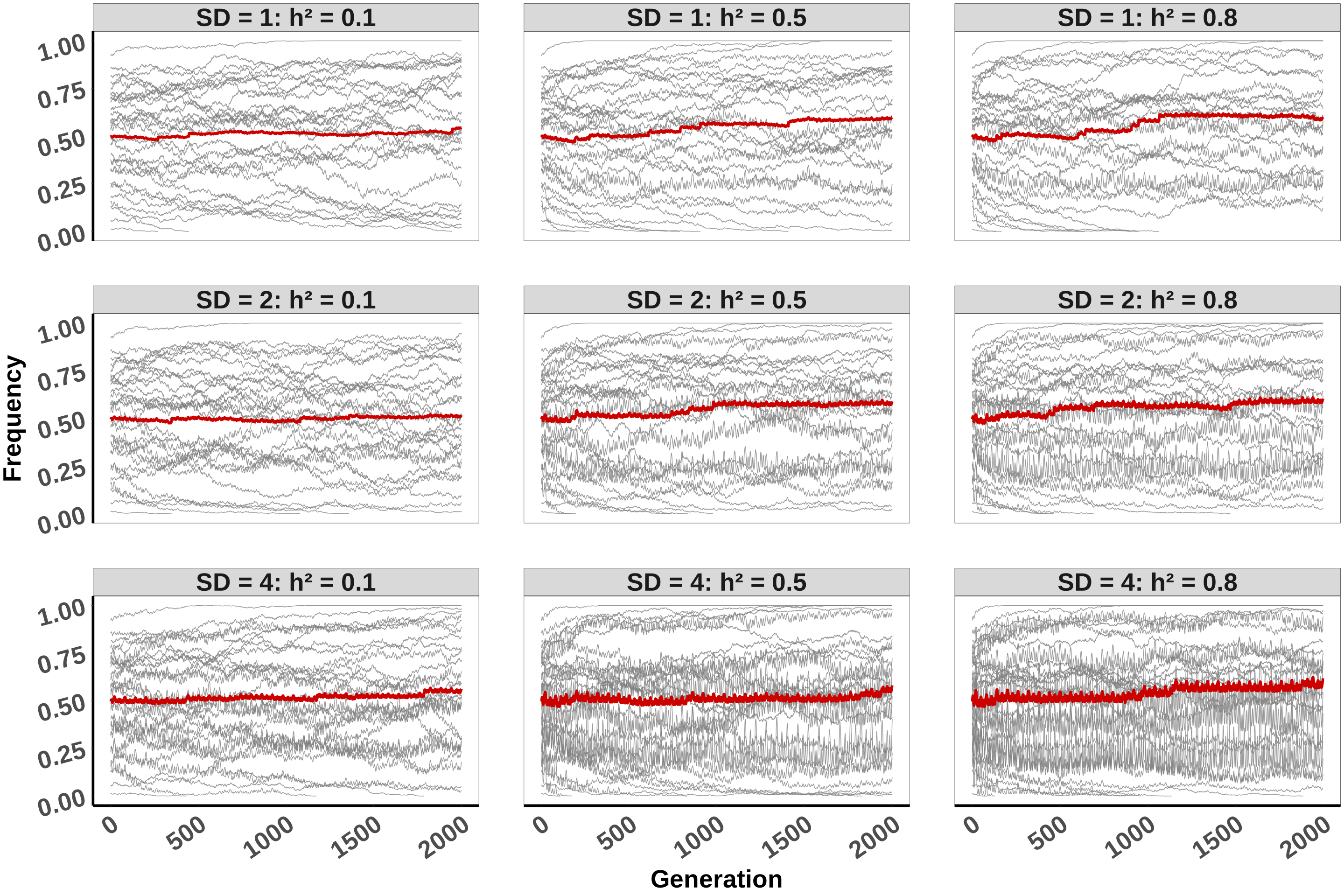

SI 8.1.1. Single locus 30 replicates allele frequency dynamics for gradual optima change with even optima and uneven season length, showing all parameter interactions. The highlighted line indicates the mean allele frequency for each generation.

##### 8.1.2. Oligogenic

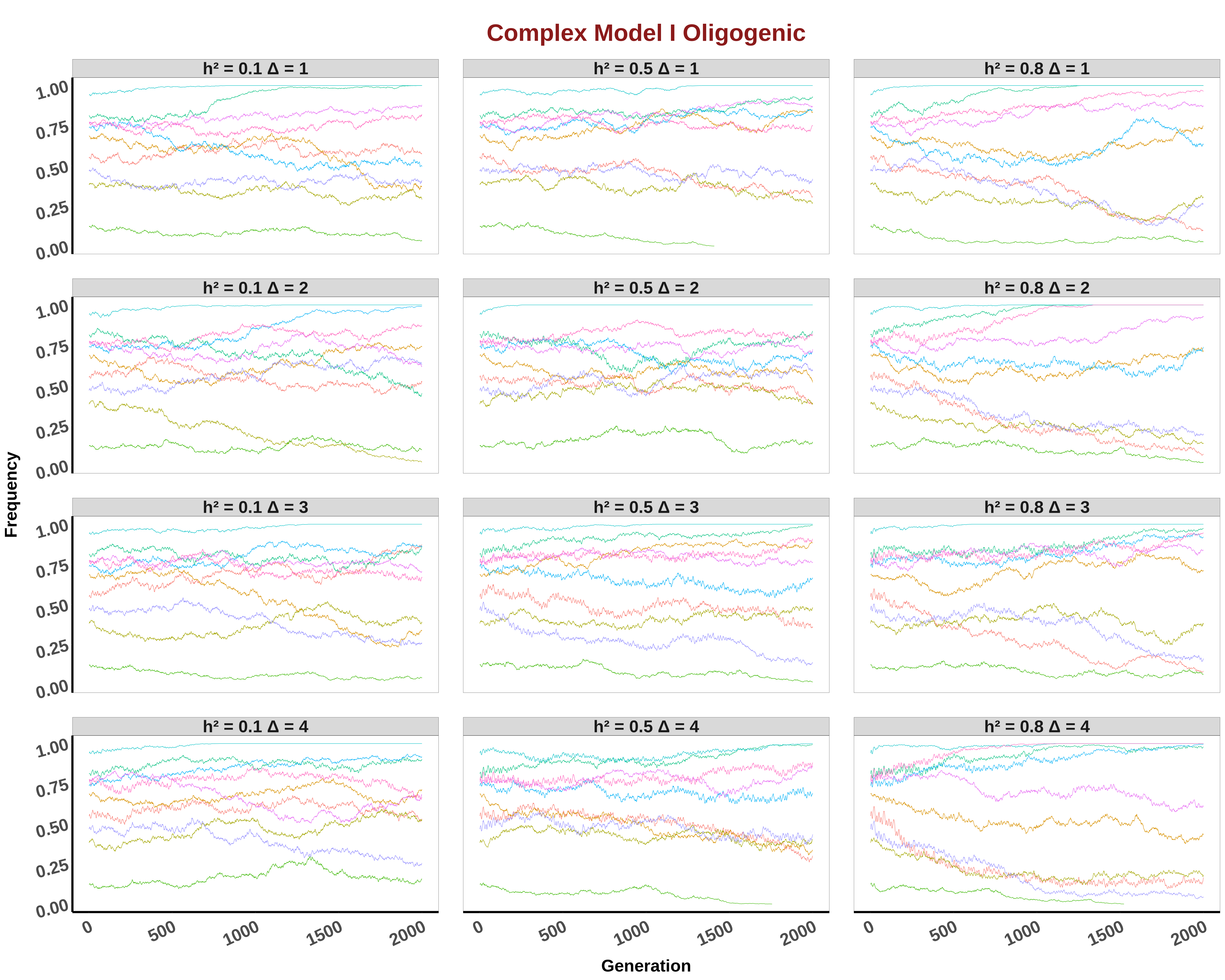

SI 8.1.2. Allele frequency dynamics for all parameter combinations from a single replicate, gradual optima change with even optima and uneven season length. Each line represents the frequency trajectory for a locus from oligogenic trait (10 loci).

##### 8.1.3. Polygenic 70 loci

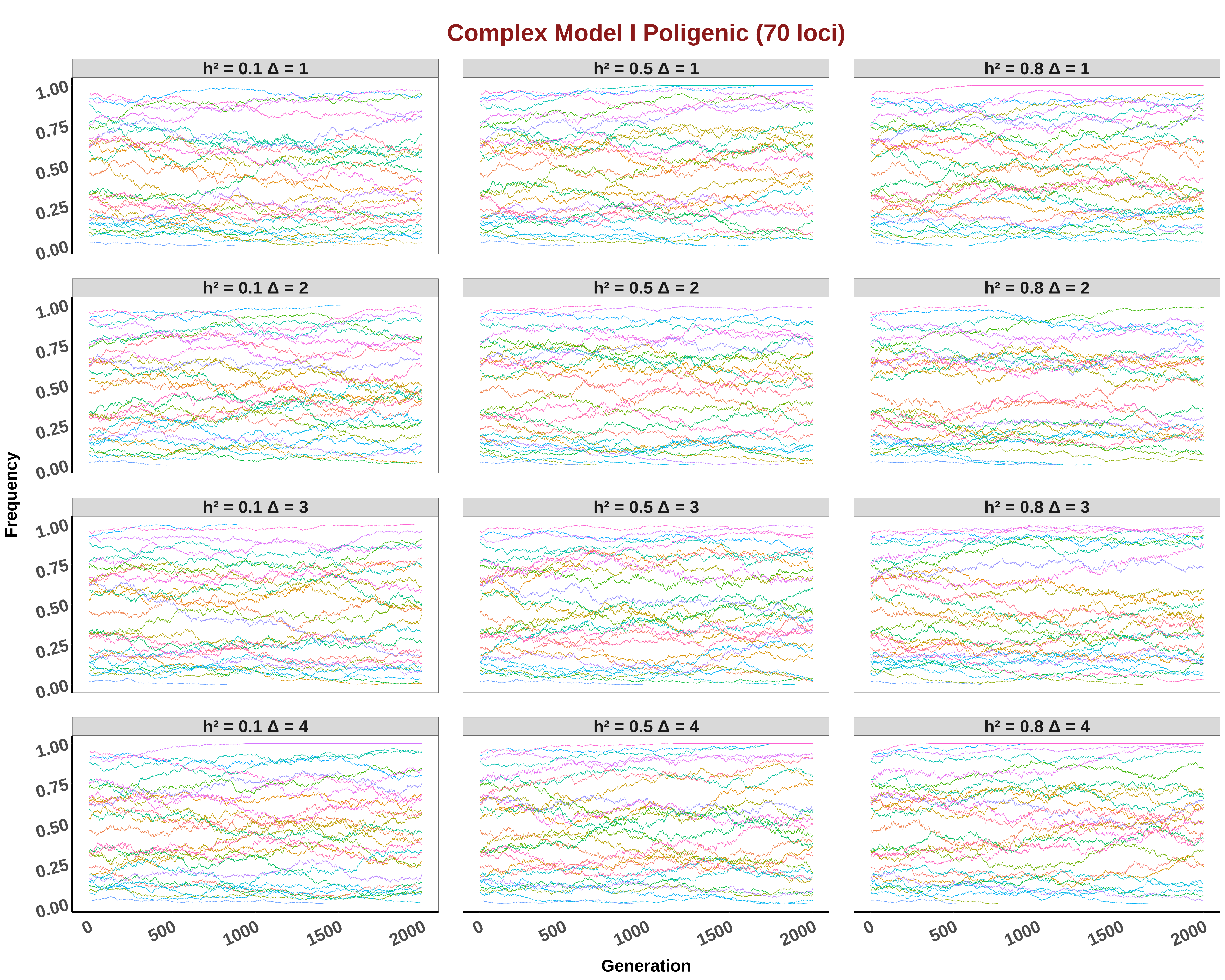

SI 8.1.3. Allele frequency dynamics for all parameter combinations from a single replicate, gradual optima change with even optima and uneven season length. Each line represents the frequency trajectory for a locus from polygenic trait (70 loci).

##### 8.1.4. Polygenic 300 loci

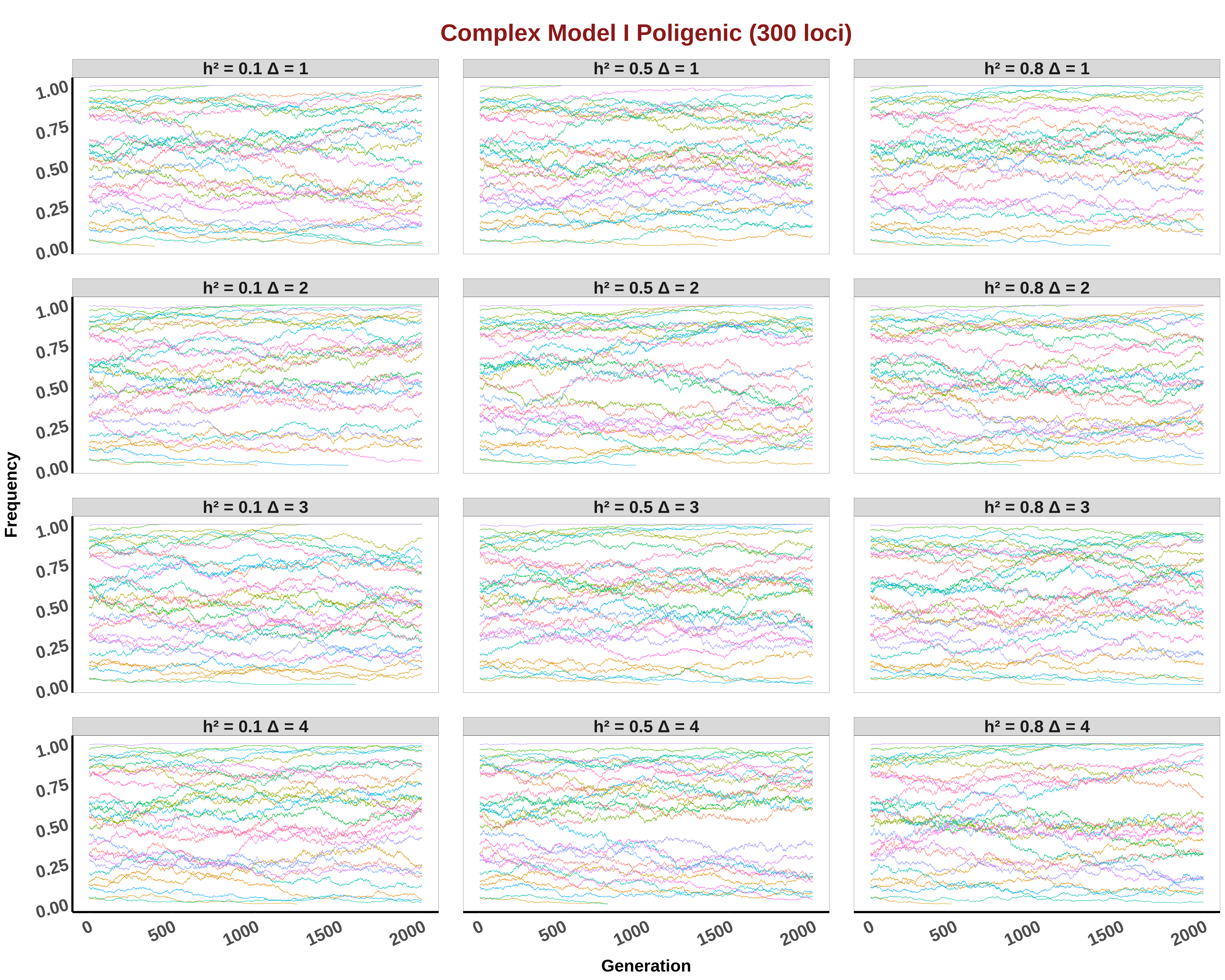

SI 8.1.4. Allele frequency dynamics for all parameter combinations from a single replicate, gradual optima change with even optima and uneven season length. Each line represents the frequency trajectory for a locus from polygenic trait (300 loci).

#### 8.2. Spectral analysis for all parameters

##### 8.2.1. Polygenic 70

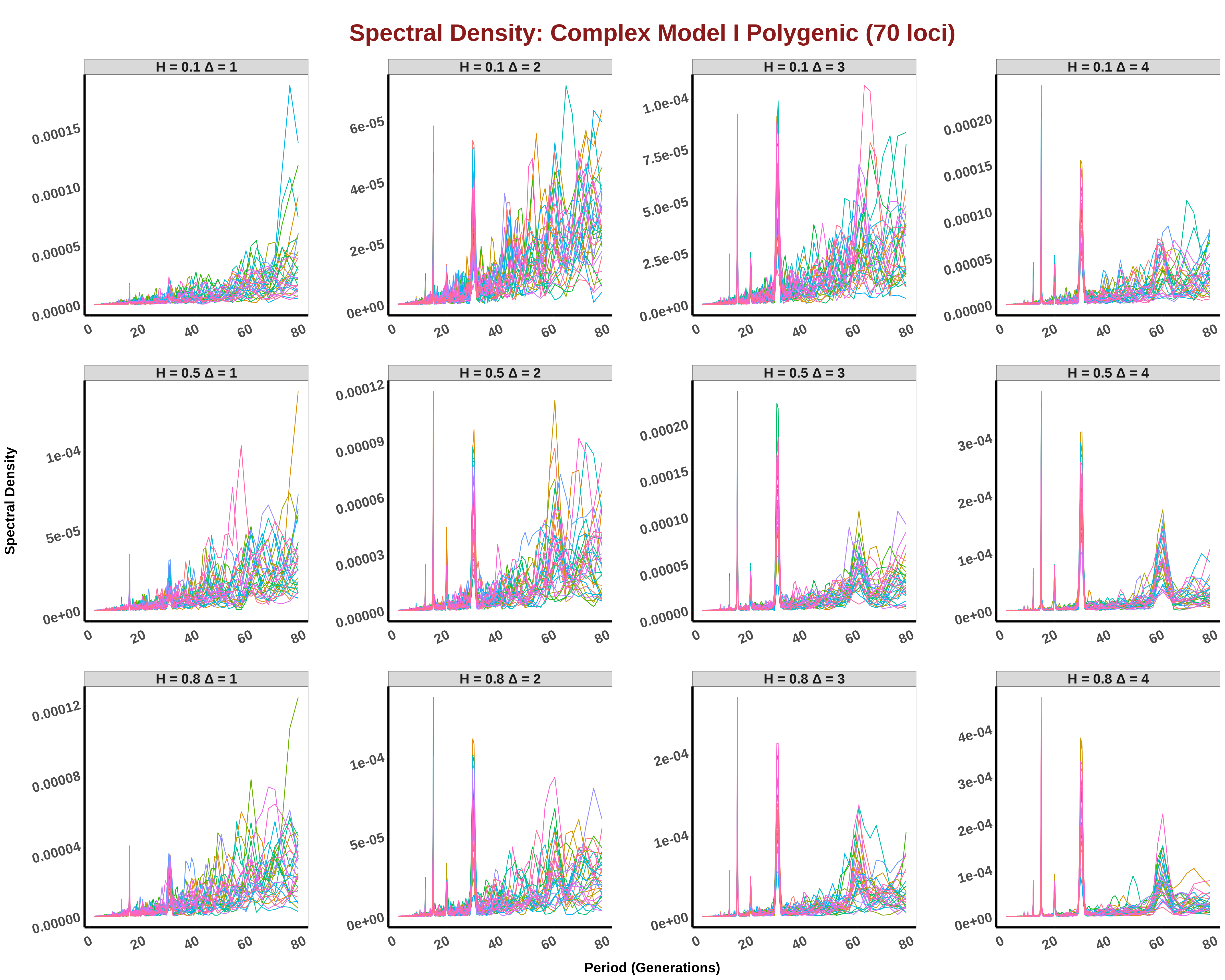

SI 8.2.1. Spectral density across 30 replicates for polygenic trait (70 loci) with gradual optima change with even optima and uneven season length, all parameter combinations. For high parameters, Each peak represents one of the four full cycle of selection periods while the peaks are murky under low heritability and selection pressure (first column).

##### 8.2.2. Polygenic 300

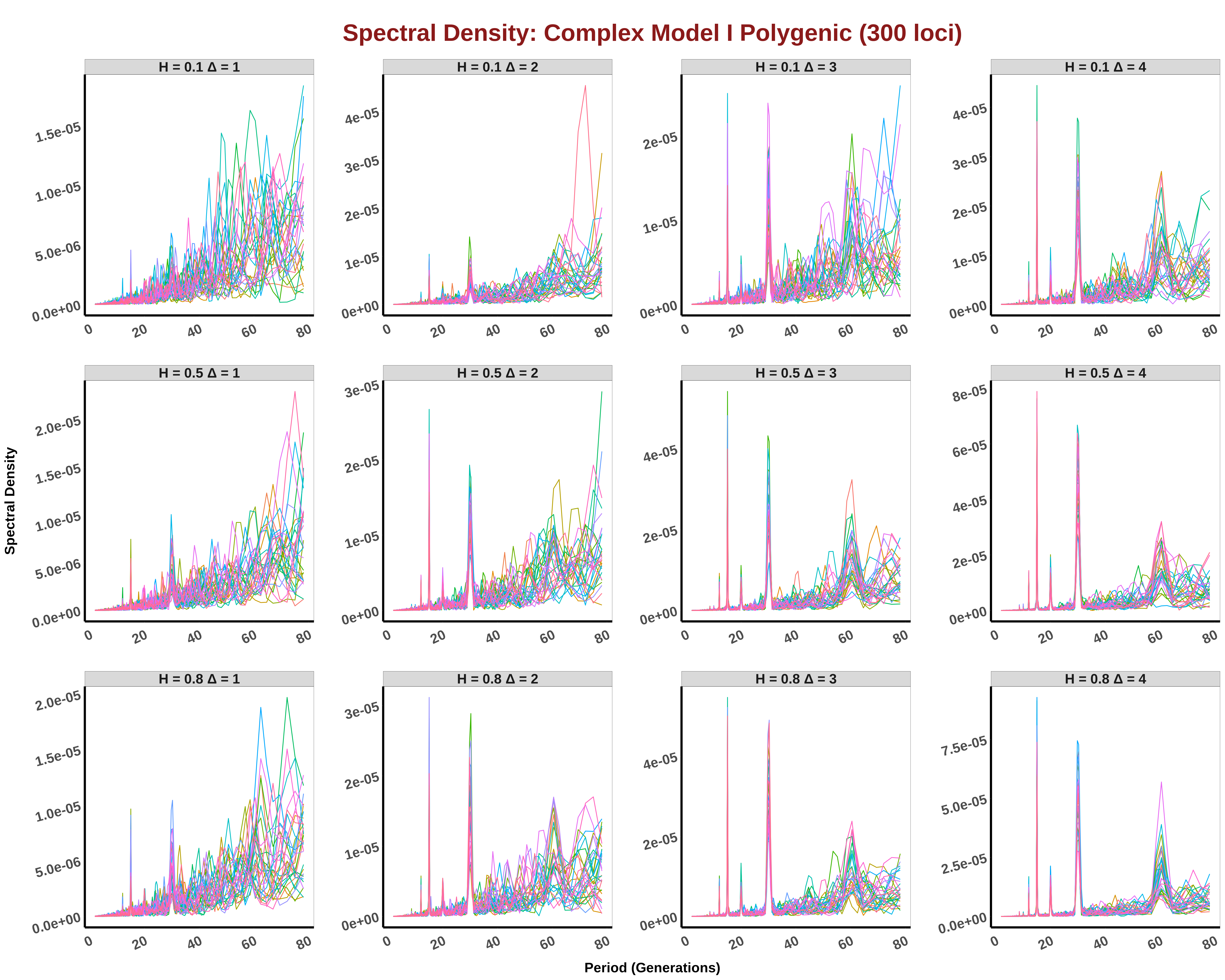

SI 8.2.2. Spectral density across 30 replicates for polygenic trait (300 loci) with gradual optima change with even optima and uneven season length, all parameter combinations. For high parameters, Each peak represents one of the four full cycle of selection periods while the peaks are murky under low heritability and selection pressure (first column).

### 9. Complex model II

#### 9.1. Allele frequency all parameters for a single replicate

##### 9.1.1. Monogenic

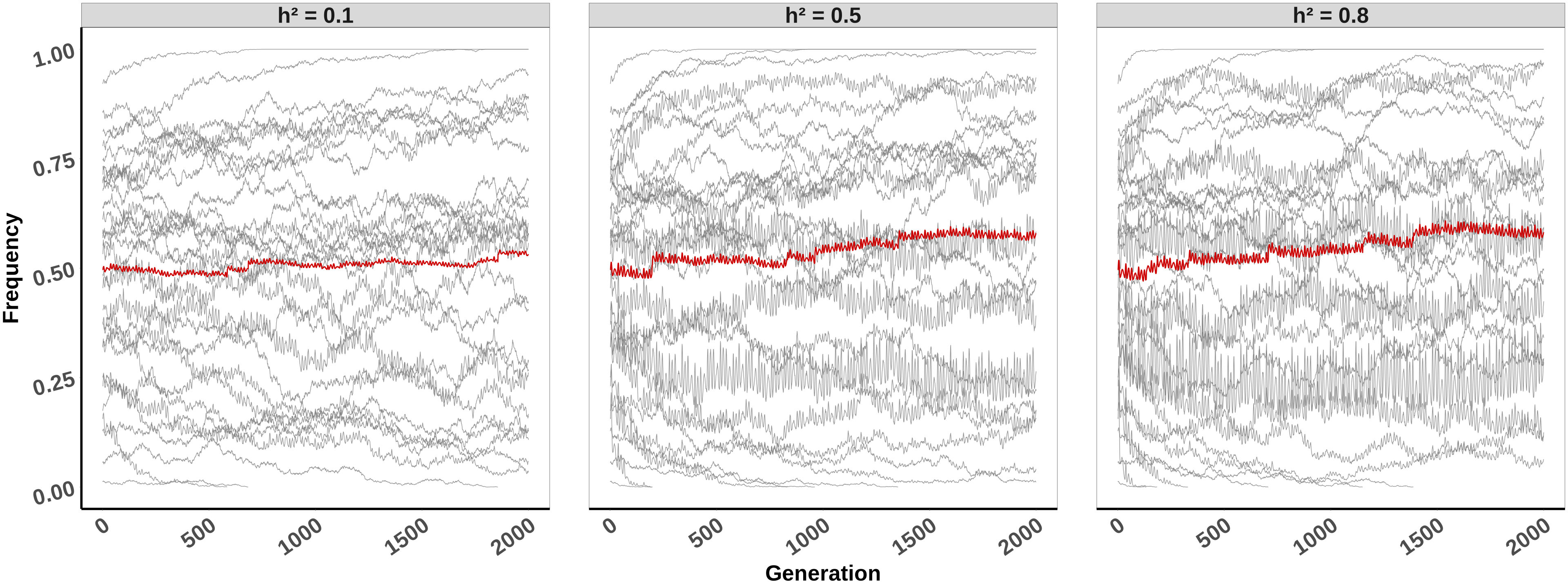

SI 9.1.1. Single locus 30 replicates allele frequency dynamics for gradual optima change with uneven optima and uneven season length, showing all parameter interactions. The highlighted line indicates the mean allele frequency for each generation.

##### 9.1.2. Oligogenic

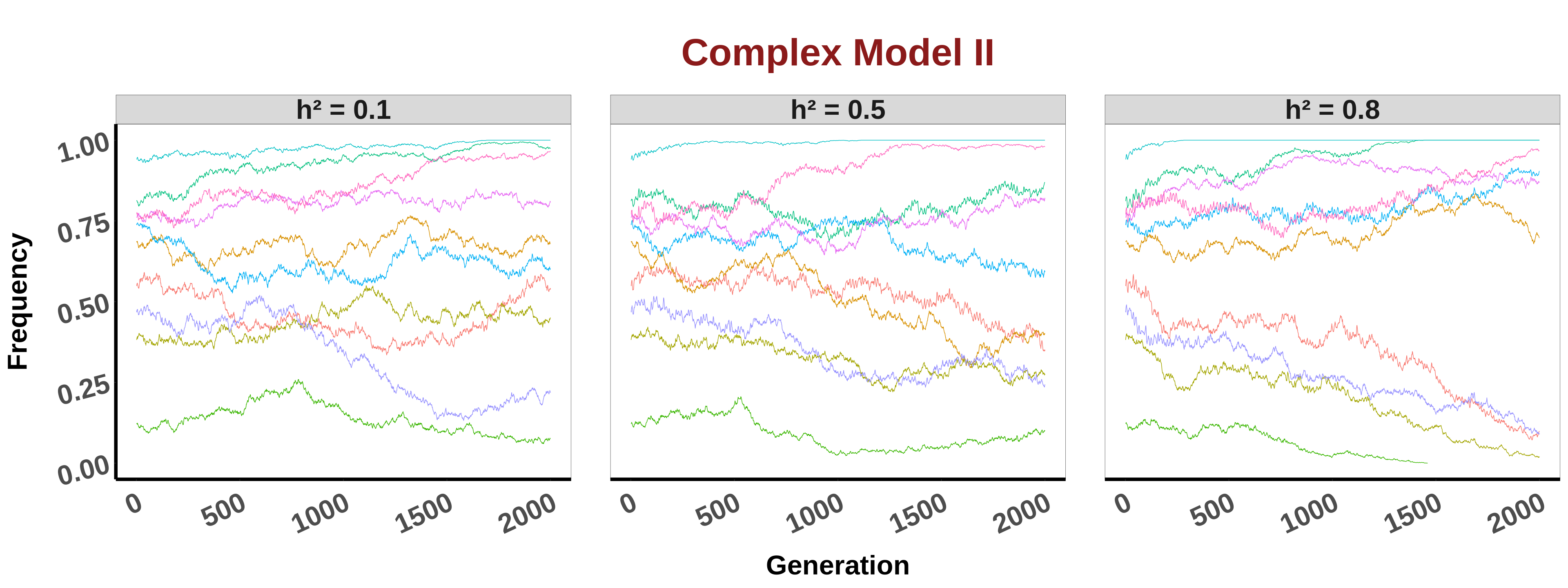

SI 4.1.2. Allele frequency dynamics for all parameter combinations from a single replicate, gradual optima change with uneven optima and uneven season length. Each line represents the frequency trajectory for a locus from oligogenic trait (10 loci).

##### 9.1.3. Polygenic 70 loci

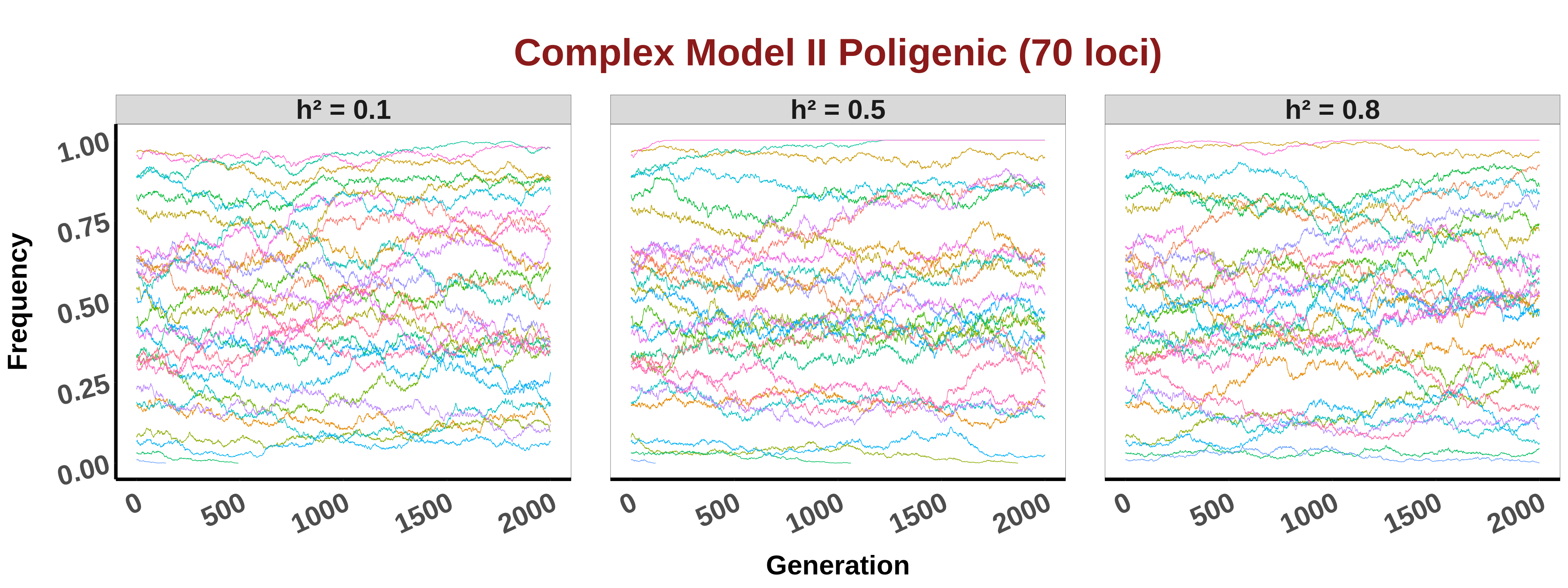

SI 4.1.3. Allele frequency dynamics for all parameter combinations from a single replicate, gradual optima change with uneven optima and uneven season length. Each line represents the frequency trajectory for a locus from polygenic trait (70 loci).

##### 9.1.4. Polygenic 300 loci

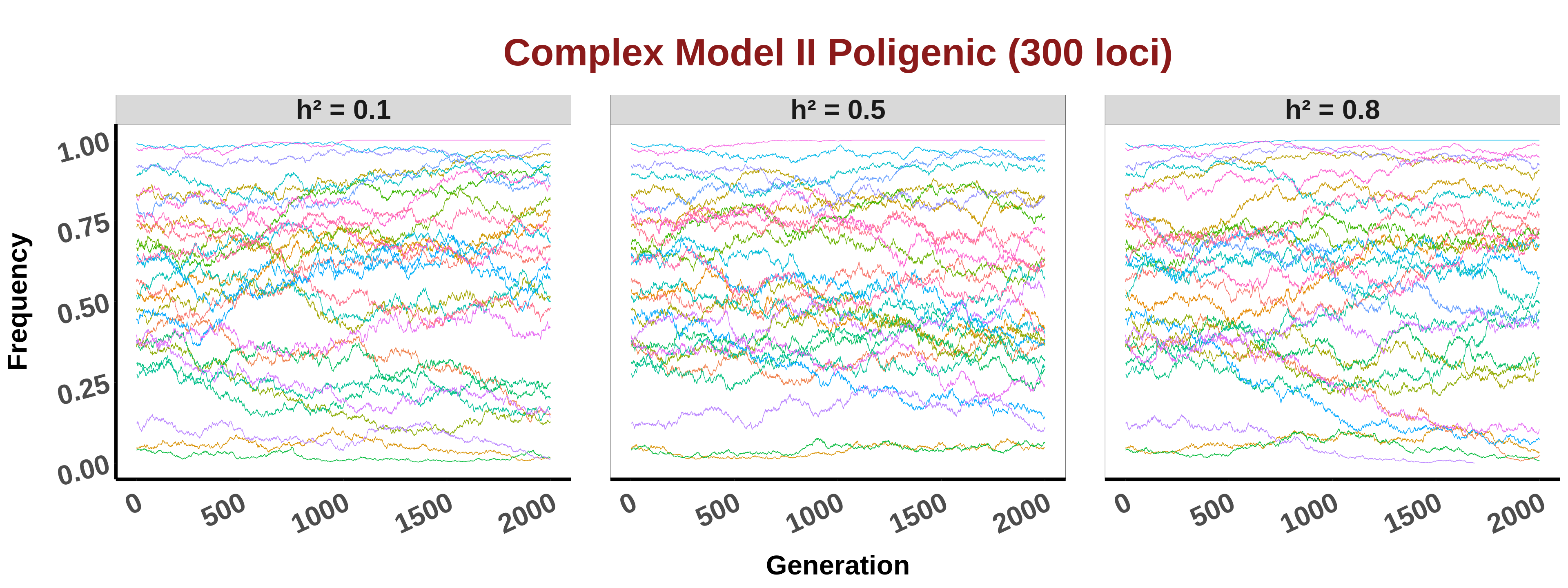

SI 9.1.4. Allele frequency dynamics for all parameter combinations from a single replicate, gradual optima change with uneven optima and uneven season length. Each line represents the frequency trajectory for a locus from polygenic trait (300 loci).

#### 9.2. Spectral analysis for all parameters

##### 9.2.1. Polygenic 70

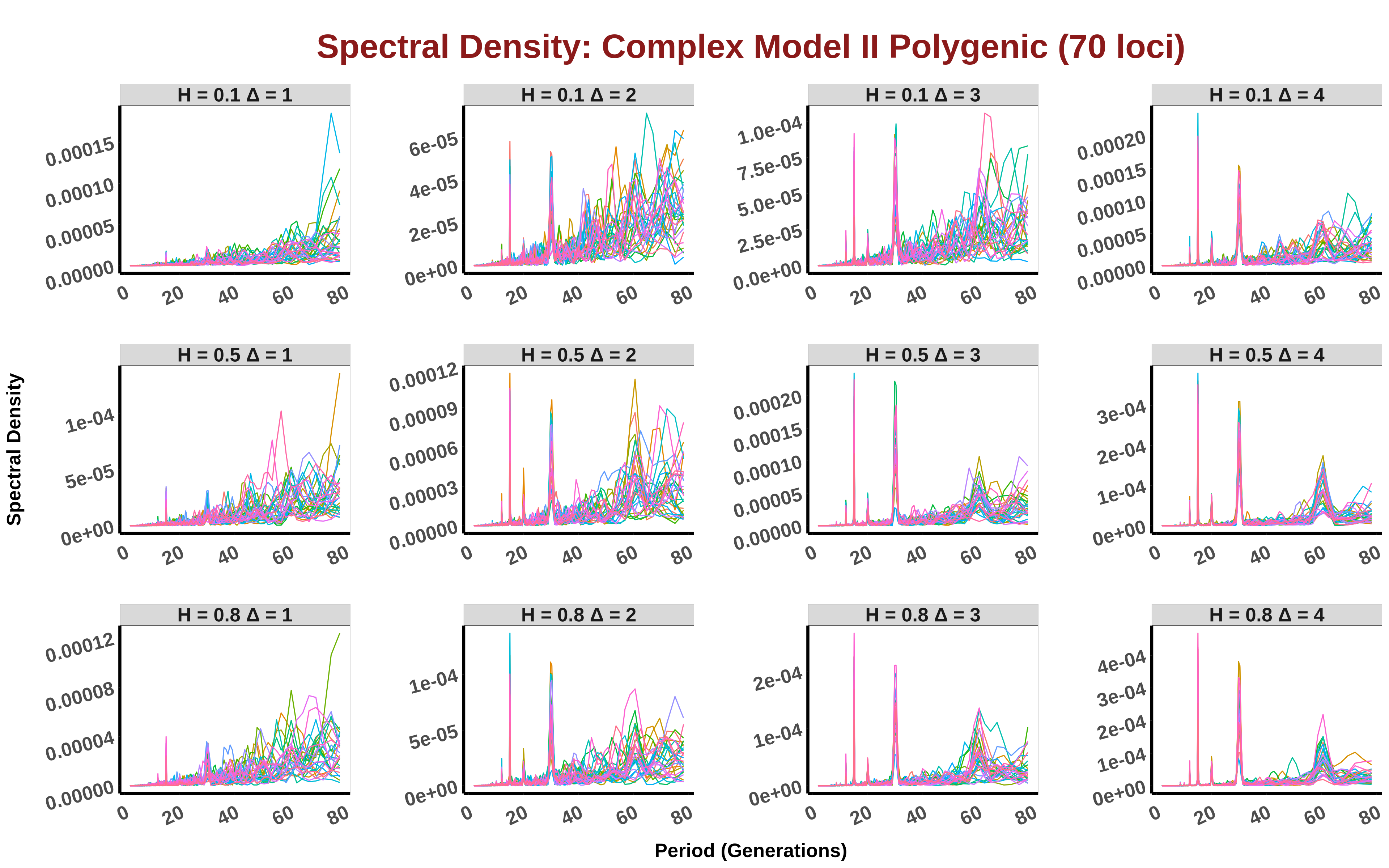

SI 9.2.1. Spectral density across 30 replicates for polygenic trait (70 loci) with gradual optima change with uneven optima and uneven season length, all parameter combinations. For high parameters, Each peak represents one of the four full cycle of selection periods while the peaks are murky under low heritability and selection pressure (first column).

##### 9.2.2. Polygenic 300

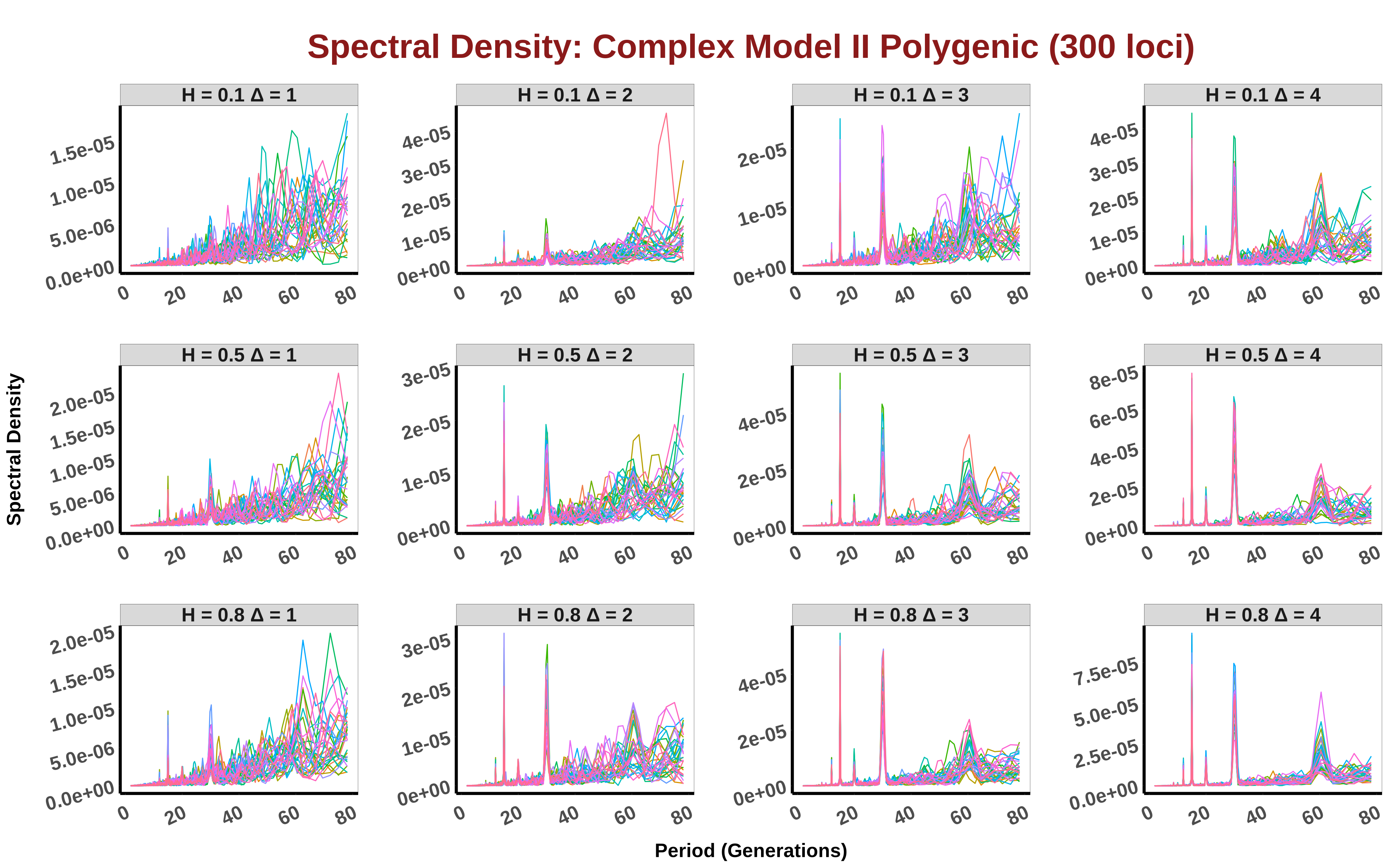

SI 9.2.2. Spectral density across 30 replicates for polygenic trait (300 loci) with gradual optima change with uneven optima and uneven season length, all parameter combinations. For high parameters, Each peak represents one of the four full cycle of selection periods while the peaks are murky under low heritability and selection pressure (first column).

#### 10. Spectral All models comparisons

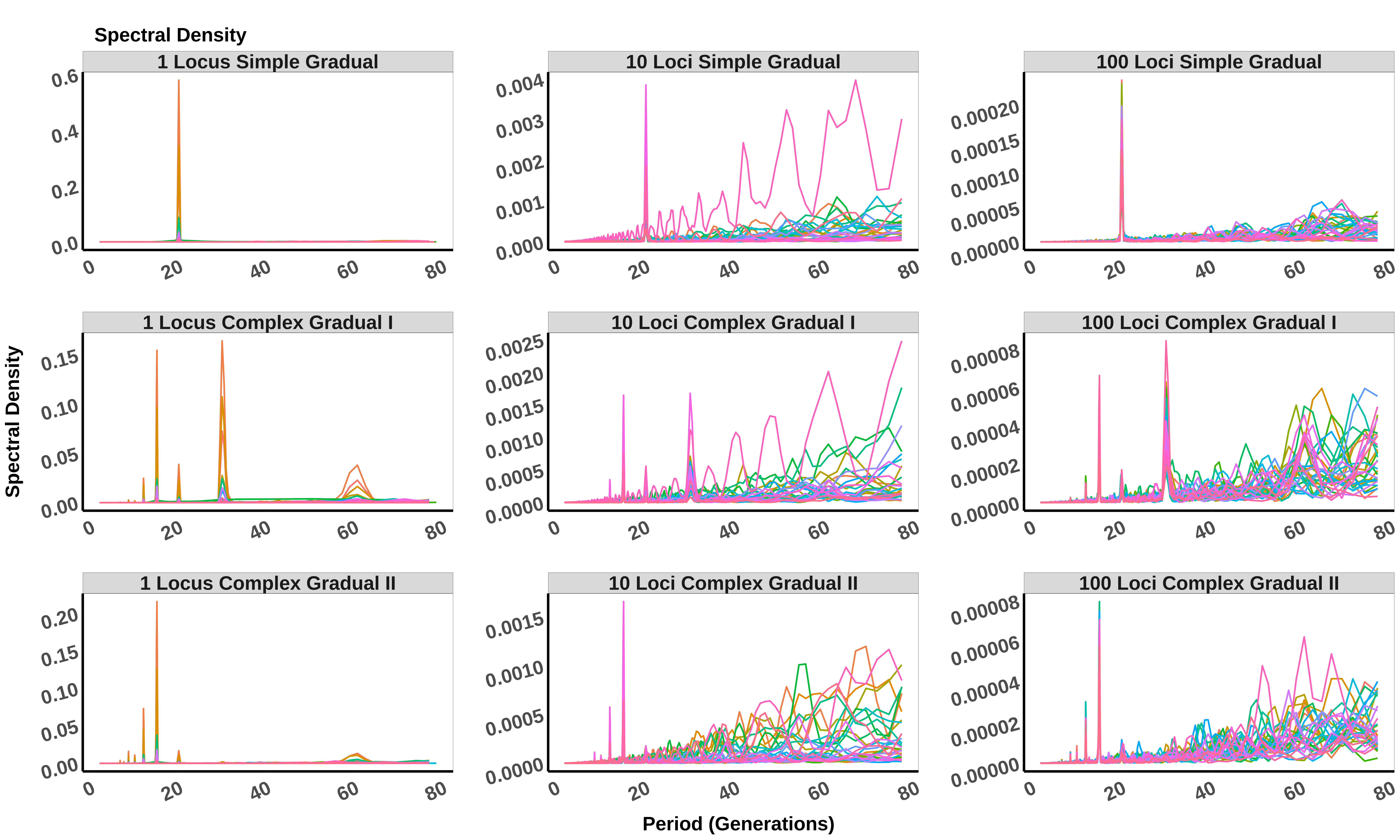

SI 10. Spectral analysis across 30 replicates for monogenic(left), oligogenic(middle), and polygenic(right) traits, with each colored is one replicate. For the simple gradual model, the parameters are h2 = 0.5, Δ = 2, and Gen = 20. Thus, the dominant spectral peaks occur at 40, consistent with a full seasonal cycle. Although the oligogenic model has more noise, the complex gradual model I (uneven season lengths and even distance to optima Δ) exhibit four distinct peaks aligned with the four-season lengths, plus a faded long period peak at 60 which signal the recurrence across multiple cycles. In complex gradual II (uneven season lengths and uneven Δ), the spectrum is noisier than other selection regimes, and it is challenging to observe all peaks corresponding to the number of considered seasons mainly because some seasons have a small selection pressure and the dynamics in those seasons are masked by the presence of seasons experiencing stronger selections.

#### 11. Fluctuating population fixation

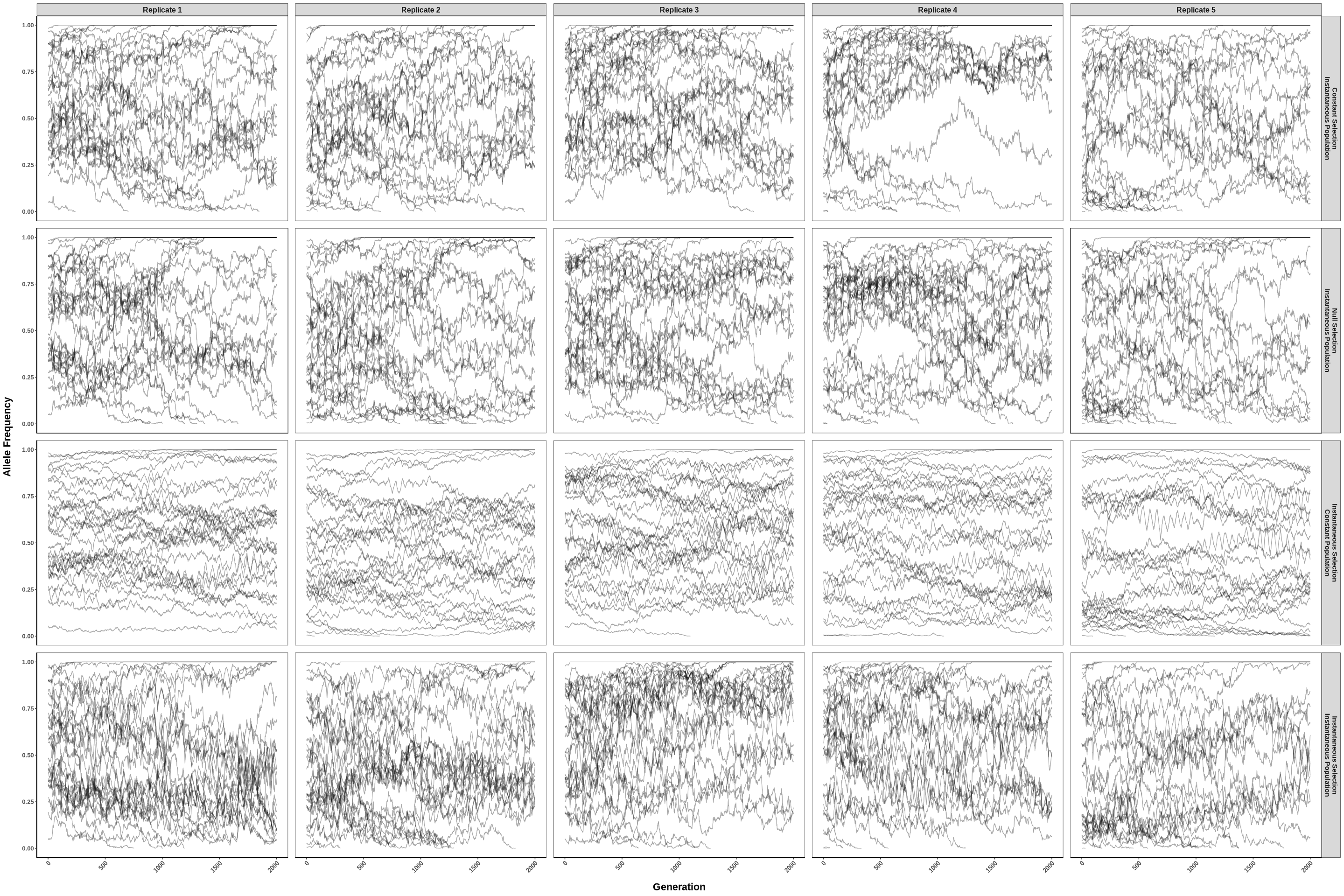

SI 11. Allele-frequency dynamics and initial QTL fixation across models with fluctuating selection and/or fluctuating population size for a polygenic model of 100 loci. The top panel shows allele-frequency dynamics for 30 randomly selected initial QTL alleles across five replicates in the high-parameter regime, while the bottom panel shows the mean ± SE percentage of initial QTL alleles fixed across 30 replicates.

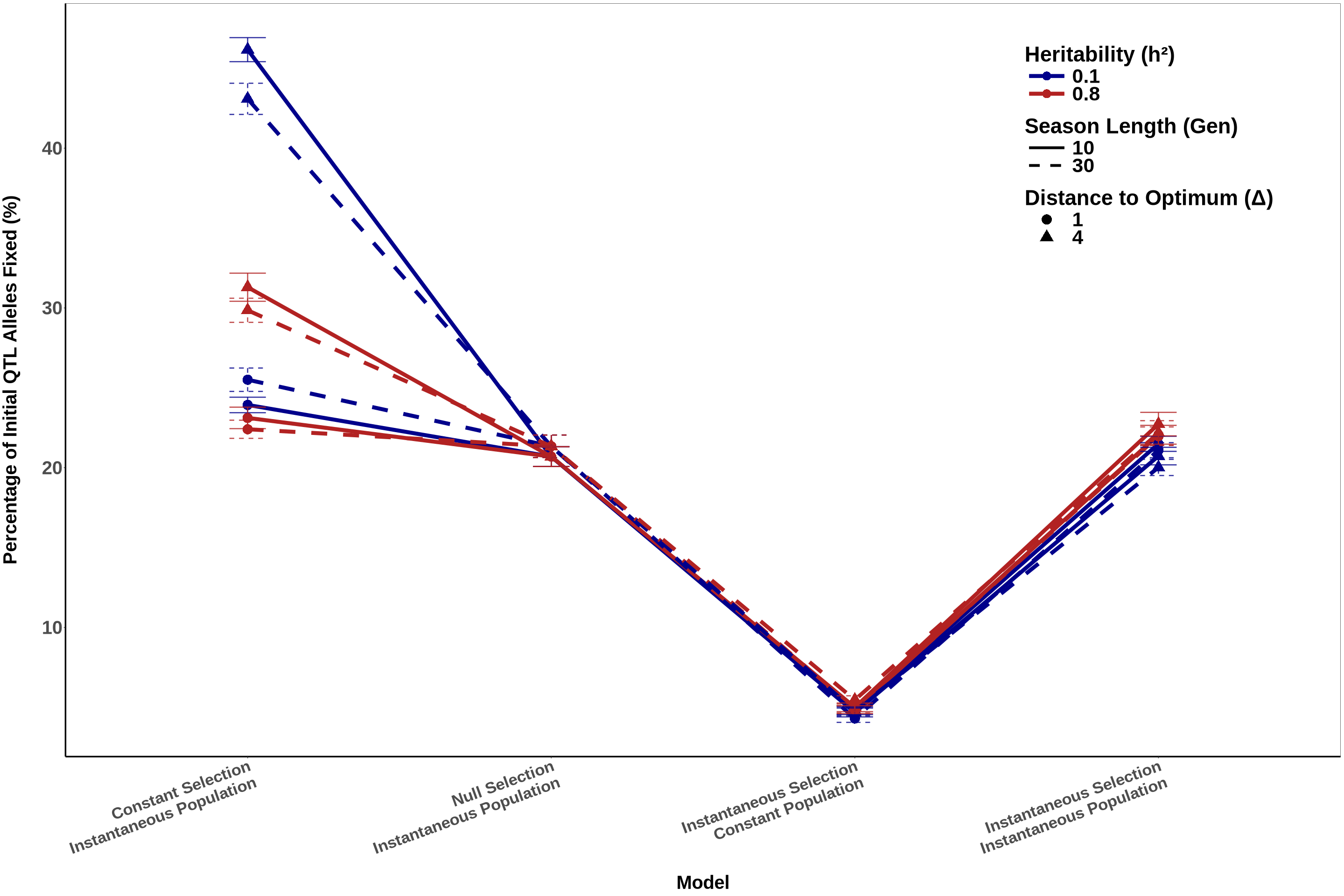

SI 11. Allele-frequency dynamics and initial QTL fixation across models with fluctuating selection and/or fluctuating population size for a polygenic model of 100 loci. The top panel shows allele-frequency dynamics for 30 randomly selected initial QTL alleles across five replicates in the high-parameter regime, while the bottom panel shows the mean ± SE percentage of initial QTL alleles fixed across 30 replicates.
